# Tissue Transglutaminase 2 has higher affinity for relaxed than for stretched fibronectin fibers

**DOI:** 10.1101/2023.08.14.553221

**Authors:** Kateryna Selcuk, Alexander Leitner, Lukas Braun, Fanny Le Blanc, Paulina Pacak, Simon Pot, Viola Vogel

## Abstract

Tissue transglutaminase 2 (TG2) plays a vital role in stabilizing extracellular matrix (ECM) proteins through enzymatic crosslinking during tissue growth, repair, and inflammation. TG2 also binds non-covalently to fibronectin (FN), an essential component of the ECM, facilitating cell adhesion, migration, proliferation, and survival. However, the interaction between TG2 and fibrillar FN remains poorly understood, as most studies have focused on soluble or surface-adsorbed FN or FN fragments, which differ in their conformations from insoluble FN fibers. Using a well-established *in vitro* FN-fiber stretch assay, we discovered that the binding of a crosslinking enzyme to ECM fibers is mechano-regulated. TG2 binding to FN is tuned by the mechanical tension of FN fibers, whereby TG2 predominantly co-localizes to low-tension FN fibers, while fiber stretching reduces their affinity for TG2. This mechano-regulated binding relies on the proximity between the N-terminal β-sandwich and C-terminal β-barrels of TG2. Crosslinking mass spectrometry (XL-MS) revealed a novel TG2-FN synergy site within TG2’s C-terminal β-barrels that interacts with FN regions outside of the canonical gelatin binding domain, specifically FNI_2_ and FNIII_14-15_. Combining XL-MS distance restraints with molecular docking reveals the mechano-regulated binding mechanism between TG2 and modules FNI_7-9_ by which mechanical forces regulate TG2-FN interactions. This highlights a previously unrecognized role of TG2 as a tension sensor for FN fibers. This novel interaction mechanism has significant implications in physiology and mechanobiology, including how force regulate ECM deposition and maturation, and how TG2 mediates cell-ECM adhesion in health and in various pathophysiological processes. Data are available via ProteomeXchange with identifier PXD043976.

## Introduction

The growing field of mechanobiology has revealed that not only the biochemical, but also the physical properties of the extracellular matrix (ECM) have a major impact on cell decision making in development, hemostasis and wound healing^1–4^, and when altered can drive pathological transformations, including progressive cancer and fibrotic pathologies^5^. Enzymatic crosslinking of ECM fibers is necessary for the mechanical stabilization during tissue growth and repair, but also plays a major role as the driver of fibrotic diseases and malignancy^6–8^. Transglutaminase 2 (TG2), also referred to as tissue transglutaminase 2, is mostly retained within the cell under homeostatic conditions, but upon tissue injury or inflammation, its expression and subsequent export to the cell surface and the extracellular environment are strongly upregulated^9–12^. Once secreted, TG2 alters the physico-chemical properties of the extracellular environment by enzymatically crosslinking various ECM proteins, making ECM stiffer and more resistant to proteolytic degradation. In turn, this triggers various downstream effects which change cell behavior promoting cell adhesion, migration and fibroblast proliferation^11,13^. While these processes are essential to stabilize the provisional ECM during wound healing, they need to be tightly regulated, and aberrant TG2 activity leads to pathological fibrosis^6,14,15^. Indeed, it was shown in healthy tissues that most externalized TG2 is catalytically inactive and only transiently activated by stress signals^16,17^. However, the crosslinking activity is only one component of the large functional arsenal of TG2. Numerous studies have shown that it also acts as a non-enzymatic scaffold protein that interacts with many ECM components and cell-surface receptors to support cell adhesion, migration, proliferation and survival, such as fibronectin, collagen, vitronectin, integrins, syndecan-4, several growth factor receptors and others^18,19^.

Fibronectin (FN) is one of the best characterized binding partners of TG2 in the ECM, which is over-expressed during development, tissue growth and repair as well as under various pathological conditions. FN polymerizes to form fibrous matrices that promote cell adhesion, migration, and proliferation. Usually, cells adhere to FN through transmembrane receptors called integrins through the Arg-Gly-Asp (RGD) integrin binding site on FNIII_10_. However, RGD-dependent cell adhesion and the associated outside-in integrin signaling may be disrupted during extensive tissue damage, matrix remodeling and ECM degradation. Blocking of the cell adhesion with synthetic RGD-peptides causes reduction in FN-integrin interaction and in the absence of TG2 leads to detachment-induced apoptosis (anoikis) in many cell types^14,19,20^. However, when TG2 is expressed on the cell surface, it can rescue cells from anoikis when RGD-dependent adhesion is blocked, thereby promoting cell survival^14,20–22^. This rescue does not depend on the enzymatic activity of TG2, but requires FN-binding together with the assistance from heparan sulfate chains of Syndecan-4 and/or non-canonical binding to β1-integrins^14,20,23^. Furthermore, TG2-FN interaction enhances deposition of FN fibers with assistance from Syndecan-4 and β1-integrins, when RGD-dependent adhesion is attenuated, thus helping to quickly restore the extracellular environment after injury^22,24^.

Given the pro-survival adhesive properties of the TG2-FN complex, it is not surprising that high expression levels of TG2 favor metastasis formation in multiple cancers^8,25–30^. Consequently, TG2 upregulation in these tumors is strongly associated with poor patient outcome^31^. This makes the TG2-FN complex a compelling drug target^7,13^. Currently, efforts are underway to develop small molecule inhibitors that disrupt the TG2-FN interface^32–35^. Thus, a deeper mechanistic and structural understanding of the TG2-FN interaction sites would not only shed light on its role in wound healing and cancer, but it would also assist the rational design of drugs.

TG2 consists of four domains: an N-terminal β-sandwich, a catalytic core, and two C-terminal β-barrels (Fig. 1A). While previous structural studies have characterized TG2 in two conformational states (“open” and “closed”) using crystallography^36,37^, evidence suggests that these two states do not adequately capture the protein’s conformational plasticity. When TG2 is bound to GDP, GTP or other purine nucleotides, it adopts an enzymatically inactive “closed” conformation. In this state, the C-terminal β-barrel domains tightly fold over the catalytic domain, obstructing access to the active site (PDB:1KV3)^37^ (Fig. 1B). However, when the active site is covalently bound to an irreversible inhibitor Z-DON, TG2 undergoes a large conformational change (PDB:2Q3Z)^36^. The β-barrel domains are prevented from interacting with the catalytic core, resulting in an “open”, extended conformation (Fig 1B). Although the PDB:2Q3Z conformation is often associated with the catalytically active TG2 due to the readily accessible active site, there are doubts regarding its true representation of the active form^38^. Firstly, PDB:2Q3Z contains a Cys370-Cys371 disulfide bridge near the active site, which inhibits TG2 catalytic activity^39,40^. Secondly, the formation of this disulfide bridge induces local changes in peptide backbone conformation, disrupting the calcium binding sites^40^. Consequently, the PDB:2Q3Z structure, bound to Z-DON inhibitor, does not include any calcium ions, despite their known requirement for the crosslinking function^36^. Finally, no crystal structure of catalytically active TG2 bound to calcium ions has been solved to date. Therefore, it is unlikely that the extended conformation observed in the Z-DON inhibitor-bound TG2 structure accurately represents the calcium-bound catalytically active protein. Indeed, small angle X-ray/neutron scattering (SAXS/SANS), hydrogen-deuterium exchange (HDX) and biosensor measurements suggest that the calcium bound TG2 assumes an “open” conformation distinct from the two known structures^40–42^. The reversible formation of the Cys370-Cys371 disulfide bond acts as a redox switch, inactivating TG2 in the oxidative environment of ECM and desensitizing it to the presence of effectors^39^. In fact, experimental evidence indicates that the Z-DON inhibitor-bound structure resembles the oxidized TG2, i.e. TG2 in the extended effector-free state^40,42^. Thus, TG2 can adopt more than the two crystallographically captured conformations and exists in at least three different states: nucleotide binding favors the “closed” conformation^37^, while high calcium concentrations predominantly induce the “open” but yet unknown conformation^40,42^. Importantly, only this open state is catalytically active. In the absence of calcium or GDP/GTP and under oxidative conditions, TG2 assumes a catalytically inactive “open” effector-free state^40^ (Fig. 1B).

**Figure 1:**
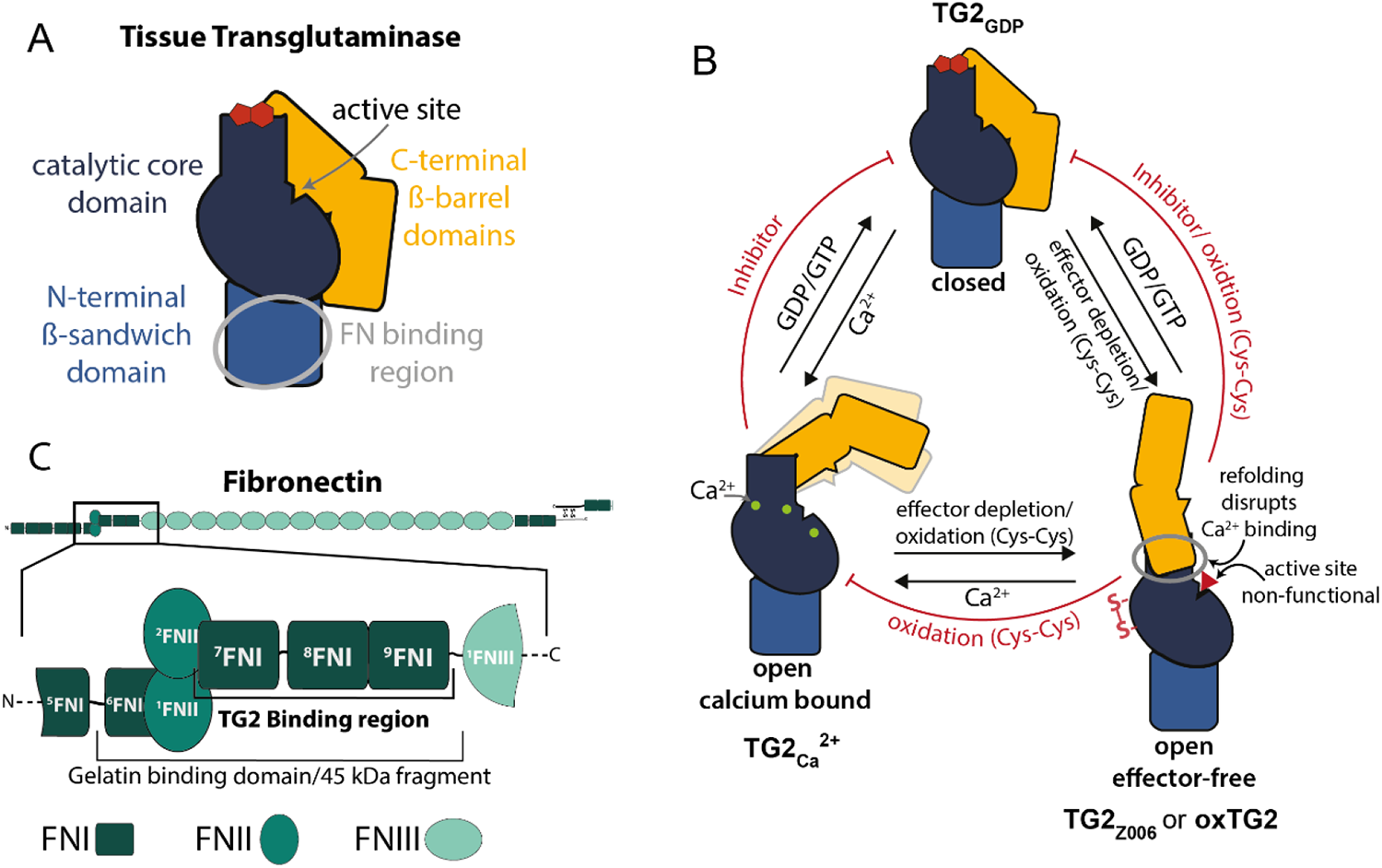
Schematic illustration of TG2’s domain composition, its conformational states and known binding interactions with fibronectin (FN). **A:** Schematic view of TG2’s domain architecture based on the crystal structure of the catalytically inactive, GDP bound state (PDB:1KV3)^37^. The FN binding region on the N-terminal domain determined by Cardoso et al. is highlighted^45^. **B:** TG2 can exist in an equilibrium of at least three distinct conformational states: closed – TG2_GDP_, open calcium bound – TG2_Ca_^2+^ and open effector-free state. The relative population of each state depends on the concentration of its allosteric effectors – calcium (green circles) – TG2_Ca_^2+^ and GDP/GTP (red polygons) – TG2_GDP_ and can be further regulated by the formation of intramolecular disulfide bridges (oxidation – oxTG2) or the binding of artificial irreversible inhibitor Z-DON (red triangle) to the active site Cys277 (TG2_Z006_). **C:** Illustration of the domain architecture of FN. Only one chain of the disulfide-linked FN homodimer is fully shown (top). The TG2 binding region in the gelatin binding domain determined by Soluri et al. (FN type I modules 7-9) is highlighted in the zoom-in^43^.

While significant progress has been made towards mapping the interactions between TG2 and FN more precisely, previous studies that sought to investigate TG2-FN interactions were either performed with soluble dimeric FN or with shorter FN fragments^21,43–48^. Thus, nothing is known about how FN fibrillogenesis might affect its interactions with TG2. Soluble FN, as well as many of its fragments adopts a quaternary structure, which is distinct from the insoluble fibrillar form in the ECM^49–51^. Although the exact structure of the fibrillar FN remains unknown, mutagenesis and then a super-resolution microscopy study has shown that that FN fibrillogenesis requires the N-terminal FnI_1-5_ domains^52^ and that FN polymerizes in a antiparallel fashion with an N-terminal overlap of almost 40 nm^53^. Cell-mediated stretching of FN fibers during FN fibrillogenesis can either create additional interaction sites for its binding partners, or structurally perturb others^54^. Many FN domains were shown to act as mechano-chemical switches: when FN fibers are mechanically stretched or relaxed, this can either destroy or open up binding epitopes, thereby altering the protein’s biochemical functions and changing down-stream outside-in cell signaling^54–62^. Intriguingly, we have previously demonstrated that the gelatin binding domain (GBD) of FN, located on the FNI_6_FNII_1-2_FNI_7-9_ region, which also overlaps with the main binding site of TG2 on FN (Fig. 1C), acts as such mechanochemical switch in FN interactions with collagen I^60^. Thus, we asked here whether changes in the tensional state of FN-fibers and their force-induced mechanical unfolding might also have an impact, or possibly regulate the binding of TG2 to FN.

To investigate whether FN-fiber tension regulates the affinity for TG2, we employed an *in vitro* FN-fiber stretch assay, which is not only a more physiologically relevant approach, but also enables control over the mechanical strain of FN-fibers. We found that TG2 preferentially binds to FN-fibers under low strain, and that mechanical stretching of FN fibers reduces TG2 affinity for them. Furthermore, we demonstrate that the mechano-regulated interaction of TG2 with FN is dependent on the spatial proximity of C-terminal β-barrel domains with the N-terminal β-sandwich of TG2. TG2 mutants lacking β-barrel domains or the wild type TG2 in the open conformation did not exhibit mechano-regulated binding to FN, indicating the presence of a secondary interaction site on TG2 C-terminal β-barrel. To investigate this possibility, we chemically crosslinked TG2 with full-length FN and with the gelatin binding domain and analyzed resulting cross-linked peptides using crosslinking mass spectrometry (XL-MS)^63^. Our XL-MS results unequivocally established that the C-terminal β-barrels of TG2 interacted with regions of FN outside of the canonical gelatin binding domain when TG2 was in the closed conformation and in complex with full-length FN in solution, specifically with the domains FNI_2_ and FNIII_14-15_. Using experimental crosslinks as distance restraints, we integrated these data to guide the structural modeling and docking of TG2 with FN using DisVis/HADDOCK^64,65^. Predicted models of TG2-FN complexes suggest that when N-terminal and C-terminal domains of TG2 were in proximity, the FNI_7-9_ subunits aligned with both N-terminal domain and C-terminal β-barrels of TG2, supporting the hypothesis of multivalent synergy binding sites. To validate the predicted model, we show that the collagen-mimicking peptide R1R2 with the known binding motif^66,67^ on FNI_8-9_ competes with TG2 in a dose dependent manner for binding to FN-fibers under low and high strain. In summary, this study provides mechanistic insights into the interactions between TG2 and both fibrillar and soluble FN. Such advanced knowledge about the mechano-regulation of the binding interface and suggests a novel model how fiber stretching tunes the interaction of TG2 with FN in the ECM. Data are available via ProteomeXchange^68^ with identifier PXD043976.

## Results

### Mechanical stretching of FN fibers reduces their affinity for TG2 bound to GDP

To mimic the high content of fibrillar FN in the extracellular environment more closely, we employed a well-established *in vitro* FN-fiber stretch assay in combination with a Förster resonance energy transfer (FRET) nanoscale FN tensional sensor, which has already allowed us to identify a number of mechano-regulated binding partners of fibrillar FN^59,60,62,69^. In this assay, fibers are manually pulled with a needle tip from a droplet of FN in solution and deposited onto an elastic silicone membrane mounted on a custom-made stretch-device^58^. FN-fibers were pulled and deposited either only parallel to the stretch axis, or both parallel and perpendicular to the stretch axis (Fig. 2A). By adjusting the strain of the silicone membrane, FN-fibers can be either stretched or relaxed. In this study, FN-fiber tension along the stretch axis of relaxed, native (silicone membrane strain unchanged), and stretched membranes are referred to as low strain (∼20%), medium strain (∼140%) and high strain (∼380%) respectively, as calibrated previously^70^. Though the native membrane is not subjected to any strain, FN-fibers are typically pre-strained to ∼140% due to the forces required to pull them out of the droplet^70^. To provide a direct readout of the conformational distribution within FN-fibers, FN dimers were labeled with multiple FRET donors (AF488) and acceptors (AF546), and to avoid inter-molecular FRET, fibers always contained only 10% of FRET-labeled FN^50^. Due to mechanical stretching, the average distance between the donors and acceptors increases, therefore higher FN-fiber tension corresponds to lower FRET-ratios, while higher FRET-ratios corresponds to more relaxed FN-fibers^50,71^. We controlled the responsiveness of our FN-FRET tensional sensors after labeling by chemical denaturation with progressively increasing concentrations of guanidine hydrochloride (Gdn HCl) using well established protocols^50,62,71^, and observed that as expected, FN-FRET ratio was decreasing as the denaturant caused FN to transition from compact to extended and then partially unfolded conformations in solution (Supplementary Fig. 2). Next, deposited FN-fibers were incubated with Alexa-647 labeled TG2 (TG2-647), and the binding was assessed by measuring the TG2-647 fluorescence intensity, normalizing it pixel-by-pixel to the directly excited FN-FRET acceptor (TG2-647/FN-546), as described previously^62^. We color-coded pixel-by-pixel FN-FRET ratios within Fn fibrils at each externally adjusted strain, to illustrate that FN displays a range of conformations, as also observed in the ECM fibrils assembled by fibroblasts^58^ (Fig. 2B). As expected, the shift toward higher FN-FRET ratios was observed on the relaxed membrane and toward lower FN-FRET ratios on the stretched membrane. Histograms of the distribution of all FN-FRET pixels from a representative fiber, showed that higher FN-FRET ratios corresponded to low strain, and the highest TG2-647/FN-546 ratios were observed on the fiber experiencing low strain (Fig. 2B). To view in one image how TG2 binding affinity changes with FN-fiber strain, we added human recombinant TG2 (hrTG2) to FN-fibers deposited as intersections and incubated in the presence of 1 mM GDP and 1 mM EDTA. We plotted all FN-FRET pixels vs TG2-647/FN-546 pixels from the same intersection as binned scatterplots (Fig. 2C). When the strain of fibers parallel and perpendicular to the stretch axes was different (stretched membrane and relaxed membrane), FN-FRET vs TG2-647/FN-546 pixels segregated into two separate groups with higher TG2-647/FN-546 ratios correlating with higher FN-FRET ratios. However, when the strain along both axes was the same (native membrane), pixels remained clustered as a single group (Fig. 2C). This demonstrates that TG2 binding affinity to FN is dependent on FN-fiber strain. We repeated this experiment using guinea pig liver TG2 (gpTG2), confirming the consistency of this result as well as its reproducibility across species (Supplementary Fig. 4).

**Figure 2:**
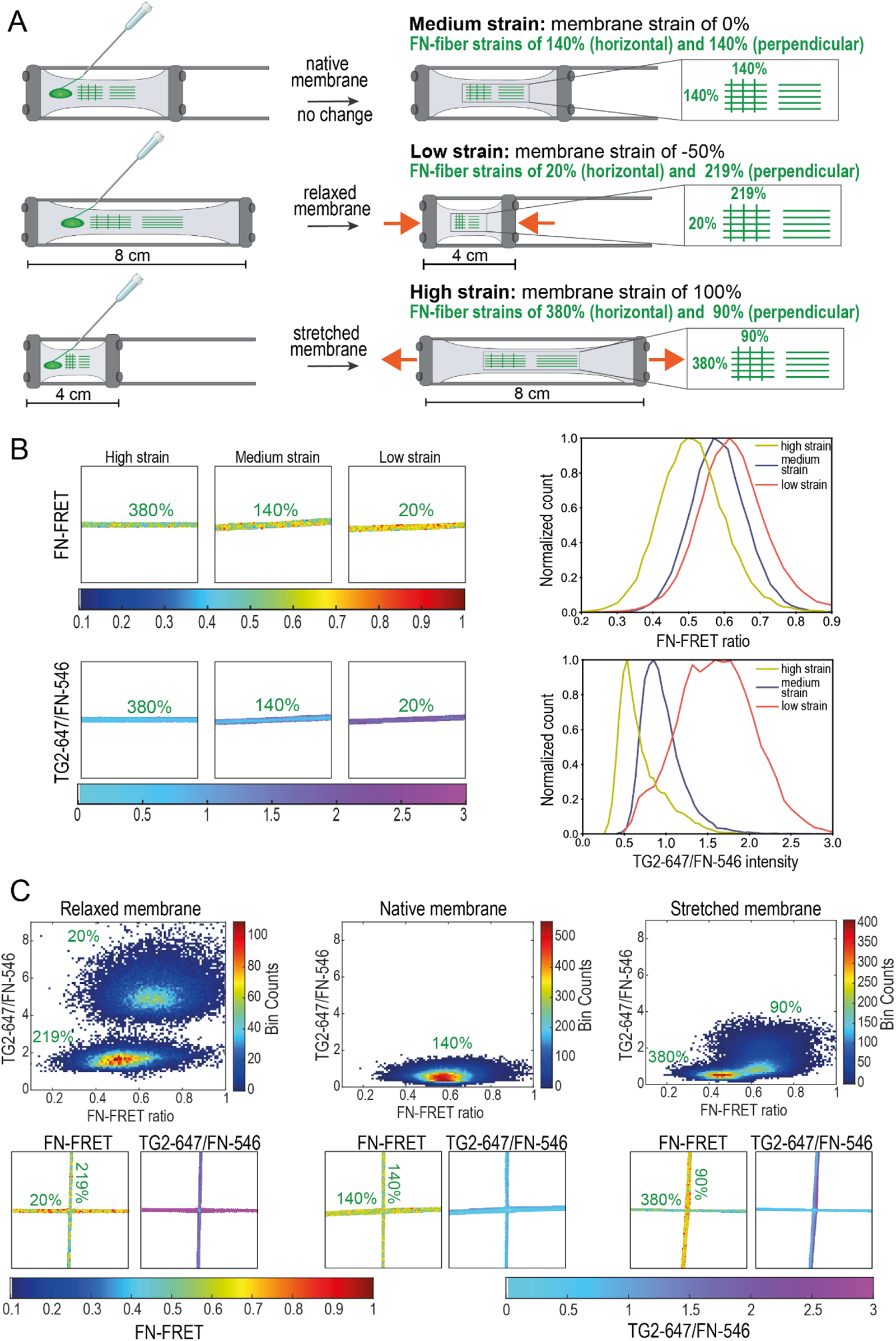
TG2 preferentially co-localizes with FN-fibers under low strain, as revealed by the FN-fiber stretch assay. **A:** A schematic view of the FN-fiber stretch assay and the experimental setup. FN-fibers were manually pulled with a needle from a droplet of concentrated FN and deposited onto a silicone membrane mounted on the custom-made stretch device. FN-fibers were pulled in two ways, either only parallel to the stretch axis, or both parallel and perpendicular as intersections. Membrane was left unchanged (native), relaxed from the pre-strained position, or stretched. Previous calibration studies converted the silicone membrane strain to the corresponding FN-fiber strain^70^ and are indicated on the cartoon. In this study, FN-fiber strains along the stretch axis of relaxed, native, and stretched membranes are referred to as low strain (20%), medium strain (140%) and high strain (380%) respectively. TG2 labeled with an Alexa Fluorophore 647 (TG2-647) was added to the deposited FN-fibers and incubated in the presence of various effectors that change its conformational state. **B: On the left:** Panels show a representative horizontal FN-fiber for low, medium, and high strain with pixel-by-pixel color-coded FRET ratio and normalized TG2-647/FN-546 intensities. In the presence of 1mM GDP and 1mM EDTA, wild type human recombinant TG2 (WT hrTG2) has higher binding affinity (colorcoded magenta) to the FN-fiber under low strain (20%). **On the right:** Distributions of all FRET ratio pixels and normalized TG2-647/FN-546 from each representative fiber shown in the panels on the left were plotted as histograms. Higher FRET-ratio (low strain) in the FN-fiber corresponds to the higher TG2-647/FN-546 signal. **C:** When FN-fibers are deposited as intersections, after membrane is relaxed or stretched, fibers along perpendicular and parallel axes are under different strains. TG2 preferentially binds the fibers under low strain (higher FN-FRET ratio values). All FN-FRET ratio and TG2-647/FN-546 pixels from the same intersection were plotted as binned scatterplots. When the strain of intersecting fibers was different (relaxed and stretched membranes), plotted pixels segregated into two separate groups with higher TG2-647/FN-546 values corresponding to higher FN-FRET ratios (low FN-fiber strain). When the strain of the intersecting fibers was the same (native membrane) plotted pixels remained clustered together as a single group.

We analyzed 15 horizontal fibers deposited parallel to the stretch axis on either native, relaxed or stretched membranes (Fig. 3A). When we incubated FN fibers with GDP-bound TG2 (TG2_GDP_), it preferentially bound to FN fibers under low strain (∼20%). TG2_GDP_ binding to FN fibers under medium (∼140%) or high strain (∼380%) decreased significantly, compared to the low strain. In the past we have demonstrated that stretching of FN-fibers may increase non-specific binding of proteins due to higher exposure of hydrophobic residues which is the consequence of stretch-induced partial protein unfolding^72^. However, we find that the binding affinity of TG2_GDP_ was significantly reduced by mechanical stretching of FN fibers, which suggests that specific TG2 binding sites on FN had been being destroyed. These findings demonstrate for the first time, to the best of our knowledge, that TG2 binding to FN is regulated by the FN-fiber tension and that it can be weakened when FN fibers are mechanically stretched.

**Figure 3:**
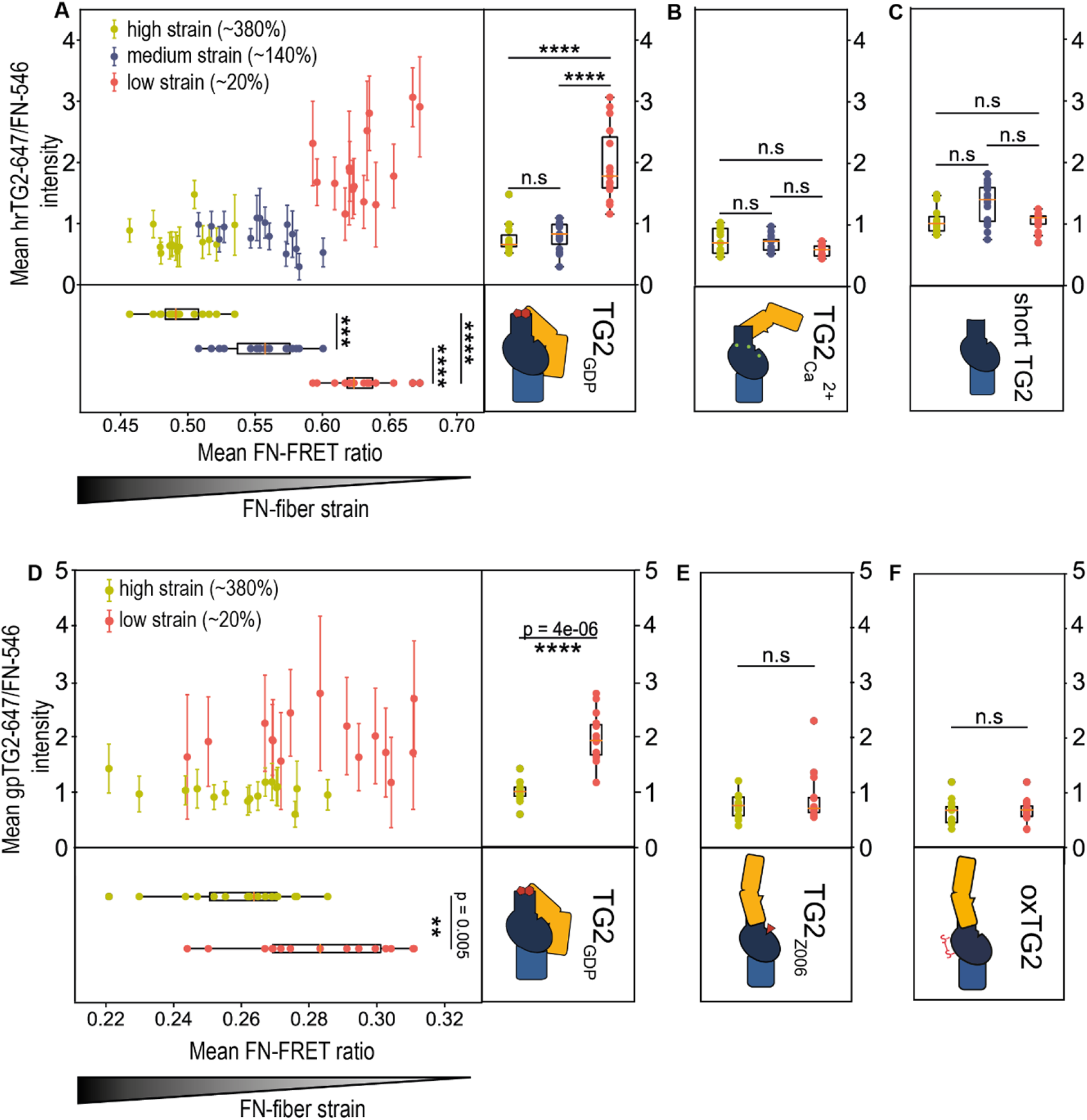
FN-fiber stretch assay data showing the dependence of mechano-regulated TG2-FN binding on TG2’s conformational states and its C-terminal β-barrels. Mean of the distribution of FN-FRET ratio from all pixels of one fiber was plotted against the mean of the distribution of the normalized TG2-647/FN-546 intensity. 15 fibers were analysed per each membrane strain (Full data sets: Supplementary Fig. 6 & 8). **A:** Human recombinant TG2 (hrTG2) has a higher binding affinity toward FN-fibers under low strain (∼20%) in the presence of 1mM GDP and 1 mM EDTA, which induce a closed TG2 conformation. **B:** hrTG2 binds FN with equal affinity in the presence of 10 mM Ca^2+^ which induces an open TG2 conformation regardless of FN-fiber strain. **C:** Short TG2 (β-barrels 1 and 2 are deleted) binds FN with equal affinity regardless of FN-fiber strain. Short TG2 was labelled separately from hrTG2 and has a different degree of labelling, therefore its fluorescent intensity should not be compared to TG2_Ca2+_ or TG2_GDP_. **D:** Like hrTG2 in (A), guinea pig liver TG2 (gpTG2), preferentially bound with higher affinity to FN-fibers under low strain (∼20%) in the presence of 1 mM GDP and 1 mM EDTA (closed-state TG2). When gpTG2 was incubated with the irreversible active-site inhibitor Z006 (Z-DON) **(E)** or oxidised with GSSG **(F)**, both of which induce an extended open-state TG2 conformation, TG2_Z006_ and oxTG2 bound FN equally, regardless of FN-fiber strain. Statistical significance was computed with Wilcoxon rank-sum statistic for two samples. P-values: (*0.01≤ p <0.05; **0.001≤ p <0.01; ***10^-5^≤ p <0.001; ****10^-6^≤ p <10^-5^; ***** p <10^-6^).

**Figure 4:**
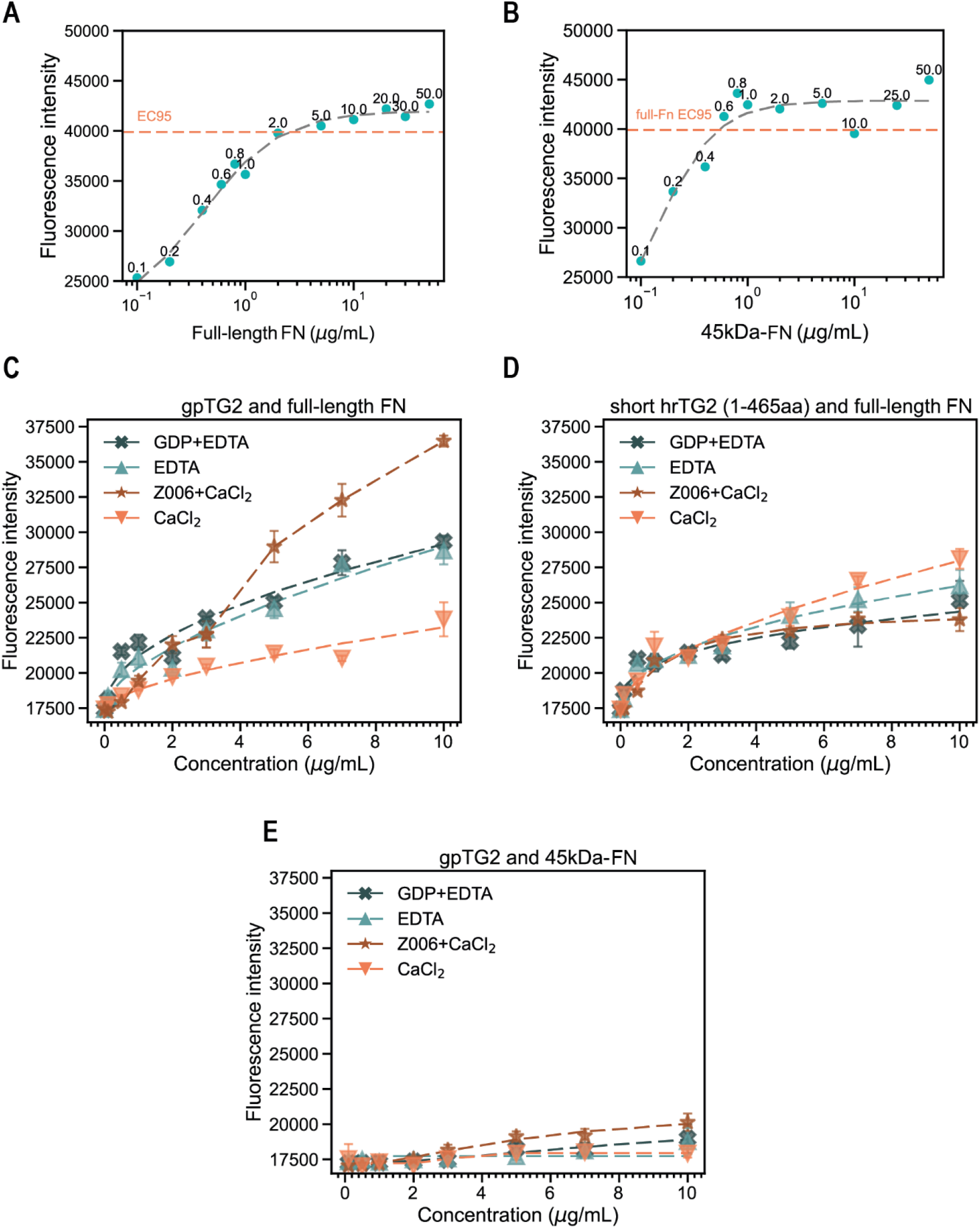
Microplate protein-binding assay data with TG2 binding to surface-adsorbed full-length FN, or to FN’s 45 kDa gelatin binding domain (GBD) fragment. **A:** 96-well microplates were coated with concentration series of full-length FN at 4°C overnight. FN was detected with a monoclonal anti-FN antibody with an epitope within the gelatin binding domain (GBD). From the fitted curve, it was determined that the maximal fluorescent intensity was reached at a concentration of 2.4 µg/ml of full-length FN 95%. **B:** 96-well microplates were coated with concentration series of 45 kDa-FN GBD at 4°C overnight and detected using the same anti-FN antibody as in (A). From the fitted curve, it was determined that the maximal fluorescent intensity observed for the full-length FN was reached at 0.6 µg/ml of 45 kDa-FN GBD 95%, and therefore this concentration corresponds approximately to the same number of TG2 binding sites on 45 kDa-FN GBD. **C:** Binding affinity of wild type guinea pig TG2 (WT gpTG2) to the surface-adsorbed FN changes depending on the presence of GDP, Ca^2+^ and Z006 TG2 effectors. **D:** Deletion of TG2’s β-barrels 1 and 2 results in indiscriminate binding affinity of short TG2 mutant to adsorbed FN, regardless of the presence of GDP, Ca^2+^ and Z006 effectors. **E:** When plates were coated with 0.6 µg/ml of FN’s 45 kDa-FN fragment, which corresponds to the same number of binding sites as when coated with 2.4 µg/ml full-length FN, WT gpTG2 had drastically reduced binding affinity toward 45 kDa-FN fragment. **\*For the coatings of microtiter plates, the 2.4 µg/ml of full-length FN and 0.6 µg/ml of 45 kDa-FN were used, which approximately correspond to the same number of binding sites for TG2, as was determined in by the titration experiments in A and B**.

### Spatial proximity between TG2’s N-terminal Domain and the C-terminal β-barrels is required for its mechano-regulated binding to FN

Since an earlier study suggested that formation of the TG2-FN complex is not altered by the presence of TG2 effectors^45^, we were surprised to find that the mechano-regulated binding was abolished upon addition of saturating amounts of Ca^2+^ (10 mM) (Fig. 3B and Supplementary Fig. 6A). In the presence of Ca^2+^, we no longer detected TG2’s high affinity binding towards FN-fibers under low strain. Instead, calcium bound TG2 (open state) showed the same, low level of binding towards fibers at all strains. A more detailed analysis revealed a dose dependent reduction of TG2 binding to FN fibers under low strain upon titration with increasing amounts of calcium (Supplementary Fig.7).

Most of the TG2 in the ECM is thought to be reversibly inactivated due to the allosteric disulfide bond formation which locks the enzyme in the extended open state^39,73^. A similar conformation is induced by some active-site irreversible TG2 inhibitors, such as an active site-specific inhibitor Z006 (Z-DON-Val-Pro-Leu-OMe, “Z-DON”) ^36,40^ (Fig. 1B). Therefore, we also investigated the binding of oxidized TG2 (oxTG2) and inhibitor Z006 bound TG2 (TG2_Z006_), in comparison to TG2_GDP_. (Fig. 3D-F and Supplementary Fig. 9). To prepare oxTG2 and TG2_Z006_, WT gpTG2 was incubated for 3 h at 37°C with 2 mM oxidized glutathione (GSSG) and 1 mM EDTA, or for 1 h at room temperature with 100 µM Z006 and 1.2 mM CaCl_2_. As before, WT TG2_GDP_ preferentially bound to FN-fibers under low strain (Fig. 3D). In contrast, both oxTG2 and TG2_Z006_ bound FN equally, regardless of the FN-fiber strain (Fig. 3E, F and Supplementary Fig.8). Since this behavior was not anticipated based on previous studies using soluble FN^45^, we next looked for a testable structural hypothesis how TG2 binding to FN could be mechano-regulated. Cardoso *et al.* have mapped the FN binding site on TG2 to a region on the N-terminal domain^45^, as indicated in Fig. 1A. Inspection of the structure suggests that the second β-barrel in the closed state is in proximity to the proposed FN binding site (Fig. 1A). The “beads-on-a-string” like structure of FN (Fig. 1C) suggests that adjacent FN modules could possibly form synergistic contacts, thereby stabilizing the TG2-FN complex on low tension FN fibers. In contrast, when TG2 is in the extended open state, the second β-barrel is out of reach and no additional contacts could be formed. To test whether β-barrels of TG2 participate in the mechano-regulated binding to FN-fibers, we have used a short TG2 variant (1-465aa, β-barrel 1 and 2 deleted) in our FN-fiber stretch assay. Indeed, short TG2 showed the same behavior as calcium activated WT TG2, oxTG2 and TG2_Z006_ and exhibited no mechano-regulated binding to FN-fibers (Fig. 3C and Supplementary Fig. 6B). This suggests that for the mechano-regulated binding, the β-barrels of TG2 need to be present and must be in spatial proximity to the N-terminal domain. Interestingly, there was no significant difference in binding between oxTG2 and TG2_Z006_ to FN-fibers under high strain (∼380%), however TG2_GDP_ binding affinity was significantly higher compared to oxTG2 and TG2_Z006_ on high strain (Supplementary Fig. 8C). Upon mechanical stretching of FN-fibers, a large distribution of conformational FN states exists within each FN-fiber that upon stretching gets gradually more shifted towards partially unfolded states^74^. Thus, the ensemble of specific binding sites on FN gets gradually destroyed by fiber stretching, not at one specific strain. This is due to the non-periodic bundling of FN molecules within the fibers, which interferes with the mechanical hierarchy in which FN domains would get unfolded in isolated molecules^74^. Consequently, the specific TG2-binding sites on FN are being destroyed gradually with increasing FN-fiber strain, as observed previously for other binding partners^50,58,60,62,70^. Thus, FN fiber stretching results in a progressive reduction of TG2_GDP_ binding. However, since the N-terminal domain and C-terminal β-barrels of TG2_GDP_ are in spatial proximity, unlike in oxTG2 and TG2_Z006_, TG2_GDP_ can maintain additional contacts to the remaining intact binding sites on FN. Thus, this result also agrees with our hypothesis.

### Catalytic activity of TG2 has no effect on the loss of mechano-regulated binding to fibrillar FN

To ensure that the loss of the mechano-regulated binding of TG2 to FN-fibers in the presence of calcium was not due to TG2 crosslinking activity or TG2 self-crosslinking^75^, we conducted a control experiment using an inactive Cys277Ser-TG2 mutant (Supplementary Fig. 9). TG2 is stabilized in the closed conformation by an H-bond between Cys277 (catalytic core) and Tyr516 (β-barrel 1)^76^. Cys277Ala/Ser-TG2 mutants which are less efficient in forming this H-bond, are known to assume predominantly an open conformation. We observed that in the presence of 1.2 mM calcium, the Cys277Ser-TG2 mutant bound FN-fibers at both high (∼380%) and low (∼20%) strains equally well, indicating that it is the open conformation of TG2 that abolishes mechano-regulated binding to FN-fibers, and not its cross-linking activity. Addition of 1 mM GDP increased the Cys277Ser-TG2 binding affinity to FN-fibers at low strain (∼20%); however, the significance of this increase was considerably less (p = 0.004) compared to the wild type hrTG2 (p < 10^-5^). Cys277/Ala/Ser-TG2 mutants are less efficient in binding GDP and tend to assume the open conformation^76^, thus explaining the reduced significance of the observed increase as well. All these results confirm that the conformational state and not the crosslinking activity of the open-state TG2 plays the central role in the loss of the mechano-regulated binding to FN-fibers.

### TG2’s affinity to surface-adsorbed FN depends on its conformation and the C-terminal β-barrel domains

While previous studies suggested that TG2 binding to surface-adsorbed FN is independent of TG2 conformational states and C-terminal domains^45^, our conclusions drawn from FN-fiber stretch assays supported the opposite conclusion. To understand whether our results were due to the use of the fibrillar FN in our stretch-assays, which has distinct conformational states compared to surface adsorbed FN that retains a more closed conformation^77^, we repeated the studies using surface adsorbed FN. For this, 96-well microtiter plates were coated with human plasma FN or 45-kDa FN gelatin-binding domain (GBD) at concentrations ranging from 1 µg/ml to 50 µg/ml overnight at 4°C. FN was detected using a monoclonal antibody that has its epitope localized within FN’s gelatin binding domain (GBD) (MAB1892, Merck), followed by staining with a secondary AF488-coupled antibody, and the fluorescence intensity was measured with a plate reader. To determine the optimal FN concentration for the plate coatings, we first plotted the adsorption isotherm for full-length dimeric FN (Fig. 4A). The concentration of full-length FN, at which 95% of maximal fluorescence intensity was observed, corresponded to 2.2 µg/ml, and was used for the FN coatings in all subsequent microplate assays. To determine the concentration of the GBD that would enable the coating of plates with a similar number of TG2 binding sites as with the full-length FN, we plotted the GBD adsorption isotherm, as we did for the full-length FN (Fig. 4B). We determined that the concentration of the GBD corresponding to 95% of the maximal fluorescence intensity of the full-length FN was 0.6 µg/ml, which we used to coat plates for subsequent experiments. To evaluate the binding of TG2 to adsorbed FN, we titrated fluorescently labelled TG2 in presence of 1 mM GDP and 1 mM EDTA, or only 1 mM EDTA, or 1.2 mM CaCl_2_, or 1.2 mM CaCl_2_ and 100 µM Z006, followed by incubation on full-length FN-coated wells for 1 h at room temperature (Fig. 4C). Unexpected from previous reports, we found that TG2 had different binding affinities to adsorbed FN, which were in fact dependent on the conformational state of TG2 induced by a specific TG2 effector. Specifically, the binding affinities of TG2 to surface adsorbed FN were in the following order from lowest to highest: TG2_CaCl2_ < TG2_EDTA_ < TG2_GDP_ < TG2_Z006_. In contrast and in agreement with our hypothesis, the short TG2 (1-465aa, β-barrel 1 and 2 deleted) exhibited comparable binding affinities under all four conditions, irrespective of the effector (Fig. 4D). The maximum binding affinity was achieved at a concentration of approximately 2 µg/ml of short TG2, but non-specific binding increased continuously at concentrations above 4 µg/ml. Furthermore, we observed that the binding affinity of TG2 significantly decreased when plates were coated with the GBD, even when the same number of TG2 binding sites was present (Fig. 4C, E). These findings indeed suggest that TG2 has a higher binding affinity for full-length FN, likely due to additional contacts it makes with FN modules outside of the conventional GBD.

These results demonstrate that the affinity by which TG2 binds to surface-adsorbed FN is tuned by the conformation of TG2 as induced by its allosteric effectors. Moreover, our data confirm that the C-terminal domains of TG2 play a crucial role in enhancing the binding to surface-adsorbed FN, highlighting their significance in TG2 interactions with both fibrillar and surface-adsorbed FN.

### C-terminal β-barrels of TG2GDP interact with the regions outside of the canonical gelatin-binding domain (GBD) of FN in solution

As the short form of TG2 without β-barrel domains no longer exhibited sensitivity to the FN-fiber strain and bound both stretched and relaxed FN-fibers equally, this implies the presence of a secondary binding site on C-terminal β-barrel domains of TG2. Yet, no information was available in the literature about the role of these domains in TG2’s interaction with FN. To explore the possibility of multivalent synergy binding sites, we used crosslinking mass spectrometry^63^ (XL-MS) to map contact sites between the two proteins. Previous similar proteomic approaches (hydrogen-deuterium exchange – HDX) using only FN’s GBD in complex with TG2_Z006_, did not identify any regions on the C-terminal β-barrel domains of TG2 that could interact with FN^45^. This is in agreement with our hypothesis, as the inhibitor Z006 is known to trap TG2 in the extended open conformation, which spatially separates the C-terminal domains from the N-terminal domain, making them inaccessible to FN^36^. To our knowledge, interacting regions of the TG2_GDP_ (closed state) and the full-length FN have not been mapped with both approaches. Thus, we performed for the first time XL-MS analysis of TG2_GDP_ in complex with full-length FN, and to better compare our findings with literature, of TG2_GDP_ in complex with the GBD.

Amine-specific crosslinking of the TG2-FN complexes was performed with disuccinimidyl suberate (DSS), and to increase the coverage of structural information, simultaneous crosslinking of acidic residues (Asp/Glu with Asp/Glu and Lys with Asp/Glu) was performed with PDH and DMTMM^78^. Crosslinked samples were further processed with trypsin or chymotrypsin digestion, SEC fractionation of the resulting peptide mixtures, followed by LC-MS/MS analysis of SEC fractions. Among the resulting FN isoforms, we were able to unequivocally identify FN isoform 1, which was used for the subsequent data analyses. Validated crosslinks were displayed on circular diagrams with the help of xVis^79^ (Fig. 5 and Supplementary Fig. 10-11). In total, 20 unique crosslinks were identified between TG2_GDP_ and FN or TG2_GDP_ and GBD complexes (Supplementary Note 2 and Fig. 5). Remarkably, the C-terminal β-barrel 2 was in close contact with FNIII_14-15_ (4 unique inter-protein crosslinks), as well as with FNI_2_ (3 interprotein crosslinks). Simultaneously, FNIII_14-15_ and FNI_2_ were in close contact with each other, as indicated by multiple intra-protein crosslinks between those regions (magenta lines in Fig. 5). From experiments with the TG2 and GBD and chymotrypsin as a protease, 8 additional unique inter-protein crosslinks were detected (Supplementary Fig. 11). Notably, we identified a crosslink between Lys30 (TG2) and Lys486 (GBD, numbering for full length FN). TG2 residue Lys30 is known to be one of the three residues (Lys30, Arg116, His134) that comprise the main FN-binding site on the N-terminal β-sandwich of TG2^45^. Interestingly, the crosslink involving Lys486 was not found when TG2 was in complex with the full-length FN (Supplementary Note 2 and Supplementary Fig.10). Instead, Lys30 on TG2 was crosslinked to Lys1837 and Lys1862 on FNIII_14_. This difference could arise from the variations in reactivity between these lysine residues and structural differences. These MS data clearly demonstrate that the C-terminal domains of TG2 interact with regions of FN outside of the canonical GBD when FN is in a compact quaternary conformation in solution, and when the C-terminal β-barrels are in spatial proximity to the N-terminal domain.

**Figure 5:**
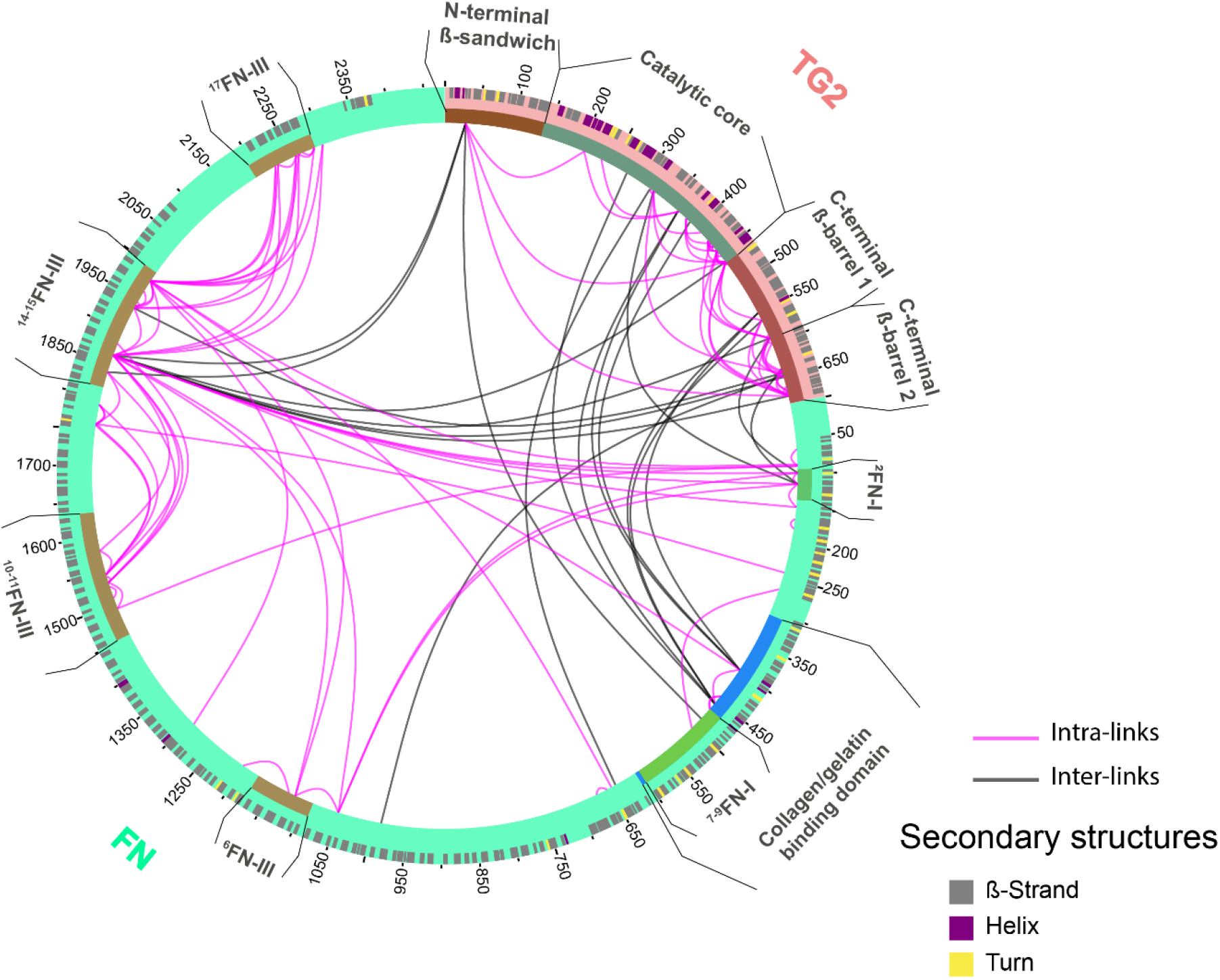
Cross-links identified with the XL-MS of the TG2_GDP_ and FN complex reveal that TG2’s C-terminal β-barrel domains interact with regions outside of the canonical GBD. **A:** Circular diagram shows simultaneously 20 unique inter-protein crosslinks identified from both TG2-FN and TG2-45 kDa-FN complexes using trypsin and chymotrypsin as proteases. To view them separately, go to Supplementary Fig. 10-11. Crosslinks only within TG2 or only within FN (intra-protein crosslinks) are shown in magenta, crosslinks between TG2 and FN (inter-protein crosslinks) are shown in black. C-terminal β-barrel 2 of TG2_GDP_ is in close contact with FNIII_14-15_ (Lys1862 and Lys1936) and FNI_2_ (Lys100 and Lys116). Simultaneously, FNIII_14-15_ and FNI_2_ are in proximity to each other, as shown by magenta intra-protein crosslinks. TG2 residue Lys30, which is one of the three residues (Lys30, Arg116, His134) comprising the main FN-binding site on the N-terminal β-sandwich^45^, was crosslinked to Lys486 (45 kDa-FN, numbering for full-length FN). On the full-length FN, Lys30 was crosslinked to residues Lys1837(FNIII_14_) and Lys1862(FNIII_14_). Visualization was done with xVis Webserver^79^.

### Crosslink-guided structural modelling of TG2 and FN

XL-MS not only identifies which regions of a protein complex are in close contact, but also provides valuable spatial information through physical distance restraints imposed by each crosslinker. The spatial information can be used for modelling and docking to determine the position and orientation of within a complex^80^. To determine the binding interface of the TG2-FN complex using low-resolution restraint data more accurately, we integrated the experimental crosslinks into a modelling pipeline^81,82^ (Supplementary Fig. 13). We selected a few regions of FN for modelling based on criteria such as the availability of templates in PDB, coverage by crosslinks, and existing knowledge in the literature regarding binding sites. The regions that met these criteria and were suitable for structural modelling were FNI_2-3_, FNI_7-9_, FNI_6_FNII_1-2_ FNI_7-9_ (GBD), and FNIII_14-15_ (Supplementary Note 1). To build crosslink-guided models, we used the I-TASSER^83^ and submitted experimental intra-protein crosslinks as distance restraints to structurally refine the available crystal structure templates, as was previously done^81^. To evaluate the compatibility of refined models with experimental restrains, we calculated Euclidean distances (ED) between β-carbons (CB-CB) of crosslinked residues using Xwalk^84^. We classify crosslinks into compatible and non-compatible based on the distance cut-off values^78,81^: ED for DSS < 35 Å, ED for DMTMM < 25 Å, ED for PDH < 35Å.

We identified 4 intra- and 3 inter-protein crosslinks for FNI_2-3_. A high-resolution crystal structure of FNI_2-3_ was available (PDB:2cg7), which we submitted as a template to I-TASSER for structural refinement with experimental crosslinks. After structural refinement, a high scoring model was selected, that satisfied all crosslinks within the distance cut-off (Supplementary Note 1). For FN’s GBD, we detected 3 intra- and 8 inter-protein crosslinks. Templates covering FNI_6_FNII_1-2_FN_7_ and FNI_8-9_ domains, except for the short linker (513-516aa) between them, were available as PDB:3mql and PDB:3ejh, respectively. However, our initial attempt to model the entire GBD with I-TASSER was not satisfactory. Thus we used ROBETTA^85^ and AlphaFold^86^ to determine the orientation of FNI_6_FNII_1-2_FNI_7_ and FNI_8-9_ with respect to each other. To evaluate the quality of predicted models of GBD, we submitted them to QMEAN^87^, which evaluates how the model performs in comparison to the experimental PDB structures. The AlphaFold predicted that the GBD model outperformed the ROBETTA model (QMEAN Z-score=1.44 vs 1.86). This indicates better compatibility of the structural model predicted by AlphaFold than ROBETTA with the experimentally determined crosslinks. Consequently, we selected the AlphaFold predicted model of GBD for docking with HADDOCK^64^.

Although we detected only one inter-protein crosslinks and no intra-protein crosslinks for the FNI_7-9_ domains, we included it in the modelling pipeline. FNI_7-9_ is well-known to be the high affinity binding region for TG2 and behaves as the whole 45-kDa GBD in mediating functions such as adhesion, spreading and migration^43^. To model FNI_7-9_, we followed the same strategy we used for the GBD. The AlphaFold predicted model of FNI_7-9_ deviated less than one standard deviation from the experimental structures deposited in the PDB (QMEAN Z-score=0.64), whereas the ROBETTA predicted model deviated by two standard deviations (QMEAN Z-score=2.02). Therefore, we selected the AlphaFold predicted model of FNI_7-9_ for further docking steps, given its overall better performance (Supplementary Note 1).

We identified 11 intra- and 7 inter-protein crosslinks for FNIII_14-15_ and submitted the high-resolution crystal structure of FNIII_14-15_ (PDB:1fnh) along with crosslinks for structural refinement using I-TASSER. A high scoring model that satisfied 10/11 intra-protein crosslinks was selected. The non-compatible crosslink (Lys1862-Lys1936) connects two interdomain residues located on FNIII_14_ and FNIII_15_. As these two domains are connected by a flexible linker, it is expected that structural rearrangements favouring a more closed conformation might occur.

For TG2, we detected 46 intra-protein crosslinks, 8 of which violated distance cut-off when we mapped them on the model. All the non-compatible crosslinks were between the interdomain residues. Specifically, 5 non-compatible crosslinks were between residues on the catalytic core and C-terminal domain, 2 between residues on C-terminal domains, and 1 between N-terminal domain and the catalytic core (Supplementary Note 1). As these domains are connected by flexible linkers, non-compatible residues suggest that TG2 undergoes some structural rearrangement upon binding to FN in solution. After performing structural refinement, one notable improvement was residue Lys30, which on the available TG2 template (PDB:4pyg) did not comply with our solvent accessibility criteria (Supplementary Note 1). The Lys30 residue was previously shown by mutagenesis studies to be crucial for FN-binding^45^. After recalculation of TG2 template using experimental intra-protein crosslinks, all three residues comprising the high-affinity FN-binding site (Lys30, Arg116, His134) satisfied the solvent accessibility criteria. Overall, by integrating experimental crosslinks as distance restraints, we were able to structurally refine available templates to be in better agreement with the experimental MS data.

### Crosslink-guided HADDOCK docking of TG2 with FN7-9 revealed a parallel alignment of FN modules with TG2’s C-terminal β-barrels

Protein complexes in solution are typically dynamic, and crosslinks can reflect an averaged ensemble of conformations. However, non-compatible crosslinks that may belong to an alternative complex conformation can negatively impact the accuracy of docking results. To address this issue, we utilized the DisVis webserver, which filters out non-compatible crosslinks and predicts key residues involved in binding at the interaction interface using experimental crosslinks^65^. Our analysis using DisVis highlighted three putative incompatible crosslinks, namely the Lys319(TG2)-Glu116(FN) crosslink between TG2 and FNI_2-3_, the Lys550(TG2)-Lys397(FN) crosslink between TG2 and GBD, and the Lys464(TG2)-Lys1862(FN) crosslink between TG2 and FNIII_14-15_ (Supplementary Note 2). While it is important to note that these crosslinks may not necessarily be false and are likely to belong to an alternative conformation of the complex, we did not further investigate this possibility due to insufficient number of crosslinks. Therefore, we excluded these crosslinks from further docking steps with HAD-DOCK to improve the accuracy of the docking results.

To identify the putative active residues at the binding interface, we filtered out the residues with less than 40% relative solvent accessibility for the backbone or side chain using NACCESS^88^, and then used DisVis interaction analysis to predict active residues at the binding interface consistent with submitted experimental crosslinks (Supplementary Note 2). Active residues, along with confirmed crosslinks mutually compatible after selection by DisVis, were submitted for separate docking runs with HADDOCK for the TG2 and FNI_2-3_, TG2 and FNI_7-9_, TG2 and FnIII_14-15_, TG2 and GBD complexes (Supplementary Note 3). After docking, we mapped crosslinks of predicted models and used Xwalk to calculate distances between crosslinked residues to identify any non-compatible crosslinks. The models of TG2-FNI_2-3_ and TG2-FNIII_14-15_ complexes satisfied all crosslinks that were submitted to guide docking, and the models with the highest HADDOCK score were selected (Fig. 6A, B). In the case of the TG2-FN_7-9_ complex, we guided docking by a single crosslink, which was compatible on all resulting models, thus we selected the model with the highest HADDOCK score. Notably, one of the predicted models of the TG2-FN_7-9_ complex supported the parallel alignment of FNI_8-9_ with C-terminal β-barrels of TG2, while FNI_7_ was aligned with the high-affinity site on the N-terminal domain of TG2 (Lys30, Arg116, His134) (Fig 6C). This docking pose strongly supports the hypothesis of multivalent binding sites and that suggests a structural mechanism that can indeed explain the mechano-regulated TG2 binding to fibrillar FN.

**Figure 6:**
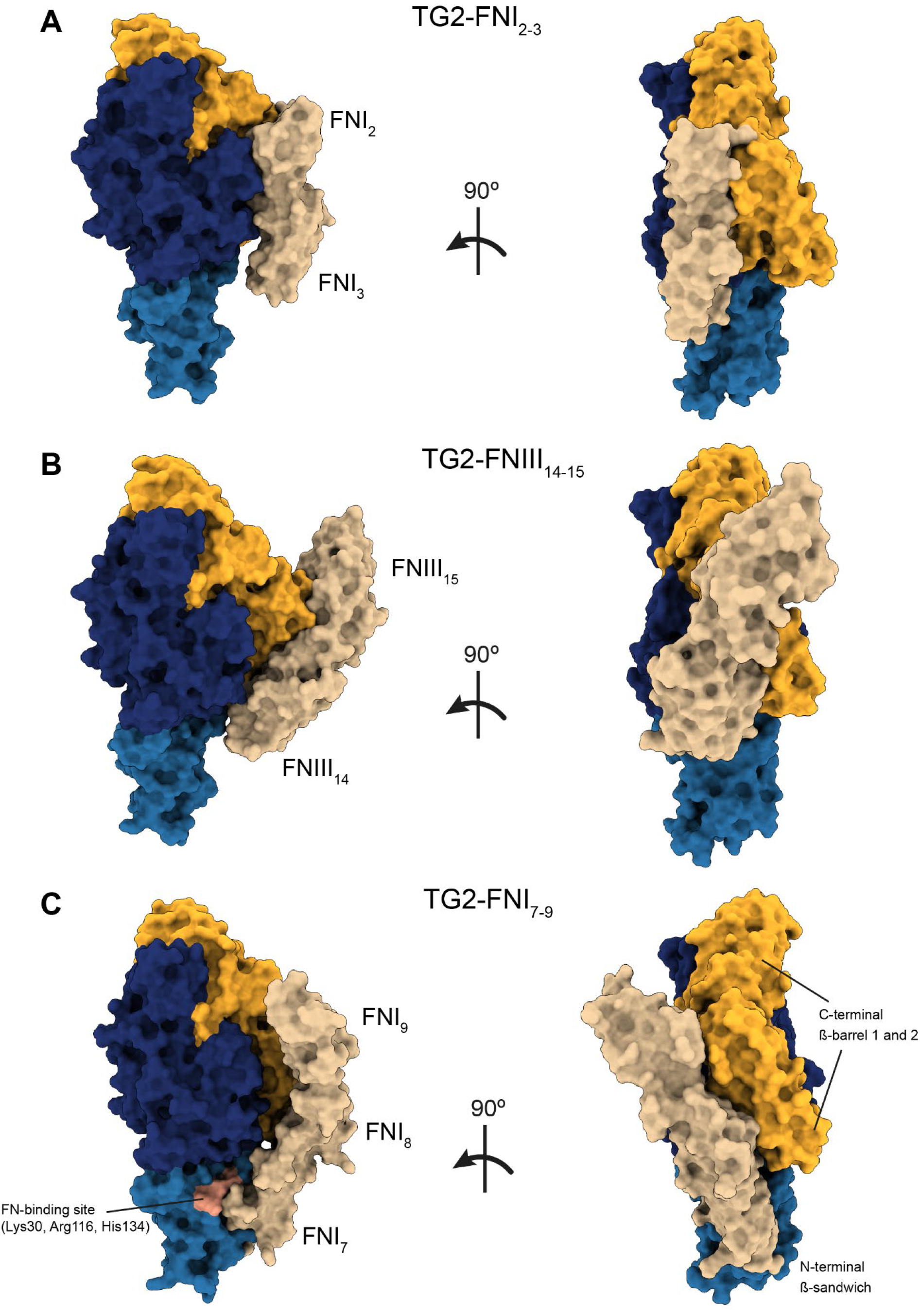
Protein-protein docking with HADDOCK of TG2 complexed with selected FN fragments guided with experimental crosslinks as distance restraints. Predicted models of **(A)** TG2 in complex with FNI_2-3_. **(B)** TG2 in complex with FNIII_14-15_, and **(C)** TG2 in complex with FNI_7-9_. The resulting docking pose of the TG2-FNI_7-9_ complex supports parallel alignment of FNI_8-9_ with the C-terminal β-barrels of TG2, while FNI_7_ is in contact with the canonical FN-binding site (Lys30, Arg116, His134, colored pink) on the N-terminal β-sandwich. Full data set with mapped crosslinks is available in Supplementary Note 3 and Supplementary Fig. 15-17.

### Collagen-mimicking peptide R1R2 and TG2GDP compete for binding to FN-fibers

To better understand the results of XL-MS experiments, where we used soluble FN in a compact conformation, in relation to the results of in FN-fiber stretch assay, in which FN was in fibrillar form, we sought to validate the predicted model for the TG2-FN complex. Previous studies have identified the TG2 and collagen binding sites on FN within the 45-kDa GBD^48,89^. Although both binding partners interact with several FN modules, FNI_8_ appears to be particularly important for collagen as well as for TG2 binding^43,89^. Our predicted model of TG2-FNI_7-9_ complex suggested an overlap between TG2 and collagen-binding sites on FNI_8_ within the GBD. To validate this model, we investigated whether simultaneous binding of TG2 and collagen to fibrillar FN is possible. To test this, we utilized the R1R2 collagen-mimicking peptide, which is derived from the SFS FN-binding protein of the pathogen *Streptococcus equi*^66^. Both R1R2 and collagen share a conserved GEXGE motif which has been shown by crystallography to bind to FNI_8-9_, and in accord with this R1R2 inhibits binding of FN to collagen I^67,89^. We could furthermore show previously that the R1R2 binding motif on FN is destroyed by stretching FN fibers^60^.

To assess the influence of collagen binding, we added increasing concentrations of R1R2 peptide to 100 μg/ml of TG2_GDP_ in 1 mM GDP, 1 mM EDTA, 1 mM MgCl_2_ in 50 mM Tris and incubated on FN fibers under low strain for 1 h. This R1R2 titration series revealed a dose dependent reduction of TG2 binding to FN, indicating that R1R2 can disrupt the specific interactions between TG2_GDP_ and FN fibers under low strain (Fig. 7A). Next, we tested how TG2 would bind FN-fibers in the presence of 100 µM R1R2 at all strains. Indeed, in the presence of high concentrations of R1R2, TG2 equally bound to FN-fibers at all strains (Fig. 7B). Interestingly, the addition of 100 µM R1R2 further significantly reduced TG2_GDP_ binding to the FN fibers under high strain, indicating that TG2 and R1R2 are still competing for the remaining specific binding sites on FN (Supplementary Fig. 12). We previously reported that FN fibers under high strain are more prone to non-specific interactions due to the exposure of hydrophobic sites^72^. Our findings thus confirm that the binding of TG2 to the FN fibers under high strain is still specific, but weaker as the ensemble of destroyed binding epitopes increases by stretching FN fibers. Therefore, our data demonstrate for the first time that the collagen-mimicking peptide R1R2 and TG2 directly compete on fibrillar FN under low as well as high fiber strains.

**Figure 7.**
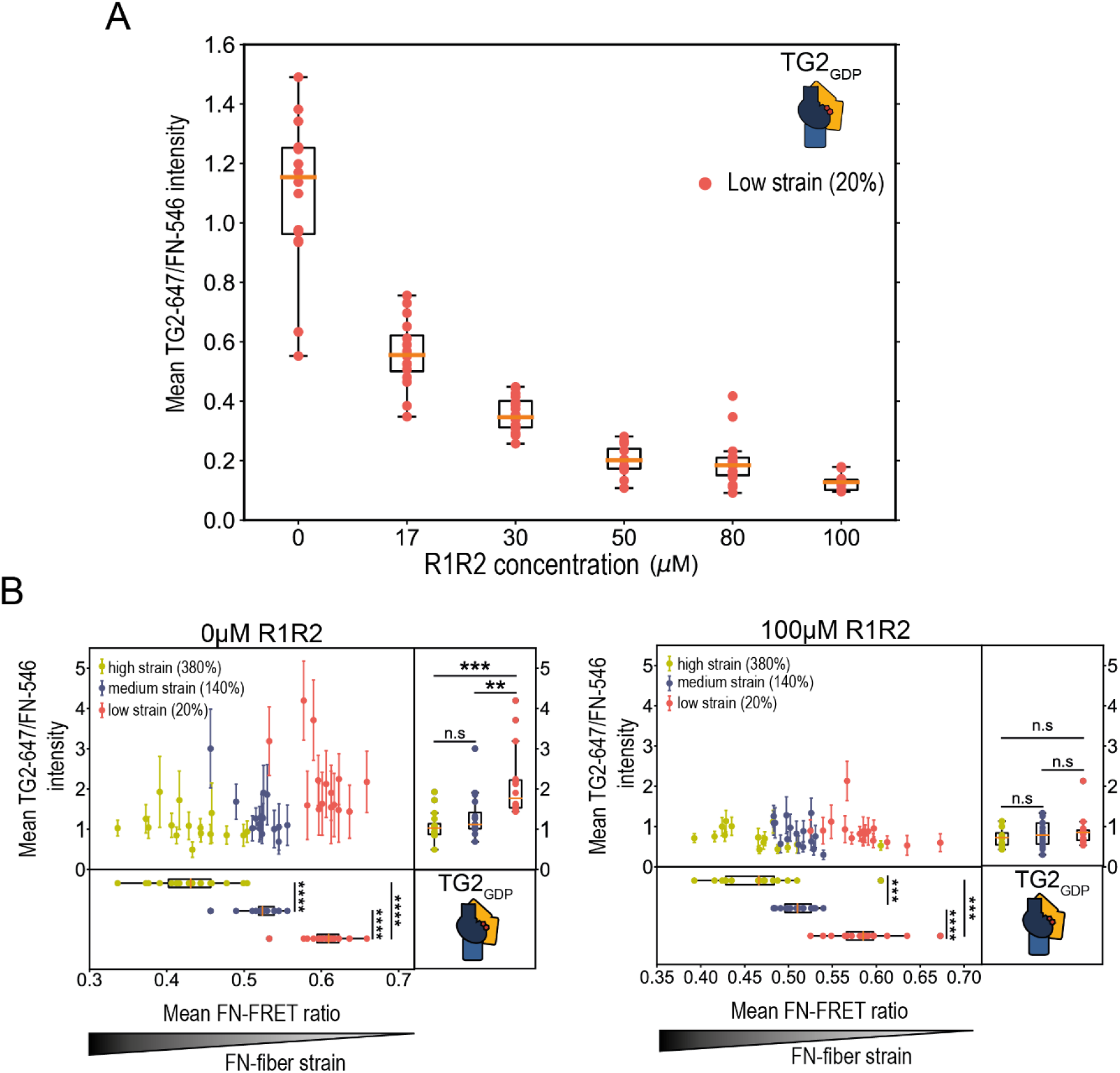
FN-fiber stretch assay reveals a dose-dependent competition between the collagen-mimicking peptide R1R2 and WT gpTG2_GDP_ for binding to FN-fibers under high, as well as low strain. **A:** In the presence of increasingly higher concentrations of R1R2, which targets FN’s GBD, TG2_GDP_ binding to the FN-fibers under low strain reduces in a dose-dependent manner. **B:** Saturating concentrations of R1R2 abolished mechano-regulated TG2 binding to FN, thus limiting TG2 binding to its known canonical binding site. In the presence of 100 µM R1R2, TG2 bound equally to FN-fibers under high, medium, and low strain. 15 fibers were analysed per each membrane strain. Statistical significance was computed with Wilcoxon rank-sum statistic for two samples. P-values: (*0.01≤ p <0.05; **0.001≤ p <0.01; ***10^-5^≤ p <0.001; ****10^-6^≤ p <10^-5^; ***** p <10^-6^)

## Discussion

Since nothing was known about the interaction of TG2 with fibrillar FN, and how ECM fiber tension might modulate TG2’s binding to FN fibers addressed for the first time whether this interaction might be regulated by the tensional state of FN-fibers. By utilizing our well-established *in vitro* FN-fiber stretch assay in combination with a FN-FRET nanoscale strain sensor^58,62^. We discovered that TG2 binds with higher affinity to FN-fibers under low strain (relaxed), rather than stretched fibers, since fluorescently labeled TG2-647 preferentially co-localized with those FN-fiber pixels that showed higher FN-FRET ratios (Fig. 2C and Supplementary Fig. 4). TG2 binding affinity to FN-fibers could thus be significantly reduced by stretching the FN-fibers (Figs. 2C and 3A). We further established that the spatial proximity of TG2’s C-terminal β-barrels to its N-terminal domain is necessary for such mechano-regulated binding to occur. Confirming this notion, the mechano-regulated binding could not be observed when its C-terminal β-barrels 1 and 2 were deleted (Fig. 3C), or when TG2 was stabilized in the open conformation in the presence of calcium (Fig.3B), or when TG2 was locked in the open effector-free conformation by the Z-DON inhibitor (Fig.3E), or finally by oxidation (Fig.3F). Further supporting our proposition, TG2 bound to both stretched and relaxed FN-fibers equally regardless of the FN-fiber strain, when the two C-terminal domains were deleted, or when they were not in spatial proximity with the N-terminal domain. Importantly and as control, by using the TG2 inactive mutant Cys277Ser, we established that this mechano-regulated binding was not dependent on the TG2 catalytic activity but was sensitive only to TG2’s conformational states (Supplementary Fig. 9).

TG2 demonstrated differential binding affinities not only to fibrillar FN, but also to surface adsorbed FN, which adopts a more compact conformation compared to FN-fibers (Fig. 4A-D). Again, TG2 binding to surface adsorbed FN was dependent on TG2’s conformation and the presence of its C-terminal β-barrels. Also, for the surface-adsorbed 45 kDa-FN fragment comprising modules FNI_6_FNII_1-2_FNI_7-9_, we confirmed that TG2’s binding affinity decreased drastically when an equivalent number of binding sites was presented, instead of full-length FN (Fig. 4E). This finding is remarkable, as it contrasts with a previous study that reported TG2 binding to the GBD with the same affinity as full-length soluble FN^48^. However, the contrasting result can likely be explained by the fact that the TG2 binding between GBD and full-length FN was in that previous study not compared in terms of the equivalent number of binding sites available to TG2. Collectively, these results indicate that the C-terminal domains of TG2 support its interaction with FN, in synergy with its canonical binding site, and that this interaction thus enhances the TG2’s affinity to FN beyond the known GBD interaction. This conclusion was confirmed by XL-MS results, which showed that TG2’s C-terminal β-barrels interact with regions of FN outside of the canonical GBD region, specifically with FNI_2_ and FNIII_14-15_ (Fig. 5). Notably, an earlier study using rotary shadowing electron microscopy reported a possible TG2 binding to FNI_4-5_ regions alongside the conventional GBD^90^. However, subsequent studies were not able to detect this interaction, possibly due to the use of separate FN fragments as opposed to the synergy site localization^43,44,46^. This XL-MS result is highly significant, since the studies that aim to develop small molecule molecular inhibitors of TG2-FN interaction, often utilize 45-kDa FN GBD instead of the full-length FN^32,33^. While our XL-MS data did not definitively indicate whether TG2’s C-terminal domains interact with FNI_2-3_ and FNIII_14-15_ simultaneously, such simultaneous interaction is suggested by spatial proximity of FNI_2-3_ and FNIII_14-15_, as is evident from numerous inter-protein crosslinks within FN connecting those two regions (Fig. 5).

Using XL-MS and crosslink-guided structural modeling tools, we have shown that, TG2 residues on both N-terminal β-sandwich and C-terminal β-barrels are predicted to interact with FN, but only when TG2 adopts a closed conformation (Fig. 6). It is important to note that a compact conformation of soluble FN, which was present in our XL-MS experiments must be distinguished from fibrillar FN. When FN is in the soluble compact conformation, FNI_2-3_ and FNIII_14-15_ modules are brought in spatial proximity attracted by long-range electrostatic interactions, but are spatially separated when FN assumes a fibrillar form^51^. However, since FN molecules are not periodically arranged in FN fibers, yet are closely-packed^53,74^, FNI_2-3_ and FNIII_14-15_ modules from some neighboring molecules could happen to be in the right distance. Additionally, one of the predicted docking models of TG2 with FNI_7-9_ suggested parallel alignment of the FNI_8-9_ modules with the C-terminal β-barrels of TG2, while FNI_7_ was in contact with the main FN-binding site (Lys30, Arg116, His134) on N-terminal domain (Fig. 6C). Such an orientation also would allow for additional stabilizing contacts between C-terminal domains and fibrillar FN and could mediate the mechano-regulated binding between TG2 and FN-fibers. Therefore, it is possible that mechano-regulated TG2 binding to FN-fibers is mediated by C-terminal TG2 domains synergistic interactions with FNI_2-3_, FNI_8-9_ or FNIII_14-15_ modules offering multiple options for the high affinity interaction that we observed in our FN-fiber stretch assays. Bringing our findings that TG2 binding is modulated by FN-fiber tension together with existing literature, we propose a novel model of mechano-regulated TG2-FN interaction (Fig. 8). When TG2’s N-terminal and C-terminal domains are in spatial proximity due to GDP/GTP binding or interaction with syndecan-4, TG2 and FN can form a high affinity interaction due to C-terminal β-barrels making synergistic contacts with FNI_8-9_ (Fig. 8A) and/or FNI_2-3_ and FNIII_14-15_ (Fig 8B). When TG2 assumes an open or effector-free open conformation due to calcium binding or oxidation, respectively, only low affinity interaction can be formed and mechano-regulated binding to FN-fibers is not possible.

**Figure 8:**
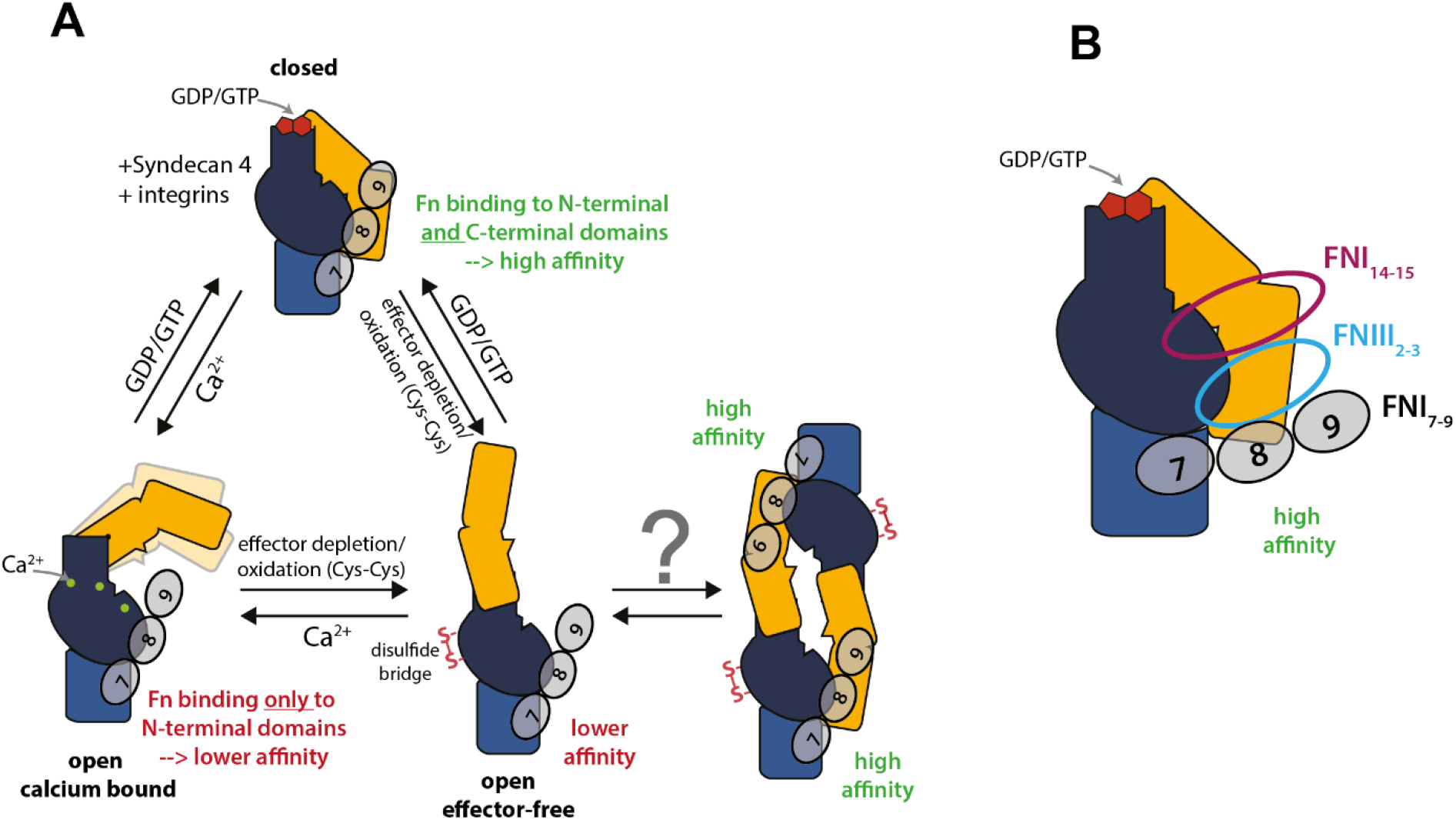
Sketch illustrating the proposed model of TG2’s mechano-regulated binding to FN-fibers and TG2’s interactions with soluble FN. **A:** TG2 interacts differently with FN-fibers depending on the conformational state of TG2. When TG2’s N-terminal β-sandwich is in spatial proximity to its C-terminal β-barrel 2, FN modules can bind both TG2 binding sites, forming a synergistic high affinity interaction. When the C-terminal β-barrels move away from the N-terminal domain due to the oxidation or calcium binding, TG2 assumes distinct open states. In this case, TG2’s C-terminal β-barrels are out of reach and FN can bind only to TG2’s N-terminal β-sandwich domain, forming a lower affinity interaction. Oxidised TG2 in the ECM (open effector-free) could form a high affinity synergy interaction with FN through dimeric TG2 association. In this case spatial proximity between N-terminal and C-terminal domains could be achieved locally, in which case the domains would be provided by two different TG2 molecules of a TG2 homodimer. Parallel alignment of FNI_7-9_ modules with TG2’s C-terminal β-barrel domains was predicted by crosslink-guided docking. **B:** Proposed interaction of TG2 with FN involving FNIII_14-15_ and FNI_2_, as revealed by XL-MS of the TG2-FN complex in solution. The C-terminal β-barrel 2 of TG2 interacts with FN modules FNIII_14-15_ and FNI_2_. At the same time, FNIII_14-15_ and FNI_2_ are in close contact with each other. TG2’s N-terminal domain interacts with FNI_7-9_ within the canonical gelatin-binding domain (GBD) of FN. Such mechano-regulated interaction mechanism that requires the presence of a synergy site might be possible between TG2 and fibrillar FN due to non-periodic arrangement of FN molecules in FN-fibers.

When discussing TG2 localization outside of the cell, two distinct pools of TG2 are typically noted: TG2 located on the cell surface and TG2 located in the ECM, which appear to perform different functions and interact with different binding partners^19^. On the cell surface, where TG2 acts as an adhesion co-receptor for FN^21^, it primarily associates with integrins and the heparan sulfate proteoglycan syndecan-4, forming ternary and/or quaternary adhesive complexes with FN^19,20,91^. Recent advances in understanding of TG2-syndecan-4 interactions suggest that this interaction stabilizes TG2 in a closed conformation, due to a composite binding site consisting of two clusters that form a single high affinity heparin-binding site when brought in proximity in a closed conformation^92,93^. This is highly relevant to our findings, since when in heterocomplex with FN, Syndecan-4 could stabilize TG2 in the closed conformation despite high (millimolar) Ca^2+^concentrations. Formation of a heterocomplex between FN-TG2 and β1-integrins, contributes to cell survival and has been implicated in several pathologies. For instance, TG2 enhances ovarian cancer cell anchoring to fibronectin, promoting metastasis^94^. In breast cancer, cooperation between TG2, integrins, and fibronectin enhances cell attachment, invasion and survival^27^. In the ECM of multiple sclerosis lesions, TG2 enhances adhesion and migration of astrocytes on fibronectin in the ECM, which contributes to glial scarring^95,96^. Noteworthy, TG2 also promotes FN-fiber deposition in multiple sclerosis lesions^97,98^ and in glioblastoma^99^. Independent studies utilized the irreversible inhibitor KCC009^100^, which targets TG2 enzymatic activity, to block TG2-mediated adhesion and migration of astrocytes on FN in multiple sclerosis^95,96^, as well as the deposition of FN matrix in glioblastoma both *in vitro* and *in vivo*^99^, and in multiple sclerosis lesions by astrocytes^97^. Although the precise mechanism of KCC009 interference with TG2-FN interactions remains unclear, this small molecule inhibitor irreversibly binds to the active site Cys277 of TG2, inactivating TG2 and trapping it in an open extended conformation^101^, similar to Z-DON inhibitor used in our study. The ability of KCC009 to disrupt TG2-mediated adhesion to FN cannot be solely attributed to the blocking of TG2 enzymatic activity, since TG2-FN adhesive interactions are independent of TG2 crosslinking activity^44^. It was furthermore discussed that TG2’s ability to cooperate with integrins to enhance deposition of FN-fibers does not require enzymatic TG2 crosslinking^24^ and that “anti-adhesive” effect of KCC009 cannot be explained by the direct blocking of the binding interface of TG2 and FN, as KCC009 does not bind directly at the TG2-FN binding interface^100^. Our findings though might suggest a mechano-regulated mechanism as KCC009 traps TG2 in the open conformation, similarly to Z-DON, disrupting TG2’s multivalent binding site and thereby reducing the affinity of TG2 for FN.

In the ECM, TG2 predominantly co-localizes with the FN matrix, although to a lesser extent, TG2 may also associate with other non-FN interaction partners, such as collagen VI^102^. Unlike on the cell surface, TG2 in the ECM is likely to adopt an open, effector-free conformation due to oxidation and the formation of a Cys370-Cys371 disulfide bridge^39^. SAXS measurements by Singh *et al.* demonstrated that constitutively open TG2 mutants form homodimers in solution^103^. The fit of TG2 in the extended conformation into the envelope suggests that the two monomers dimerize in an overlapping head-to-tail configuration. Another independent study found that homodimer formation in wild type TG2 is increased at higher temperatures^104^. The authors also used SAXS to measure the multimerization state of TG2 and similar to Singh *et al.,* found that the fitting into the SAXS envelope resulted in a head-to-tail homodimer configuration. It can be speculated that a fraction of TG2 in this dimeric state could be immobilized on FN-matrix fibrils in the ECM and be stored there in inactive state until the requirement for TG2 crosslinking arises to stabilize the matrix in case of injury (Fig. 8A). As we previously discovered that FN fibers are highly stretched under homeostasis in most healthy organs^105^, this suggests a plausible mechanism for storing inactivated oxidized TG2 dimers bound with lower affinity to stretched than to relaxed FN-matrix fibers. As FN fibers get cleaved at sites of injury, or proteolytically cleaved at sites of inflammation, the structurally relaxed FN will compete effectively for TG2 binding. We thus propose here that this tension dependent switch might allow TG2 storage in ECM of healthy organs, and for fast recruitment of TG2 to damaged FN fibers in case of injury. Supporting this idea, it has been shown that TG2 bound to FN-matrix can be reduced by the enzymatic activity of Thioredoxin and recruited in this way for the crosslinking activity of matrix proteins^73^. Future research should investigate whether TG2 homodimer can bind fibrillar FN and if such binding can be regulated by FN-fiber strain.

Therefore, our results indicate that, like for Syndecan-4, the FN-binding epitope on TG2 is conformational. Interestingly, studies have identified anti-TG2 autoantibodies that induce a shift in a pool of effector-free TG2 either toward the “closed” or “open” conformation, depending on the epitope on TG2 they bind to^40,106^. Although the exact reasons are not known, this could be a common regulatory mechanism shared by many TG2 binding partners to exert an effect on TG2 conformation and, hence, its function. Our findings shed light on the role of TG2-FN in cell-ECM interactions in various physiological processes. As we recently demonstrated that FN fibers are under low tension in cancer tissue and virally infected lymph nodes, our findings that the interaction of TG2 with FN is tuned by its fiber tension is highly significant. We now suggest that TG2 could serve as a FN-fiber strain sensor modulating cell-ECM interactions, such as cell adhesion, cell spreading and cell migration depending on the tensional state of ECM fibers, thereby modulating a tissue response in wound healing and homeostasis.

## Materials and Methods

### Reagents

Reagents were purchased from Sigma Aldrich, if not mentioned otherwise. Guinea pig liver transglutaminase (gpTG2), Zedira, T006; human tissue transglutaminase (hrTG2, recombinantly produced in *E.coli*), Zedira, T002; short human tissue transglutaminase, aa 1-465 (short hrTG2, barrel 1and 2 deletion mutant, recombinantly produced in *E.coli*), Zedira (T167); inactive human tissue transglutaminase (Cys277Ser mutant, recombinantly produced in *E.coli*), Zedira (T018); Z-DON-Val-Pro-Leu-OMe, Zedira, (Z006); R1R2 peptide^66,107^ (amino acid sequence: GLNGENQKEPEQGERGEAG-PPLSGLSGNNQGRPSLPGLNGENQKEPEQGERGEAGPP) was manufactured by GenScript ; Alexa Fluor 647 NHS Ester (A20006), ThermoFisher; Alexa Fluor 488 NHS Ester (A20000), ThermoFisher; Alexa Fluor 546 C5 Maleimide (A10258), ThermoFisher; fibronectin proteolytic fragment from human plasma collagen/gelatin-binding domain (GBD), 45kDa, F0162, Sigma Aldrich; anti-fibronectin antibody, gelatin binding domain, close IST-10, from mouse 1 mg/ml (MAB1892), Sigma Aldrich; secondary pAB (ab150107) DK anti-mouse (to MS IgG) (2mg/ml), Abcam; guanosine 5’-diphosphate disodium salt (GDP), G7127 Sigma Aldrich; silicone sheeting, .010’’ NRV G/G 40D 12’’x12’’, SMI specialty manufacturing, inc.; Slide-A-Lyser Dialysis Casette, 20,000MWCO, 66005 ThermoScientific; centrifugal filter units Amicon Ultra – 0.5ml, 30K UFC503096 Merck Millipore; disposable PD-10 desalting columns, GE17-0851-01 Merck Millipore; Amicon Ultra 0.5 ml Ultracel 30k Centrifugal filter units, Merk, Millipore; L-Glutathione oxidised (GSSG), G4376-5G, Lot#SLCJ0220, MW: 612.63g/mol.

### FN isolation from human plasma

Fibronectin (FN) was isolated from human plasma according to a previously described protocol^50^. Briefly, 2 mM phenylmethylsulphonyl fluoride (PMSF) and 10 mM ethylenediaminetetraacetic acid (EDTA) were added to human plasma (Zürcher Blutspendedienst SRK) and centrifuged at 15,000 g for 40 min. Next, plasma was passed through a size-exclusion column (PD-10 Desalting columns, GE Healthcare) and loaded onto a gelatin-sepharose 4B column (VWR Schweiz). After washing with phosphate buffered saline (PBS+10mM EDTA), NaCl (1 M in PBS) and arginine (0.2 M in PBS), FN was eluted with 1.5 M arginine in PBS. For FRET labelling of FN with donors and acceptors, the gelatin-sepharose 4B column was washed with 1M NaCl, 1M urea, and was eluted from the column with 6 M urea. FN purity was checked by SDS-PAGE and western blotting (data not shown). The purified FN was aliquoted and stored in 1.5 M arginine in PBS at -80°C.

### Fluorescence labelling of FN and transglutaminases

FN and TG2 were labelled with ALEXA fluorophores (AF) on primary amines with AF488-NHS-ester (for FN-fiber stretch assay with calcium titration) and AF674-NHS-ester respectively, to obtain FN-488 and TG2-674 according to the manufacturer’s instructions (ThermoFisher). Briefly, the buffer was exchanged to carbonate-bicarbonate labelling buffer (0.2 M NaHCO_3_ – Na_2_CO3, pH 8.5) using Slide-A-Lyser dialysis cassettes. Next, FN was incubated with 40-fold molar excess of AF488-NHS-ester, and transglutaminases with 45-fold molar excess of AF647-NHS-ester for 1 h at RT on rotator. Free dye was removed by buffer exchange with 50 mM Tris, pH 7.4, using Slide-A-Lyser dialysis cassettes. The protein purity, concentration and degree of labelling were assessed by spectrophotometric measurements of absorbances at 280 nm, 488 nm, and 647 nm. Proteins were aliquoted, snap frozen in liquid nitrogen and stored at -20°C.

### FN labelling with donors and acceptors for FRET

FN was labelled for FRET as previously described^50,71^. Briefly, FN-dimer was doubly labelled on primary amines with AF488 (donor), and with AF546 (acceptor) on the four free cryptic cysteines in modules FNIII_7_ and FNIII_15_. Briefly, FN was denatured in 6 M Urea by addition of equal volume of 8 M guanidine hydrochloride and incubated for 1 h at RT with 20-fold molar excess of AF546-maleimide. Free dye was removed with buffer exchange to carbonate-bicarbonate buffer (0.2 M NaHCO_3_ – Na_2_CO3, pH 8.5) using size-exclusion chromatography (PD-10 Desalting columns, GE Healthcare). Next, FN in carbonate-bicarbonate buffer was immediately incubated on rotator with a 40-fold molar excess of AF488-NHS-ester for 1 h at room temperature. Afterwards, free dye was removed by buffer exchange to PBS, pH 7.4 using PD-10 Desalting columns. The protein purity, concentration and degree of labelling were assessed by spectrophotometric measurements of absorbances at 280 nm, 488 nm, and 546 nm. FN-FRET was aliquoted, snap-frozen in liquid nitrogen and stored at -80°C. To construct FN-FRET denaturation curve in order to assess the correlation of the FRET signal to the loss of FN tertiary and quaternary structure in solution, FN-FRET was chemically denatured with increasing concentrations of guanidine hydrochloride as was previously described^50,71^. Briefly, the coverslips were blocked with 4% bovine serum albumin (BSA) for 1 h at RT, then washed with PBS. FN-FRET was mixed with an equal volume of guanidine hydrochloride in PBS to a final concentration of 0 M, 0.5 M, 1 M, 2 M, 3 M, and 4 M and incubated for 1 h at RT. To separate FN dimer to monomers, disulfide bonds between two FN monomers were reduced with 50 mM DTT for 1 h at RT, before incubating FN-FRET with 1 M guanidine hydrochloride. A droplet from each solution was added onto the prepared coverslip and imaged using Olympus FV1000 confocal microscope as described below.

### FN fiber stretch assay

FN fiber stretch assay was performed as was previously described^58,62^. FN fibers were manually pulled from a droplet of 0.4 mg/ml FN solution (to avoid inter-molecular FRET, not more than 10% FN-FRET probes were added) with the needle tip and deposited on the silicone sheet mounted on the custom-made uniaxial stretch device. For the single fiber studies, 15 fibers were deposited parallel to the stretch axis. To create intersecting fibers, 10 fibers were deposited both parallel and perpendicular to the stretch axis (Fig. 2A). Next, FN-fibers were washed with PBS, and silicone sheets were stretched, relaxed, or left unchanged (native), subjecting the silicone membranes to strains of 100%, -50% or 0% respectively. Previous calibration studies converted the silicone membrane strains to the respective FN-fiber strains of 380% (stretched membrane: high FN-fiber strain), 20% (relaxed membrane: low FN-fiber strain) and 140% (native membrane: medium FN-fiber strain) parallel to the stretch axis and 90% (stretched membrane), 219% (relaxed membrane) and 140% (native membrane) perpendicular to the stretch axis^70^. Though the native membrane is not subjected to any strain, FN-fibers are typically pre-strained to ∼140% due to the forces required to pull them out of the droplet^70^. Next, FN-fibers were blocked with 4% BSA for 30 min at RT, then washed with PBS. Transglutaminases were induced into a specific conformation by 30 min pre-incubation with corresponding allosteric effectors, immediately added to FN-fibers at a concentration of 10 µg/ml TG2-647 in 50mM Tris, and incubated for 1h at RT. TG2 solutions were added to FN-fibers and incubated on fibers for 1 h at RT. This was followed by 3 times washes for 5 min each with 50 mM Tris and fixation with 4% PFA in PBS for 15 min at RT. FN-fibers were imaged immediately using Olympus FV1000 confocal microscope.

### Induction of conformational changes of TG2 with its allosteric effectors

To induce the closed conformation (TG_GDP_), 10 µg/ml TG2-647 was pre-incubated for 30 min at RT in 1 mM GDP, 1 mM EDTA in 50 mM Tris. To induce catalytically active open conformation, TG2 was pre-incubated for 20 min at RT in 1.2 mM CaCl_2_ in 50 mM Tris. To induce catalytically inactive extended conformation (effector-free on Fig. 2A), TG2 was pre-incubated with 100 µM “Z-DON” Z006 (Zedira) in 5 mM CaCl_2_ in 50 mM Tris for 30 min at RT. To get rid of free Z006, the solution was passed through centrifugal filter units (Ultracel 30k, Merk, Millipore), according to manufacturer’s instructions, and TG2 bound to Z006 “Z-DON” (TG2_Z006_) was resuspended in 50 mM Tris or 1.2 mM CaCl_2_ in 50 mM Tris. To oxidise TG2 (oxTG2), guinea pig liver TG2 (gpTG2) was incubated in 2 mM GSSG and 1 mM EDTA in 50 mM Tris for 3 h at 37°C. oxTG2 was passed through centrifugal filter units and resuspended in 50 mM Tris.

### A dose-response competition experiment with R1R2 peptide

A dose-response competition study with R1R2 peptide and a constant concentration of TG2_GDP_ (10 µg/ml gpTG2) was performed using only horizontally pulled FN-FRET fibers (parallel to the stretch axis) on relaxed membranes (low FN-fiber strain). After blocking FN-fibers with BSA, 10 µg/ml TG2_GDP_ labelled with AF647 was added to the fibers together with R1R2 peptide at one of the concentrations: 0 µM, 17 µM, 30 µM, 50 µM, 80 µM and 100 µM. TG2_GDP_ and R1R2 were diluted in 1 mM GDP, 1mM EDTA, 1mM MgCl_2_ in 50 mM Tris. Solutions were incubated on FN-fibers for 1 h at RT, FN-fibers were washed 3 times for 5 min with 50 mM Tris and fixed with 4% PFA in PBS for 15 min at RT.

### Calcium dose-response experiment

For calcium dose-response experiments, FN labelled only with AF488 was used and guinea pig liver (gpTG2) was labelled with AF-647. Like R1R2 competition studies, only horizontally pulled FN-AF488 fibers (parallel to the stretch axis) on relaxed membranes (low FN-fiber strain) were used for this experiment. 10 µg/ml gpTG2 was pre-incubated for 20 min at RT with 1 µM, 100 µM, 250 µM, 500 µM, 1.2 mM, 5 mM, and 10 mM CaCl_2_ in 50 mM Tris. 10 µg/ml gpTG2 pre-incubated in 1 mM GDP, 1 mM EDTA in 50 mM Tris and 10 µg/ml gpTG2 pre-incubated in 1 mM EDTA in 50 mM Tris were used as controls. gpTG2 solutions were incubated on the fibers for 1 h at room temperature, followed by 3 times for 5 min washes and fixation with 4% PFA in PBS. z-stacks of 3 focal planes with steps of 1micron were obtained with Nikon Visitron Spinning disk confocal, 60x water immersion objective, using 488 and 640 lasers and confGFP and confCy5 emission filter cubes. Images were analysed with custom-written Python script. Briefly, Gaussian filter (sigma=1) was applied to both FN and TG2 channels, FN channel was thresholded using Otsu algorithm and binary mask created, the mask was applied to both the FN and TG2 channel, replacing all background pixels with NaN, which were excluded from analysis. Then, TG2 signal was normalized pixel-by-pixel to the FN signal (TG2-647/FN-488), and the mean normalized TG2 intensity was calculated over the maximum intensity projection of the normalized TG2 signal.

### FRET confocal microscopy of FN-fibers with bound TG-647

Z-stacks parameters were set so that the acquisition of stacks of three images always started from the top surface of an FN-fiber, moving inside the fiber interior (toward silicone membrane), first by 1 and then by 2 µm. Z-stack images were acquired with Olympus FV1000 confocal microscope with 0.9NA 40X water immersion objective, according to a procedure described in detail previously^62^. Briefly, image acquisition was performed in three sequential steps with five channels using excitation and detection windows as depicted in Supplementary Fig. 1. Donor, acceptor, and FRET intensities were measured with 12 nm bandwidths over acceptor and donor emission peaks. TG2-647 excitation and detection were always performed in a separate channel (channel 5), to avoid unwanted crosstalk from the donor AF488 and acceptor AF546 signals upon excitation with 488 nm and 543 nm lasers. FRET-signal correction due to the bleed-through of both donor and acceptor signals into the FRET-excited acceptor channel was performed as was described in detail previously^62,108^, and an example of the calculation of β- and γ-correction factors is shown in Supplementary Fig.1. Acquisition parameters (laser intensity and PMT voltage) were set to maximize the detected signal, while minimizing bleaching and background signal, and were kept constant throughout the experiment.

### Ratio-metric FRET image analysis and a correlation of TG2-647 binding with FN-FRET ratio

Ratio-metric FRET signal calculation and a correlation of TG2-647 signal to the FN-FRET ratios was performed pixel-by-pixel using custom written Matlab script, which was validated and described in detail before^62^. Briefly, images from different channels were aligned using “imregister” Matlab function, thresholded, mean background signal subtracted, and a mask generated. To remove the pixels at the edges of fibers, where light scattering can create false FRET values, edges of the mask were eroded, then a local 3x3 averaging filter and a one-deviation Gaussian averaging filer were applied. FRET ratios were calculated by dividing pixel-by-pixel the corrected intensity from the acceptor emission due to FRET (channel 1 in Supplementary Fig.1) by the donor emission (channel 2 in Supplementary Fig.1). Since the intensity of TG2 signal is dependent on the amount of FN to which it binds, we normalized the TG2-647 signal (channel 5 in Supplementary Fig.1) by dividing it pixel-by-pixel by the directly excited acceptor AF546 (channel 3 in Supplementary Fig.1): TG2-647/FN-546. Examples of raw confocal images and corresponding color-coded pixel-by-pixel result after the image processing are shown in Supplementary Fig. 3 (intersecting fibers) and Supplementary Fig. 5 (horizontal fibers). To plot the mean, maximum intensity projections (MIP) of obtained three-dimensional arrays were taken and the mean intensity of FN-FRET ratio and mean intensity of TG2-647 signal were calculated from MIP and plotted as mean scatterplots. To relate each pixel FN-FRET value with its corresponding TG2 pixel intensity, they were plotted versus each other as two-dimensional density plots (Supplementary Fig. 4).

### Statistical analysis for FN-fiber stretch assays

Since not all data had a normal distribution, statistical significance between groups was assessed by the nonparametric two-sided Wilcoxon rank sum statistic for two samples, which does not imply data distribution normality, as was performed previously in a similar study^62^. Statistical significance was assessed for *0.01≤ p <0.05; **0.001≤ p <0.01; ***10^-5^≤ p <0.001; **** 10^-6^≤ p <10^-5^; ***** p <10^-6^

### Microplate FN-TG2 binding assay

To determine the concentrations of full-length FN and with 45kDa-FN at which microtiter plates would be coated with approximately similar number of binding sites for TG2, titration experiments were carried out. For this, 96-well black flat non-transparent bottom microtiter plates were coated with 0.1, 0.2, 0.4, 0.6, 0.8, 1, 2, 5, 10, 20, 30 and 50 μg/ml full-length FN or 45 kDa-FN GBD in PBS at 4°C overnight. Wells were washed with PBS 3 times for 5 min and blocked for 30 min with 4% BSA in PBS, followed by another wash. Full-length FN and 45 kDa-FN GBD were detected with monoclonal mouse anti-FN antibody with epitope within GBD (MAB1892, Sigma), clone IST-10 (1:1000 in 2% BSA in PBS) for 1 h at RT, followed by incubation with anti-mouse pAB ab150107 AF647 (1:1000 in 2% BSA in PBS) for another 1 h at RT, and finally fixation with 4% PFA for 10 min. After each antibody incubation wells were washed with PBS 3 times for 5 min. Fluorescence intensity was measured with Tecan Spark plate reader. Obtained data were fit using a four-parameter logistic (4PL) curve. Next, the concentration of the full-length FN at which 95% of maximum intensity was reached were calculated from the curve as 2.4 μg/ml. Then, the concentration of 45 kDa-FN GBD, which corresponded to the 95% of full-length FN intensity was calculated from the curve as 0.6 μg/ml. In all subsequent experiments, microtiter plates were coated with 2.4 μg/ml of full-length FN and 0.6 μg/ml of 45 kDa-FN GBD.

For experiments with wild type gpTG2 and short hrTG2, 96-well black flat non-transparent bottom microtiter plates were coated with either 2.4 μg/ml full-length FN or 0.6 μg/ml 45 kDa-FN at 4°C overnight. After incubation, wells were washed 3 times for 5 min with 50 mM Tris and blocked for 30 min with 4% BSA in 50 mM Tris at RT. TG2-647 solutions of 0.1, 0.5, 1, 2, 3, 5, 7, 10 μg/ml were prepared in either 1mM GDP, 1 mM EDTA in 50 mM Tris, or 1 mM EDTA in 50 mM Tris, or 1.2 mM CaCl_2_ in 50 mM Tris, or 1.2 mM CaCl_2_, 100 μM Z006 in 50 mM Tris and pre-incubated for 20 min at RT. Transglutaminase dilutions were added to the FN-coated wells and incubated at RT for 1 h. Wells were washed 3 times for 5 min with 50 mM Tris and fixed with 4% PFA for 10 min. Fluorescence intensity was measured with Tecan Spark plate reader. Obtained data were fit using a four-parameter logistic (4PL) curve and plotted.

### Chemical cross-linking

TG2 was incubated either with either full-length FN, or the 45-kDa FN fragment in 1 mM GDP, 1 mM EDTA in PBS for 1 hr at room temperature. Cross-linking experiments were performed with 50 µg total protein at concentrations of approximately 0.7 mg/ml for samples containing full-length fibronectin and at approximately 0.25 mg/ml for samples containing the 45-kDa FN (GBD). All steps were performed on a ThermoMixer (Eppendorf) at 750 rpm.

Lysine cross-linking with a 1:1 mixture of non-deuterated and deuterated disuccinimidyl suberate (DSS-d_0_/d_12_, Creative Molecules) was initiated by adding the cross-linking reagent from a freshly prepared 25 mM stock solution in dimethyl formamide to a final concentration of 1 mM^109^. The reaction was allowed to proceed for 60 min at room temperature (23 °C) before adding ammonium bicarbonate to a final concentration of 50 mM and incubating further for 30 min.

Cross-linking of carboxyl groups and of amine groups with carboxyl groups was performed using non-deuterated and deuterated pimelic dihydrazide (PDH-d_0_, ABCR, and PDH-d_10_, Sigma-Aldrich) and the coupling reagent 4-(4,6-dimethoxy-1,3,5-triazin-2-yl)-4-methylmorpholinium (DMTMM) chloride (Sigma-Aldrich). Stock solutions were prepared in phosphate-buffered saline at concentrations of 140 mM (PDH, 1:1 mixture of d_0_ and d_10_) and 520 mM (DMTMM). Two different reaction conditions were used for combined PDH/DMTMM cross-linking: 44 mM PDH and 88 mM DMTMM or 22 mM PDH and 11 mM DMTMM, respectively^110^. Samples were incubated for 60 min at room temperature and the reaction was stopped by passing the sample solutions through a Zeba spin desalting column (ThermoFisher Scientific) according to the manufacturer’s instructions. Quenched samples solutions were subsequently evaporated to dryness in a vacuum centrifuge.

### Sample processing and mass spectrometric analysis

Dried samples were dissolved in 50 µL of 8 M urea solution. Cysteine thiols were reduced by addition of tris(2-carboxyethyl)phosphine to a final concentration of 2.5 mM and incubation for 30 min at 37 °C, and free thiol groups were alkylated with iodoacetamide (final concentration of 5 mM, incubation for 30 min at room temperature, protected from light) prior to enzymatic digestion.

Proteolysis with endoproteinase Lys-C and trypsin was performed as follows: The reduced and alkylated samples were diluted to 5.5 M urea with 150 mM ammonium bicarbonate solution and Lys-C (Wako) was added at an enzyme-to-substrate ratio of 1:100. After incubation at 37 °C for 2 h, the solutions were further diluted to 1 M urea with 50 mM ammonium bicarbonate, and trypsin (Promega) was added at an enzyme-to-substrate ratio of 1:50. Trypsin digestion was then allowed to proceed overnight.

Proteolysis with chymotrypsin (Roche) was performed by diluting the solutions to 1 M urea with 50 mM ammonium bicarbonate and adding the protease at an enzyme-to-substrate ratio of 1:50. Samples were incubated for 2 h at 25 °C.

Enzymatic digestion reactions were stopped by adding 100% formic acid to a final concentration of 2% (v/v) and samples were purified by solid-phase extraction using 50 mg SepPak tC18 cartridges (Waters). Purified samples were fractionated by peptide size exclusion chromatography (SEC) on an Äkta micro FPLC system equipped with a Superdex 30 Increase column (300 × 3.2 mm, GE/Cytiva). The separation was carried out with a mobile phase consisting of water/acetonitrile/trifluoroacetic acid (30/70/0.1, v/v/v) at a flow rate of 50 µL/min. For each sample, three or four 100 µL fractions from the high mass region were individually collected and evaporated to dryness.

Liquid chromatography-tandem mass spectrometry was performed on an Easy nLC-1200 HPLC system coupled to an Orbitrap Fusion Lumos mass spectrometer equipped with a Nanoflex electrospray ionization source (all ThermoFisher Scientific). Approximately one µg or less for each SEC fraction was injected and the peptides were separated on an Acclaim PepMap RSLC column (150 mm × 75 µm, 2 µm particle size, ThermoFisher Scientific) at a flow rate of 300 nl/min. The mobile phase gradient was set to 11% B to 40% B in 60 min, with mobile phase A = water/acetonitrile/formic acid (98/2/0.15, v/v/v) and B = acetonitrile/water/formic acid (80/20/0.15, v/v/v). The mass spectrometer was operated in the data-dependent, “top speed” acquisition mode with a cycle time of 3 s. Precursor ion spectra were acquired in the orbitrap analyzer at a resolution setting of 120000. Precursors with a charge state between +3 and +7 were selected for collision-induced fragmentation in the linear ion trap with a normalized collision energy of 35%. For most experiments, fragment ions were detected in the orbitrap analyzer at a resolution setting of 30000. For some samples, one of the two replicate injections was performed with MS/MS detection in the linear ion trap to achieve higher sensitivity. In that case, fragmentation conditions remained unchanged, and the linear ion trap was operated with a “rapid” scan rate.

### Identification of cross-linked peptides

Preliminary experiments found unequivocal evidence for UniProt proteoforms P02751-1 and -9, with isoform 1 having the most proteotypic peptides. This isoform was therefore included in the sequence database and all numbering refers to it. In addition, the database consisted of the sequence of transglutaminase and the sequence of three contaminants identified from E. coli: CH60, SLYD, and ARNA. Target/decoy searches were performed using a decoy database containing the reversed sequences of all database entries.

MS/MS spectra were searched against these databases using xQuest, version 2.1.5 (available from https://gitlab.ethz.ch/leitner_lab)^111^. The cross-linking specificities were set as follows: For DSS, K and the protein N terminus; for PDH, D and E; for DMTMM, K with D and K with E. For Lys-C/trypsin digestion, protease specificity was set to cleavage after K and R unless followed by P, with a maximum of two missed cleavages per peptide allowed (excluding the cross-linking site, if applicable). For chymotrypsin, cleavage was considered after F, L, M, W, and Y, with a maximum number of missed cleavages set to four. The mass error was set to ±15 ppm for orbitrap detection and to 0.2 Da and 0.3 Da for the detection of “common” and “xlink” fragment ion types for detection in the linear ion trap, respectively. All identifications were further filtered for a %TIC value ≥0.1 and a main xQuest score of ≥20 for fragment ion detection in the ion trap and ≥25 for fragment ion detection in the orbitrap analyzer, respectively. MS/MS spectra of the remaining candidates were evaluated and rejected if they did not contain at least four bond cleavages in total per peptide or at least three consecutive bond cleavages per peptide.

### Integrative structural modeling and docking

A detailed description of the entire modeling process, which allows for replication of the results is provided in Supplementary Notes 1-3, overview of the modeling workflow is provided in Supplementary Fig. 13. Briefly, I-TASSER^83^ was used for structural refinement of available PDB templates of TG2, and FN fragments using experimental crosslinks within those regions as distance restraints (Supplementary Note 1). ROBETTA^85^ and AlphaFold^86^ were used for modeling of FN_7-9_ and 45-kDa FN (GBD), followed by evaluation of resulting structures using QMEAN^87^. Intra-protein crosslinks were used both as distance restraints for structural refinement and for evaluation of resulting models based on their compatibility/non-compatibility with the imposed distance cutoff. To assign compatibility/non-compatibility criteria, Euclidean distances between β-carbons (CB-CB) of the crosslinked residues were calculated using Xwalk^84^. Euclidean distance cut-offs we set as follows: for DSS < 35 Å, ED for DMTMM < 25 Å, ED for PDH < 35Å, and crosslinks above the set cut-off value were considered as incompatible. Inter-protein crosslinks were validated and active residues predicted using DisVis^65^ (Supplementary Note 2). To predict active residues involved in the interaction, we submitted to DisVis server residues with relative solvent accessibility of at least 40% for either the main chain or the side chain. The relative solvent accessibility was calculated using NACCESS^88^. Inter-protein crosslinks that were predicted by DisVis to be incompatible were excluded from further docking steps. Docking was performed with HADDOCK^64^. Detailed description of each docking run, and all used distance restraints are provided in Supplementary Note 3. Experimental inter-protein crosslinks were mapped onto predicted models and Euclidean distances calculated using Xwalk. Obtained structures were visualized with ChimeraX^112^.

### Data availability statement

The mass spectrometry proteomics data have been deposited to the ProteomeXchange Consortium^68^ via the PRIDE^113^ partner repository with the dataset identifier PXD043976.

## Significance and conclusions

We discovered here that the binding of crosslinking enzyme to ECM fibers is mechano-regulated. Mechanical forces acting on ECM fibers can regulate the interactions of TG2 with FN and we suggest a novel mechanism of this mechano-regulated TG2-FN interactions based on combining FN-fiber stretch assay with XL-MS studies to decipher a structural mechanism. A previously unknown role of C-terminal β-barrel domains of TG2 is proposed, i.e. that they serve as a synergy binding site for FN, and are essential for tuning the mechano-regulated binding. In contrast to what was previously assumed, we show that TG2’s conformational states affect TG2 interactions with FN, since mechano-regulated binding is only possible when N-terminal and C-terminal domains of TG2 are in spatial proximity. Thus, our study improves the understanding of TG2-FN binding interface. This knowledge will also help in the future design of interfering compounds targeting TG2-FN interactions. We thus propose a new role for TG2 as an FN-fiber strain sensor and a modulator of cell-ECM interactions in response to the ECM tensional state. Our model also offers a new perspective into how mechanical forces acting on ECM fibers can reciprocally regulate critical cellular processes of adhesion, spreading, and migration through the mechano-regulation of crosslinking enzymes binding with the ECM.

## Contributions

KS, LB, VV conceived the study and designed the research. All FN-fiber stretch experiments and microplate protein-binding experiments were performed by KS. Analysis of all confocal images and of plate reader data and data visualization were performed by KS. LB wrote the maximum intensity projection script for Python. FLB under supervision of AL performed all crosslinking experiments and data analysis. KS visualized XL-MS data performed integrative structural modeling and docking with input from AL. PP performed calculation of solvent accessibilities with NACCESS and contributed ideas to the discussion. SP added insights. KS, LB, AL and VV wrote the manuscript. All authors read and edited the manuscript. Research was funded by grants to SP and VV.

## Acknowledgements

The authors thank Chantel Spencer for isolating plasma fibronectin, Dr. Mario Benn for discussions and general support, Zhe Lin for involvement in early preliminary experiment, Dr. Szymon Stoma (ScopeM) for checking the Python script which outputs normalized pixel-by-pixel FN-488/TG2-647 mean intensity values, Scientific Center for Optical and Electron Microscopy (ScopeM) of ETH Zurich, Joachim Hehl and Tobias Schwartz from ScopeM for support with Nikon Spinning Disk imaging. AL acknowledges Paola Picotti for access to instrumentation and laboratory infrastructure. The authors are grateful to Prof. Ludvig Sollid and Dr. Jorunn Stamnaes at the University of Oslo for discussions on their previous TG2-FN studies.

FLB was supported by the Swiss-European Mobility Program (SEMP). This study was funded in part by the Velux Stiftung Foundation Project Nr. 1227 (SP), as well as the ETH Zurich (VV).

