## Supplementary Information for "Tissue Transglutaminase 2 has higher affinity for relaxed than for stretched fibronectin fibers"

**Table S1: FN-FRET and TG2-AF647 image acquisition parameters**

| Phase | Channel | Laser excitation (nm) | Detection range (nm) | Description |
|---|---|---|---|---|
| **1** | 1 | 488 | 566-578 | FRET excited acceptor |
| | 2 | - | 514-526 | Donor FN488 |
| **2** | 3 | 543 | 566-578 | Direct acceptor excitation FN546 |
| | 4 | - | 514-526 | Bleed through |
| **3** | 5 | 633 | 645-745 | TG2-AF647 |

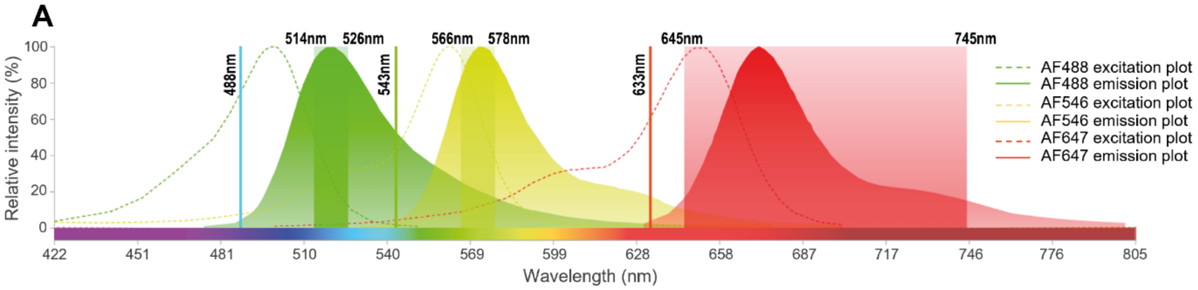


Calculation of correction factors ß and γ for estimation bleed through

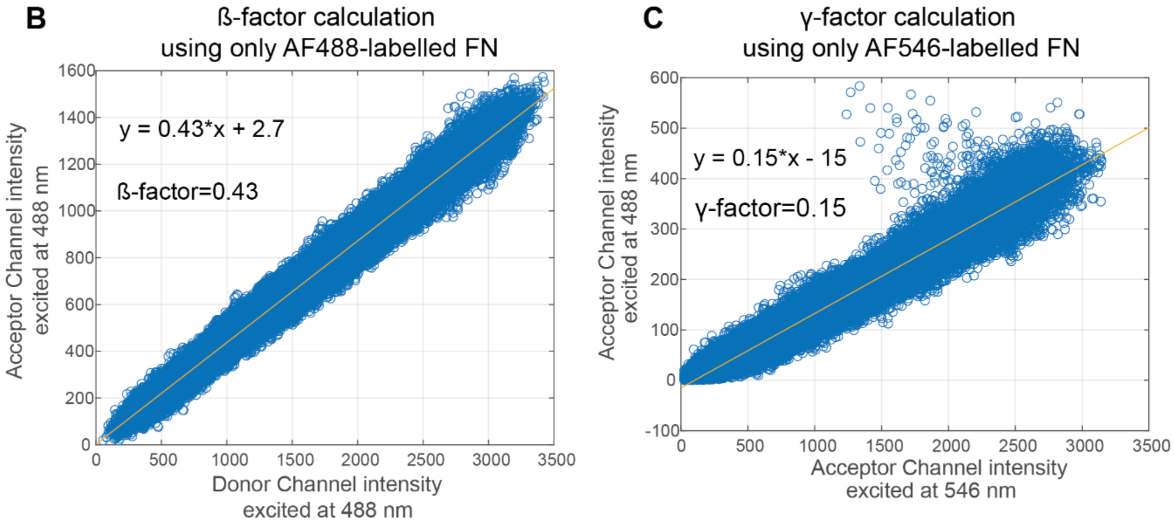

**Supplementary Figure 1: FN-FRET and TG2-647 image acquisition and correction parameters.** Table S1 summarizes image acquisition parameters, which were used for all FN-FRET and TG2-647 confocal imaging experiments. Image acquisition was performed with Olympus FV-1000 in 3 sequential steps with 5 channels as was previously described[1]. (A) Plots depicting excitation and emission spectra for the AF488 (donor), AF546 (acceptor) and 647 (TG2) fluorophores are shown. Excitation wavelengths and detection windows of each channel in table S1 are marked as well. Donor, acceptor, and FRET intensities were measured with 12nm bandwidths over acceptor and donor emission peaks. TG2-647 excitation and detection was always performed in a separate channel (channel 5), to avoid unwanted crosstalk from the donor FN488 and acceptor FN546 signals upon excitation with 488nm and 543nm lasers. The plot shows an overlap between the AF-488 and AF-546 spectra, which upon excitation with 488nm laser would result in the leak of AF-488 into the "FRET excited acceptor" channel 1 and, therefore, requires correction. (B and C) As was experimentally shown in the past, upon excitation with 488nm laser, there is a bleed through of both AF-488 (donor) and AF-546 (acceptor) signals into the FRET-excited acceptor channel (channel 1)[1]. Since this will influence the FRET ratios ($I_A/I_D$), the correction of the FRET signal is required. An example for calculation of β-factor and γ-factors for the correction of the FRET signal is provided in (B) and (C) respectively, and was performed as was described in detail previously[1,2]. β-factor and γ-factors for correction were always re-calculated for each experiment whenever the laser intensities or detection voltages were adjusted for the image acquisition.

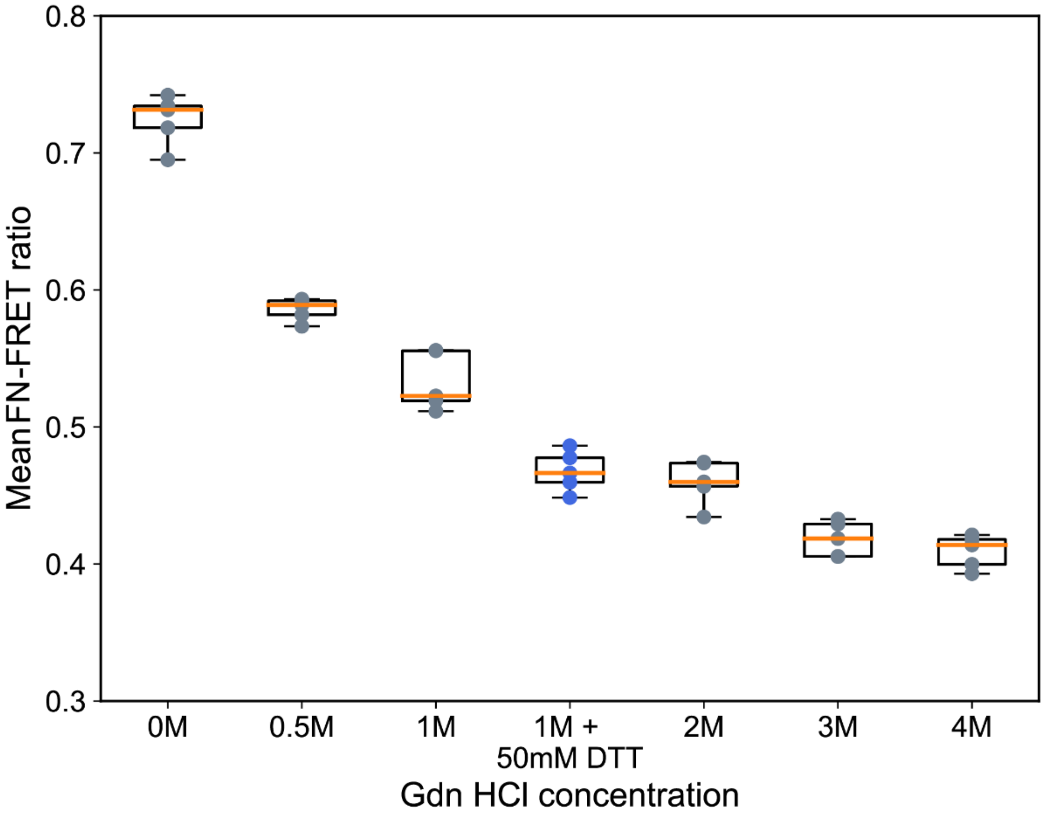


**Supplementary Figure 2: FN-FRET denaturation with guanidine hydrochloride in solution.** Increasing concentrations of guanidine hydrochloride (GdnHCl) lead to loss of FN tertiary and then secondary structure in solution upon chemical denaturation, which is in turn reflected in progressively lower FN-FRET ratio ($I_A/I_D$), confirming the responsiveness of our FN-FRET probe to a range of FN conformations, as was described before [3,4]. Treatment of FN-FRET with 50 mM dithiothreitol (DTT) for 1 h before mixing with 1mM GdnHCl (blue circles) reduced structurally important intra-module and inter-subunit disulfide bonds of FN separating FN-homodimer into monomers, which caused further reduction in FRET ratio, as opposed to the treatment with only 1M GdnHCl, as was previously described[5].

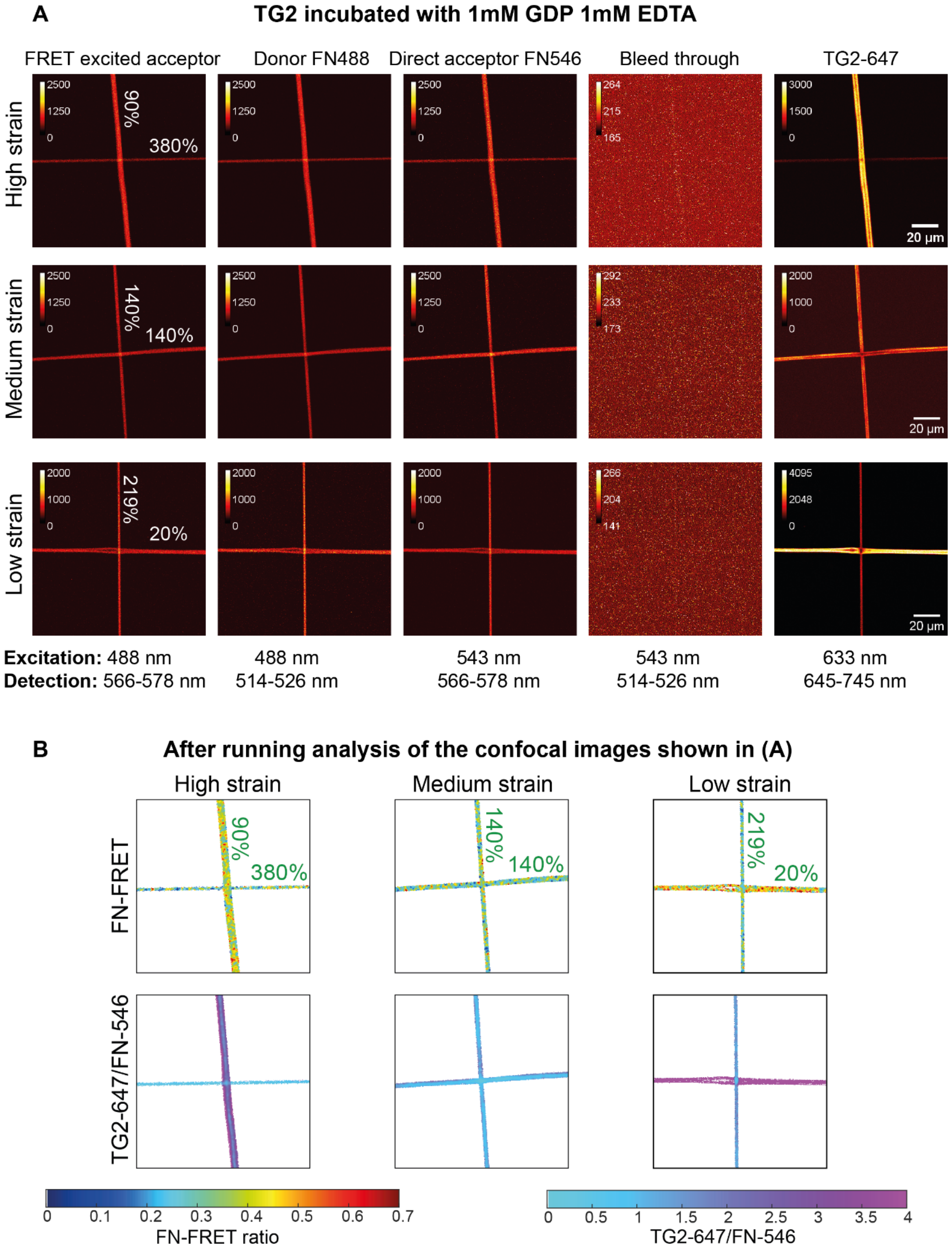

A
TG2 incubated with 1mM GDP 1mM EDTA
FRET excited acceptor
Donor FN488
Direct acceptor FN546
Bleed through
TG2-647
High strain
Medium strain
Low strain
90%
380%
140%
140%
219%
20%
20 µm
Excitation: 488 nm
Detection: 566-578 nm
488 nm
514-526 nm
543 nm
566-578 nm
543 nm
514-526 nm
633 nm
645-745 nm
B
After running analysis of the confocal images shown in (A)
High strain
Medium strain
Low strain
FN-FRET
TG2-647/FN-546
FN-FRET ratio
TG2-647/FN-546

**Supplementary Figure 3: TG2-647 binding to FN-FRET fibers deposited as intersections in the presence of 1mM GDP and 1mM EDTA, imaging, and image analysis illustration.** (A) Confocal images of a representative intersection for each membrane strain (low strain (relaxed membrane), medium strain (native membrane), high strain (stretched membrane)) are shown. Images were acquired as described in Supplementary figure 1. FN-fiber strains induced by the silicone membrane strain are indicated, as was previously reported[6]. (B) Confocal images shown in (A) were analyzed with the custom written Matlab script, which allows pixel-by-pixel correlation of normalized TG2 binding to the FN tensional state assessed by pixel-to-pixel FN-FRET ratio measurements[1]. Lower FN-FRET ratios correspond to a higher FN tensional state, and higher FN-FRET ratios corresponds to lower FN-FRET tensional states. Pixels of FN-intersections were color coded based on their corresponding FN-FRET ratio values. FN-fibers under a specific strain, display a distinct range of heterogeneous FN-conformations as was previously observed in the ECM fibers assembled by fibroblasts[3,7,8]. TG2 fluorescence intensities were normalized pixel-by-pixel to the directly excited FN-546 (TG2-647/FN-546), and these pixels were also color coded based on the TG2-647/FN-546 value. TG2 preferentially binds FN-fibers under low strain. Multiple intersections for each membrane strain were probed and analyzed in the same manner (Supplementary figure 4).

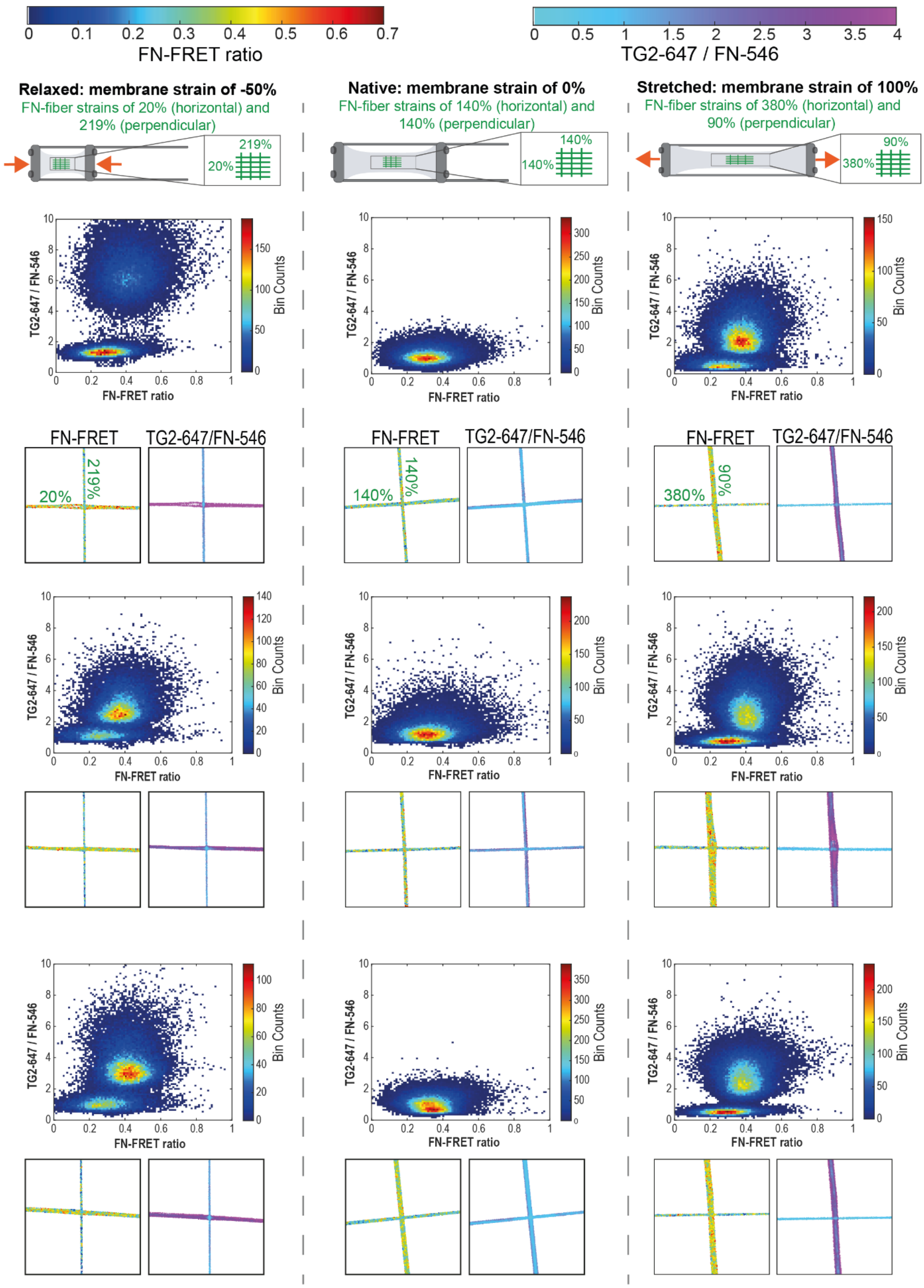


**Supplementary Figure 4: FN-fiber stretch assay performed with WT gpTG2 in the presence of 1 mM GDP + 1 mM EDTA on FN-fibers deposited as intersections (continued to the next page).**

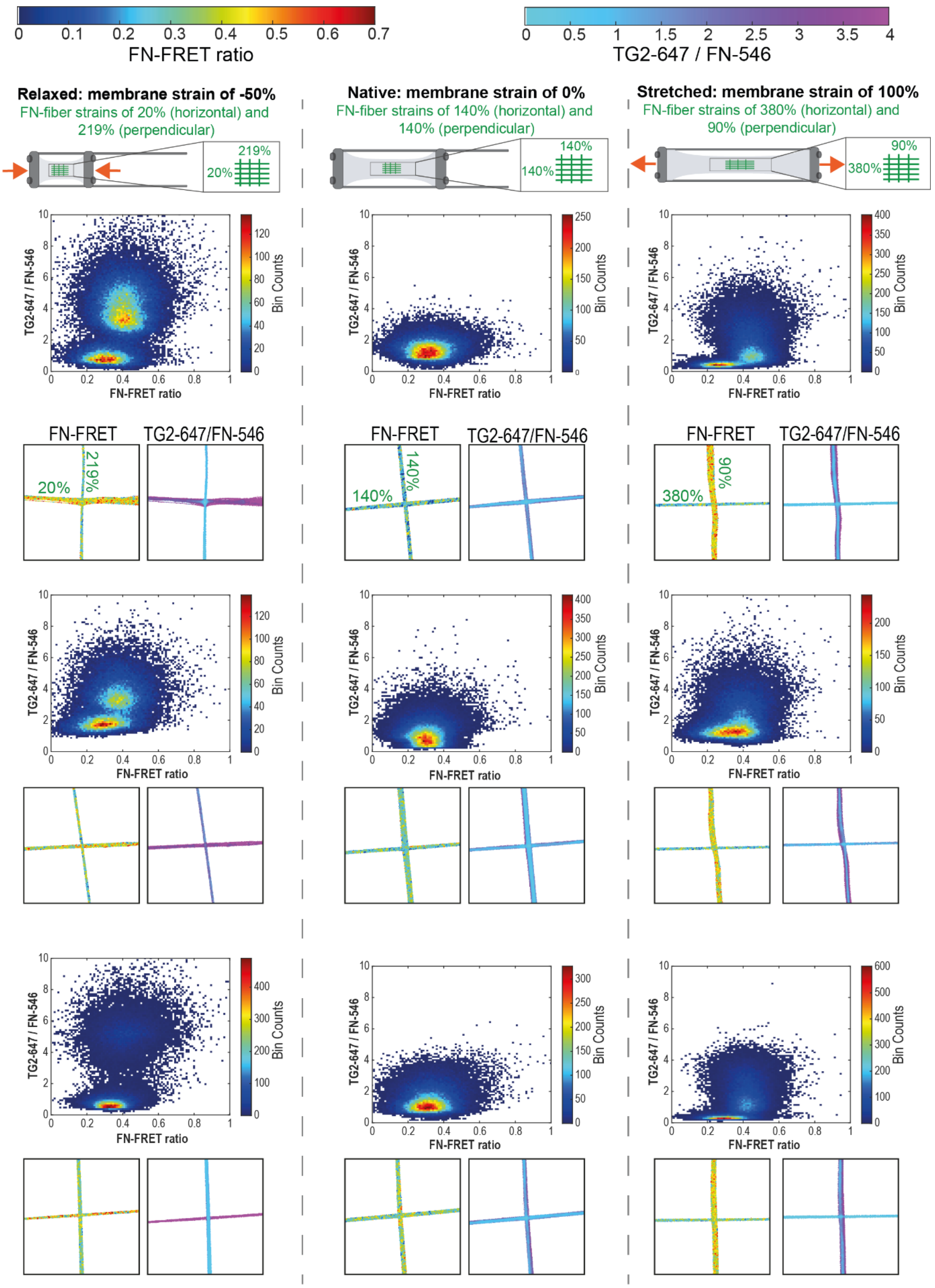
FN-FRET ratio
TG2-647 / FN-546
Relaxed: membrane strain of -50%
FN-fiber strains of 20% (horizontal) and 219% (perpendicular)
Native: membrane strain of 0%
FN-fiber strains of 140% (horizontal) and 140% (perpendicular)
Stretched: membrane strain of 100%
FN-fiber strains of 380% (horizontal) and 90% (perpendicular)
TG2-647 / FN-546
FN-FRET ratio
Bin Counts
FN-FRET
TG2-647/FN-546
20%
219%
140%
380%
90%

**Supplementary Figure 4: FN-fiber stretch assay performed with WT gpTG2 in the presence of 1 mM GDP+ 1 mM EDTA on FN-fibers deposited as intersections.** In the presence of 1 mM GDP and 1 mM EDTA, gpTG2 preferentially binds to FN fibers when they are under low tensional state (higher FN-FRET ratio). FRET-labelled FN fibers were deposited as intersections (horizontally and perpendicularly to the membrane stretch axis) on the silicone membrane mounted on the custom-made stretch device, which was mechanically stretched, relaxed, or left unchanged. The strains to which FN-fibers were subjected due to the silicone membrane stretching or relaxation are indicated, as was reported in a previous study[6]. On the native membrane, FN-fibers were pre-strained to 140% due to the forces required to pull fibers out of the droplet. FN-FRET and TG2-647/FN-546 pixels from each shown intersection were plotted as binned scatterplots. When the strain along the horizontal and perpendicular axes was different (stretched and relaxed membranes), FRET vs TG2-647/FN-546 pixels segregated into two separate groups. When the strain along the horizontal and perpendicular axes was the same (native membrane), pixels remain as a single population. This shows that TG2 binding to FN is regulated by FN tensional state with higher binding affinity to FN under low strain. 6 intersections per each membrane strain are shown.

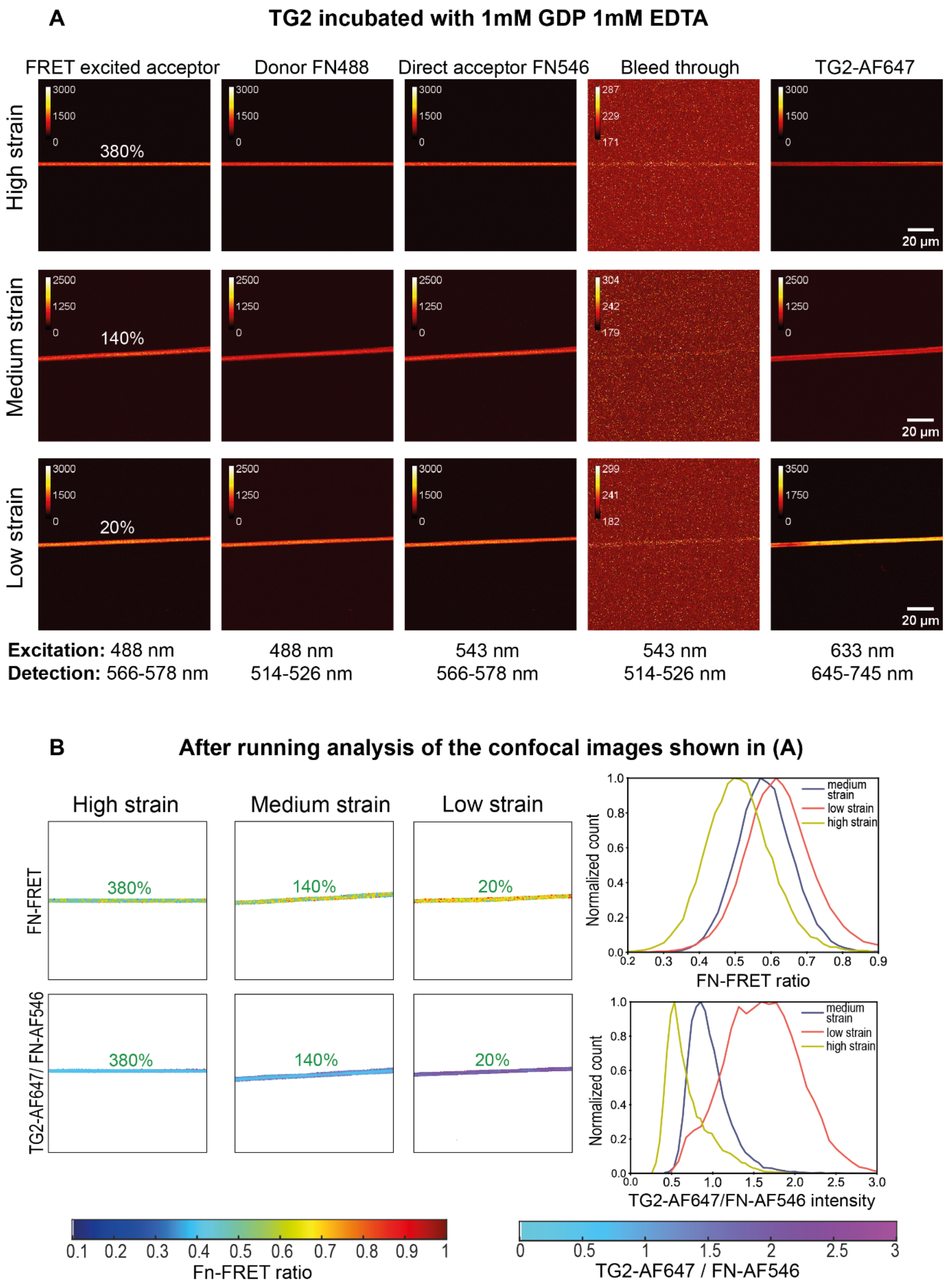

A
TG2 incubated with 1mM GDP 1mM EDTA
FRET excited acceptor
Donor FN488
Direct acceptor FN546
Bleed through
TG2-AF647
High strain
Medium strain
Low strain
380%
140%
20%
20 µm
Excitation: 488 nm
Detection: 566-578 nm
488 nm
514-526 nm
543 nm
566-578 nm
543 nm
514-526 nm
633 nm
645-745 nm
B
After running analysis of the confocal images shown in (A)
High strain
Medium strain
Low strain
FN-FRET
TG2-AF647/ FN-AF546
Normalized count
FN-FRET ratio
medium strain
low strain
high strain
TG2-AF647/FN-AF546 intensity
Fn-FRET ratio
TG2-AF647 / FN-AF546

**Supplementary Figure 5: TG2-647 binding to FN-FRET fibers horizontally deposited in the presence of 1 mM GDP and 1 mM EDTA, imaging, and image analysis illustration.** (A) Confocal images of a representative horizontally deposited FN-fiber for each membrane strain (low strain (relaxed membrane), medium strain (native membrane), high strain (stretched membrane)) are shown. Images were acquired as described in Supplementary figure 1. FN-fiber strains induced by the silicone membrane strain are indicated, as was previously reported[6]. (B) Confocal images shown in (A) were analyzed with the custom written Matlab script, which allows pixel-by-pixel correlation of normalized TG2 binding to the FN tensional state expressed by FN-FRET ratio[1]. Lower FN-FRET ratio corresponds to the higher FN tensional state, and higher FN-FRET ratio corresponds to the lower FN-FRET tensional state. Pixels of FN-intersections were color coded based of their corresponding FN-FRET ratio values. As can be seen, FN-fibers under each tensional strain, display a distinct range of heterogeneous FN-conformations as was previously observed in the ECM fibers assembled by fibroblasts[3,7]. FN-fibers which were subjected to low membrane strain are characterized by a range of pixels corresponding to the higher FN-FRET ratios, as expected. TG2 fluorescence intensity was normalized pixel-by-pixel to the directly excited FN-546 (TG2-647/FN-546), and these pixels were color coded based on the TG2-647/FN-546 value. Distributions of all FN-FRET and TG2-647/FN-546 pixels of a representative fiber were plotted as normalized histograms. TG2 preferentially binds FN under low strain. To increase the statistical significance, typically 15 horizontal fibers were analyzed in this manner per membrane strain in all subsequent experiments.

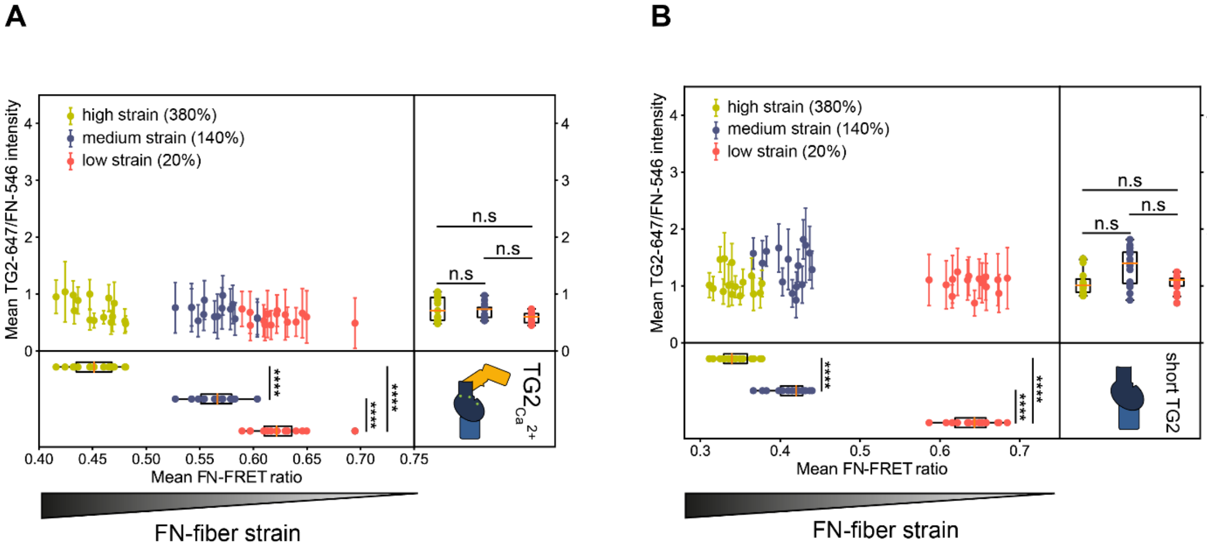


**Supplementary Figure 6**: **FN-fiber stretch assay with WT hrTG2 and short hrTG2(1-465aa).** Mean of the distribution of FN-FRET ratio in one fiber was plotted against the mean of the distribution of the pixel-by-pixel normalized TG2-647/FN-546 intensity. 15 fibers were analysed per each membrane strain. (A) WT hrTG2 incubated on FN-fibers under low, high, and medium strain in the presence of 10 mM $Ca^{2+}$ bound FN equally regardless of the FN-fiber strain. Higher FN-FRET ratio is indicative of low FN-fiber strain and vice versa. (B) short hrTG2 (β-barrels 1 and 2 deleted, 1-465aa) incubated on FN-fibers under low, high, and medium strain bound FN equally regardless of the FN-fiber strain. Higher FN-FRET ratio is indicative of low FN-strain and vice versa. Statistical significance was computed with Wilcoxon rank-sum statistic for two samples. P-values: (*$0.01 \leq p < 0.05$; **$0.001 \leq p < 0.01$; ***$10^{-5} \leq p < 0.001$; ****$10^{-6} \leq p < 10^{-5}$; ***** $p < 10^{-6}$)

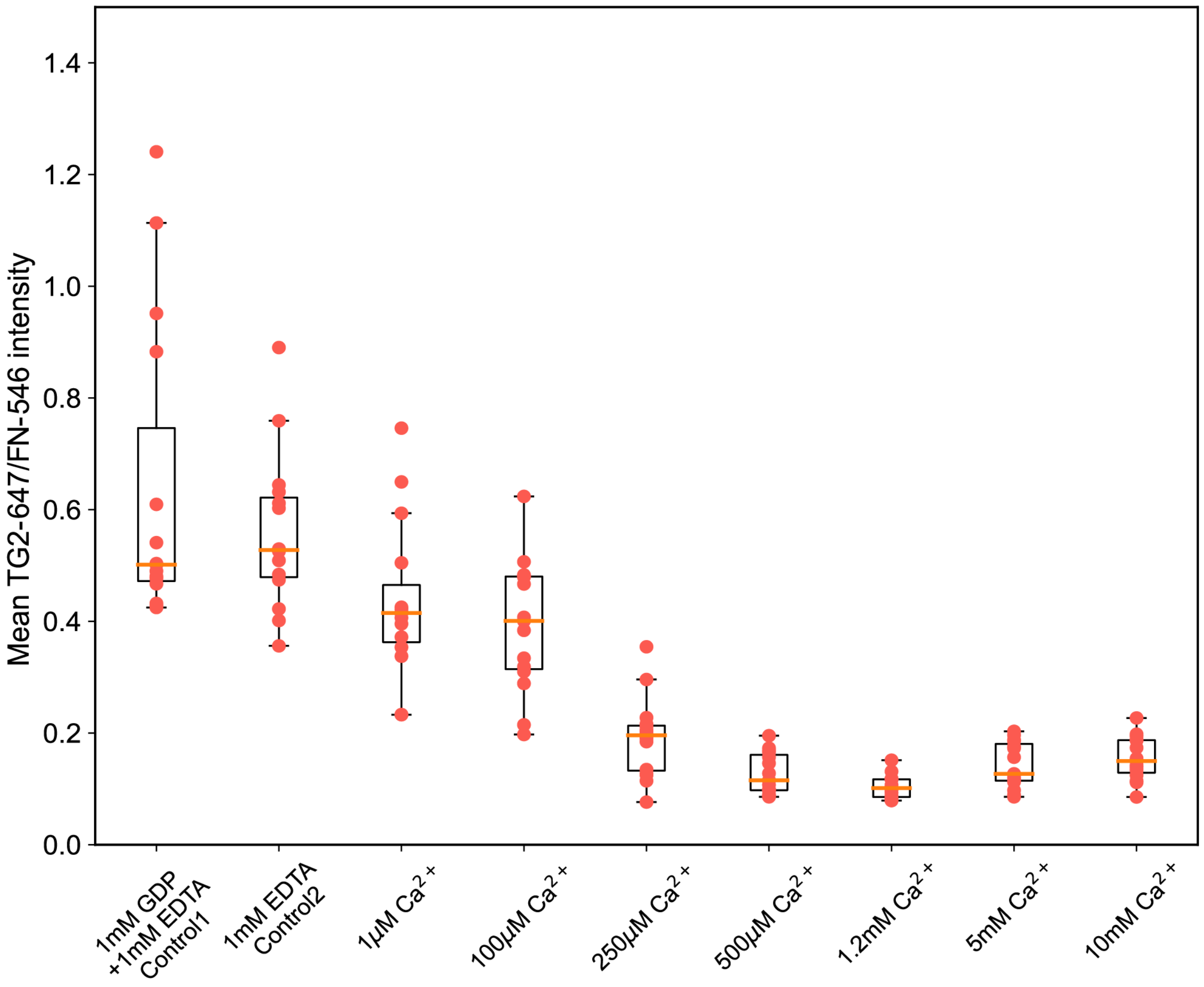


**Supplementary Figure 7: FN-fiber stretch assay performed with WT gpTG2 in the presence of increasing $Ca^{2+}$ concentrations on horizontally deposited FN-fibers under low strain (20%).** The means of pixel-by-pixel normalized TG2-647/FN-488 are plotted as boxplots. WT gpTG2 binding affinity for fibrillar FN under low strain (20%) decreases in a calcium dose-dependent manner. Progressively increasing calcium concentrations reduce gpTG2 binding affinity until it reaches the baseline affinity at 500µM calcium and does not decrease anymore. TG2 binding affinity to FN is the highest in the presence of 1mM GDP (control 1) or GDP-free EDTA (control 2). 15 individual fibers were analyzed per each condition.

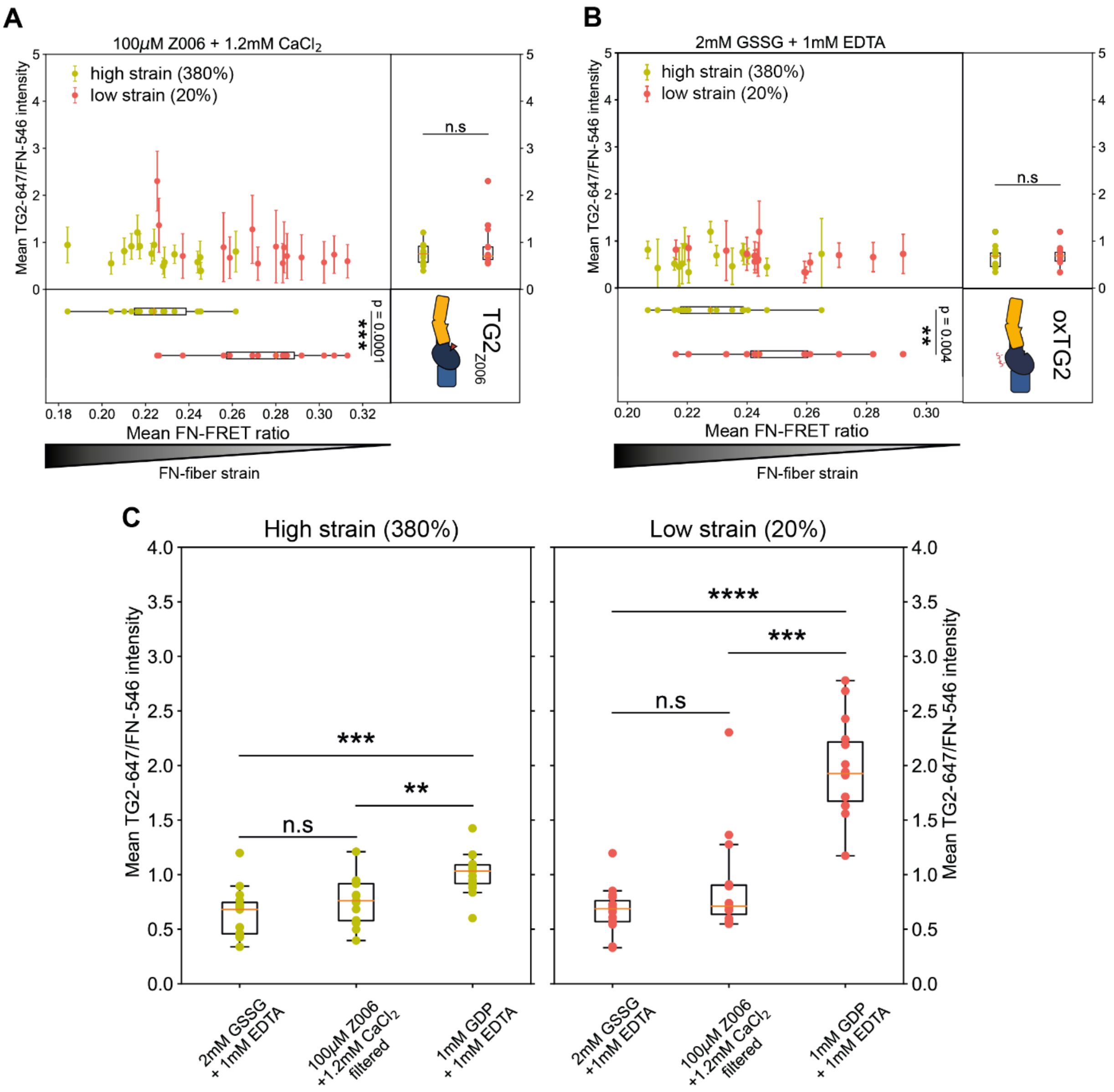


**Supplementary Figure 8: FN-fiber stretch assay with WT gpTG2.** Full data sets of the results shown in Fig 3D-3F. Mean of the distribution of FN-FRET ratio in one fiber was plotted against the mean of the distribution of the pixel-by-pixel normalized TG2-647/FN-546 intensity. 15 fibers were analysed per each membrane strain. When WT gpTG2 is trapped in the extended open-state either with inhibitor Z006 **(A)** or through oxidation **(B)**, TG2 binds FN-fibers under high (~380%) and low (~20%) strain equally. **C:** Means of TG2-647/FN546 intensity shown in (A) and (B) were replotted as boxplots to make comparisons between oxTG2, $TG2_{Z006}$ and $TG2_{GDP}$ when FN-fibers experience the same strain. There was no significant difference in binding between oxTG2 and $TG2_{Z006}$ to FN-fibers under high (~380%) and low (~20%) strain, however $TG2_{GDP}$ binding affinity was significantly higher compared to oxTG2 and $TG2_{Z006}$ on FN-fibers under high and low strain. 15 individual fibers were analyzed per each condition. Statistical significance was computed with Wilcoxon rank-sum statistic for two samples. P-values: (*$0.01 \leq p < 0.05$; **$0.001 \leq p < 0.01$; ***$10^{-5} \leq p < 0.001$; ****$10^{-6} \leq p < 10^{-5}$; ***** $p < 10^{-6}$)

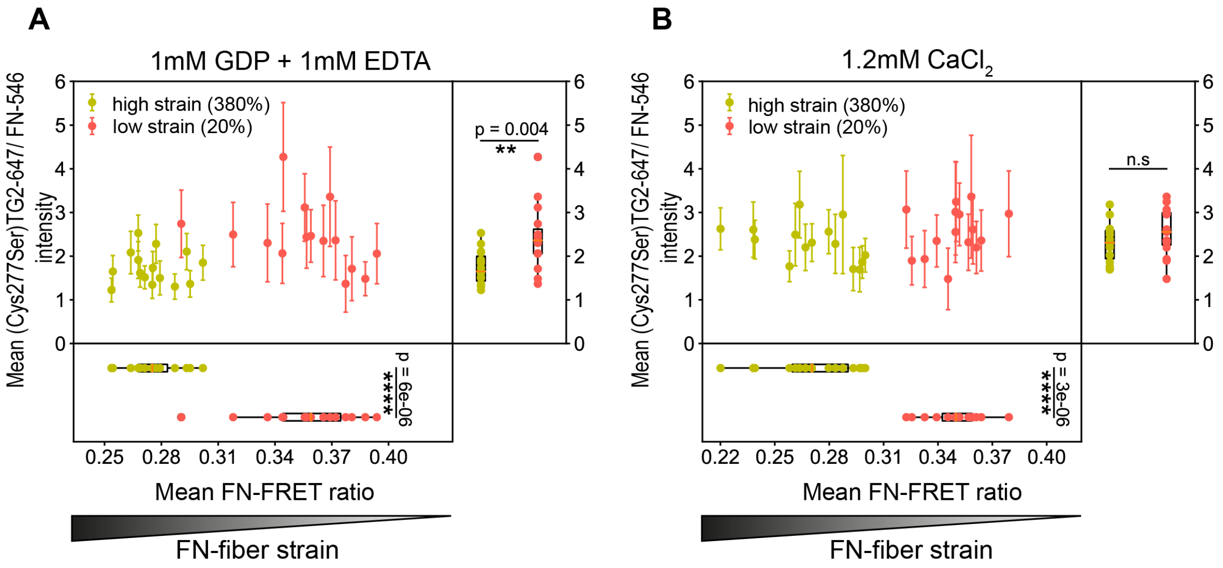


**Supplementary Figure 9: FN-fiber stretch assay performed with hrTG2's inactive Cys277Ser mutant**. **A:** In the presence of 1mM GDP, the Cys277Ser hrTG2 preferentially bound FN fibers under low strain, although the significance of this difference was less (p=0.004) compared to the significance of the difference in affinity we observed with the WT gpTG2 (p < 4e-06). This is consistent with the report that hrTG2's Cys277Ser mutant is unable to bind GTP/GDP efficiently[9]. **B:** In the presence of 1.2 mM $Ca^{2+}$, Cys277Ser hrTG2 bound FN fibers under high (380%) and low (20%) strain equally, similarly to the result with the WT TG2 and 1.2 mM $Ca^{2+}$. Since hrTG2's Cys277Ser mutant is prone to adopt an open conformation[9], this result confirms that it is the open conformational state, and not the crosslinking activity of the open-state of TG2, which plays the role in the loss of the strain-sensitive binding to FN. 15 fibers were analysed per each membrane strain. Statistical significance was computed with Wilcoxon rank-sum statistic for two samples. P-values: (*0.01≤ p <0.05; **0.001≤ p <0.01; ***$10^{-5}$≤ p <0.001; ****$10^{-6}$≤ p <$10^{-5}$; ***** p <$10^{-6}$)

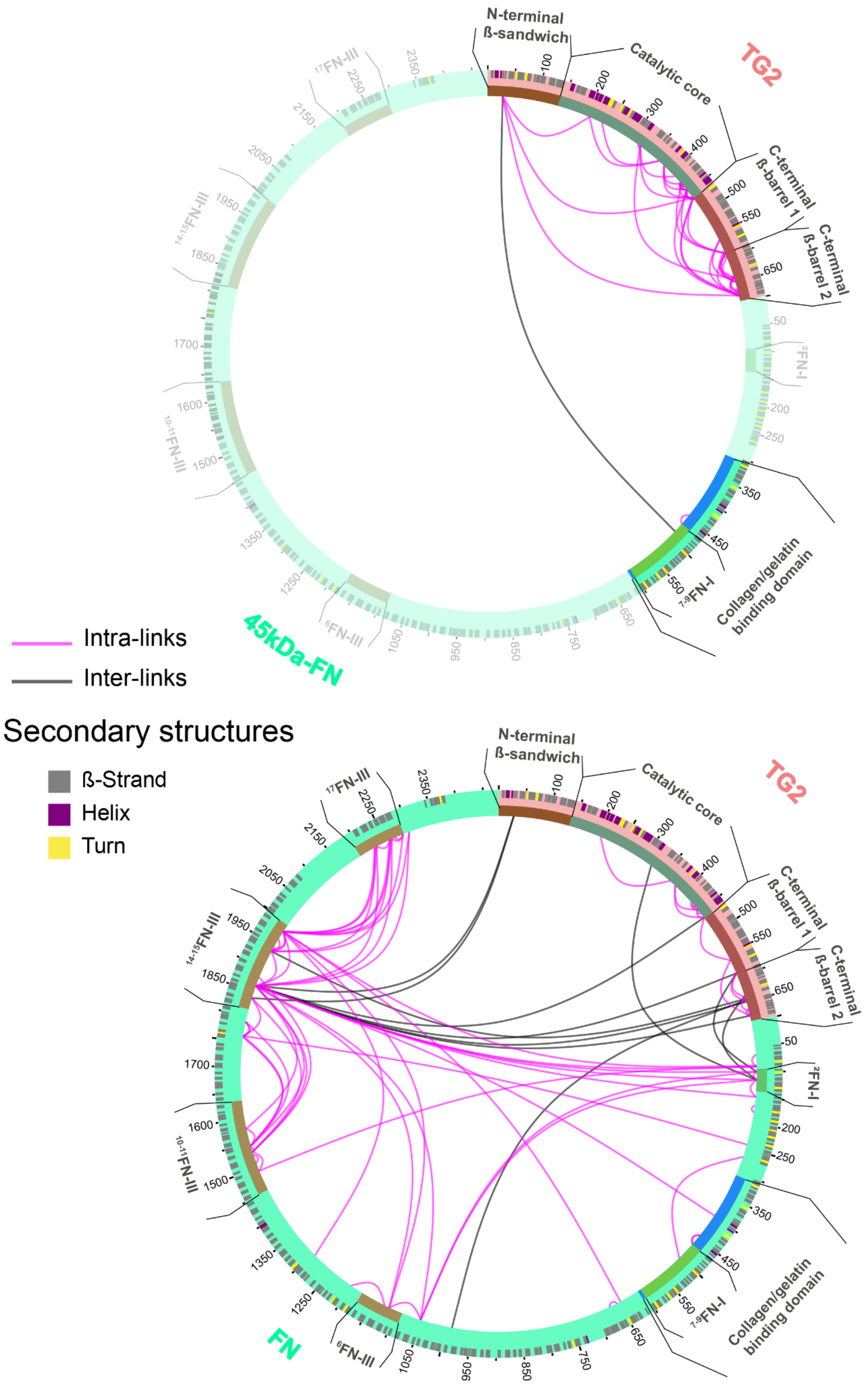
N-terminal ß-sandwich
Catalytic core
C-terminal ß-barrel 1
C-terminal ß-barrel 2
TG2
45kDa-FN
Collagen/gelatin binding domain
7-9FN-I
2FN-I
6FN-III
10-11FN-III
14-15FN-III
17FN-III
Intra-links
Inter-links
Secondary structures
ß-Strand
Helix
Turn
FN

**Supplementary Figure 10: XL-MS of TG2/45 kDa-FN complex and TG2/FN complex using trypsin as a protease.** Circular diagrams show crosslinks identified from TG2/45 kDa-FN complex (top) and from TG2/FN complex (bottom). Inter-protein crosslinks are shown in black and intra-protein crosslinks in magenta. **Top:** The GBD is shown in solid colors and the rest of the FN is dimmed. A single inter-protein crosslink could be identified connecting FN residue Lys486 ($FNI_7$, numbering for full-length FN) with TG2 residue Lys30 (N-terminal β-sandwich). **Bottom:** TG2 in complex with full-length FN, Lys30 (TG2) was crosslinked to residues Lys1837 and Lys1862 ($FNIII_{14}$). The C-terminal β-barrel 2 interacts with $FNIII_{14-15}$ (residues Lys1862 and Lys1936) and $FNI_2$ (residues Asp110 and Lys116). Simultaneously, $FNIII_{14-15}$ and $FNI_2$ regions are in close contact with each other, as shown by the magenta intra-protein crosslinks. Crosslinks are mapped using FN monomer. Visualization was done with xVis Webserver[10]**.**

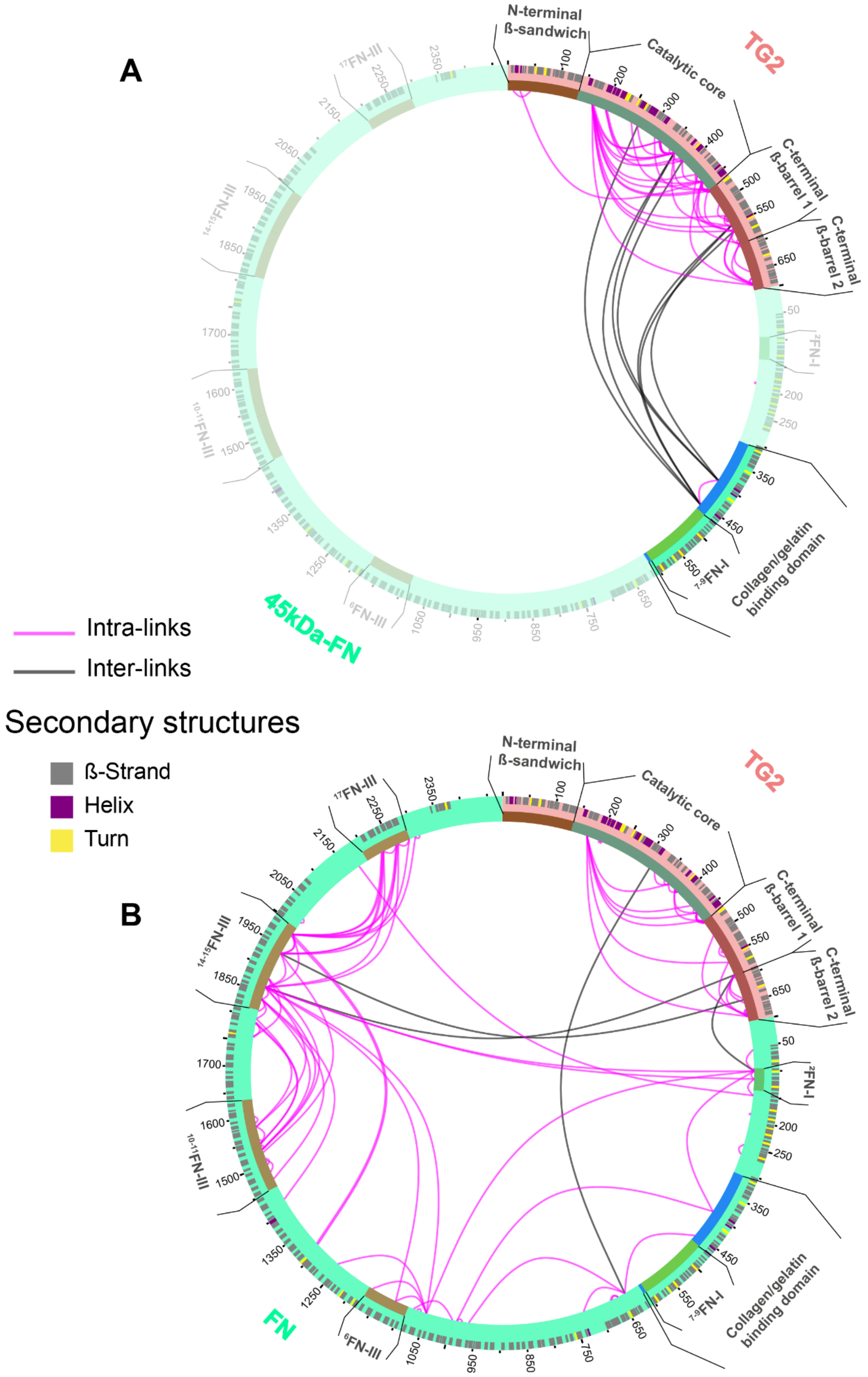


**Supplementary Figure 11: XL-MS of TG2-45kDa-FN complex and TG2-FN complex using chymotrypsin as a protease.** Continued to the next page.

C

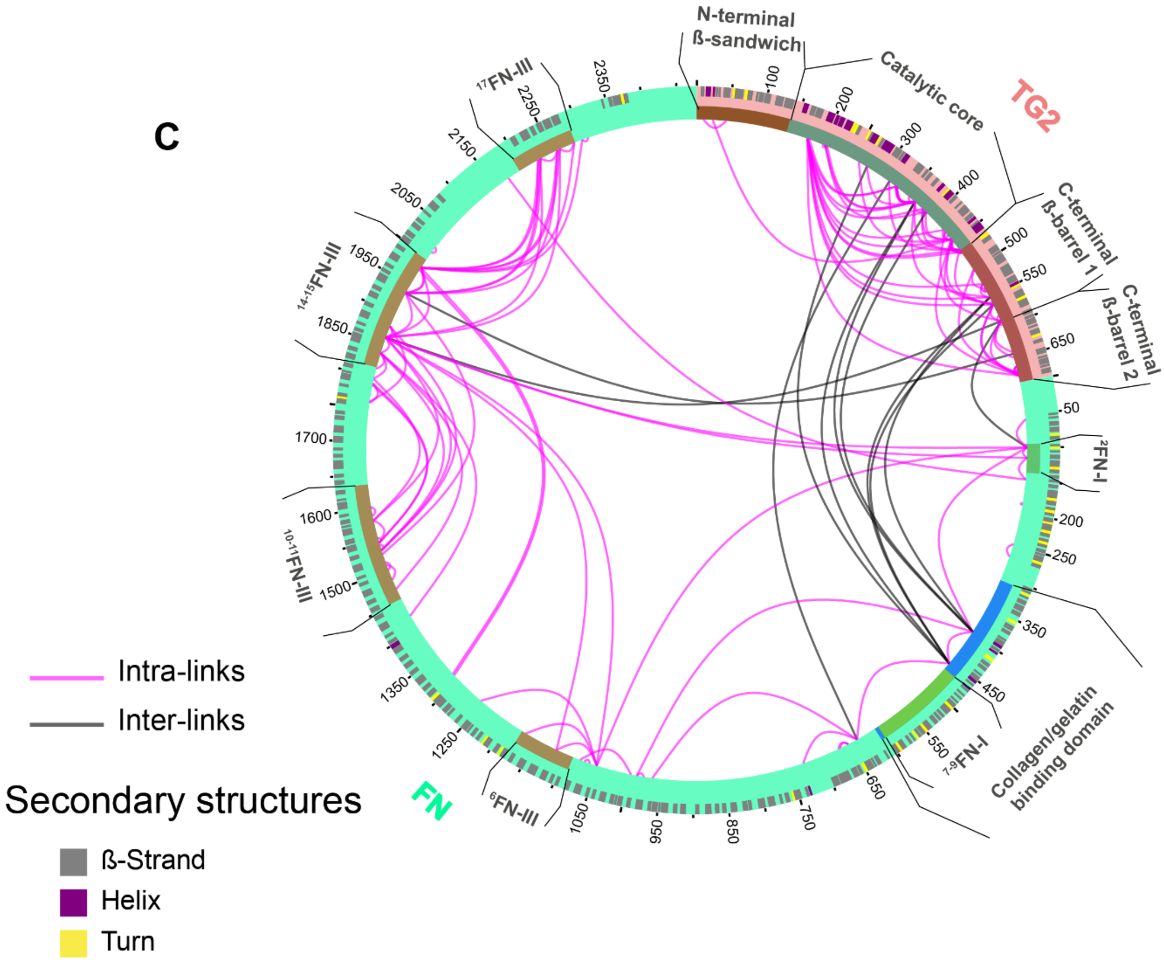


**Supplementary Figure 11: XL-MS of TG2-45kDa-FN complex and TG2-FN complex using chymotrypsin as a protease.** Circular diagrams show crosslinks identified from TG2/45 kDa-FN complex (top) and TG2/FN complex (bottom). Inter-protein crosslinks are shown in black and intra-protein crosslinks in magenta. **A:** The GBD is shown in bright colors and the rest of the FN is dimmed. Using chymotrypsin as a protease we were able to identify additionally crosslinked sites on Lys397 and Lys457 ($FNII_1$ and $FNII_2$ respectively, numbering for full-length FN). **B:** The C-terminal β-barrel 2 interacts with $FNIII_{14-15}$ (residues Lys1862 and Lys1936) and $FNI_2$ (residue Asp110). Simultaneously, $FNIII_{14-15}$ and $FNI_2$ regions are in close contact with each other, as shown by the magenta intra-links. **C:** Shows all crosslinks from A and B visualized together in one circular diagram. Crosslinks are mapped using FN monomer. Visualization was done with xVis Webserver[10].

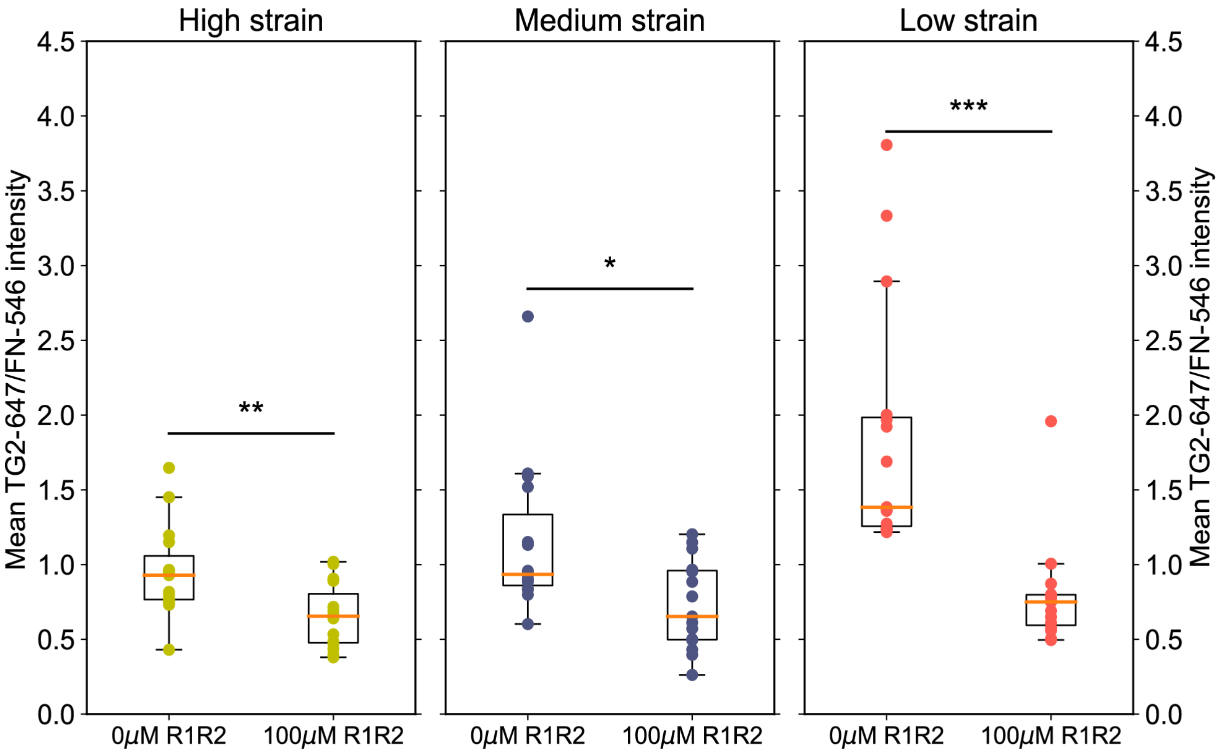


**Supplementary Figure 12: Competition FN-fiber stretch assay between the collagen-mimicking peptide R1R2 (Streptococcus equi) and WT gpTG2 in the presence of 1mM GDP and 1mM EDTA.** Means of TG2-647/FN-546 intensity shown in Fig.7C were replotted as boxplots to make comparisons between $TG2_{GDP}$ binding in the presence of 0μ MR1R2 and 100 μM R1R2 when FN-fibers experience the same strain. Mechanical stretching of FN-fibers reduces their affinity for $TG2_{GDP}$ due to the destruction of many TG2 binding sites on FN. Addition of 100 μM R1R2 further significantly reduces $TG2_{GDP}$ binding affinity to FN-fibers experiencing high (380%), medium (140%) and low (20%) strains. 15 fibers were analysed per each condition. Statistical significance was computed with Wilcoxon rank-sum statistic for two samples. P-values: (*$0.01 \leq p < 0.05$; **$0.001 \leq p < 0.01$; ***$10^{-5} \leq p < 0.001$; ****$10^{-6} \leq p < 10^{-5}$; ***** $p < 10^{-6}$)

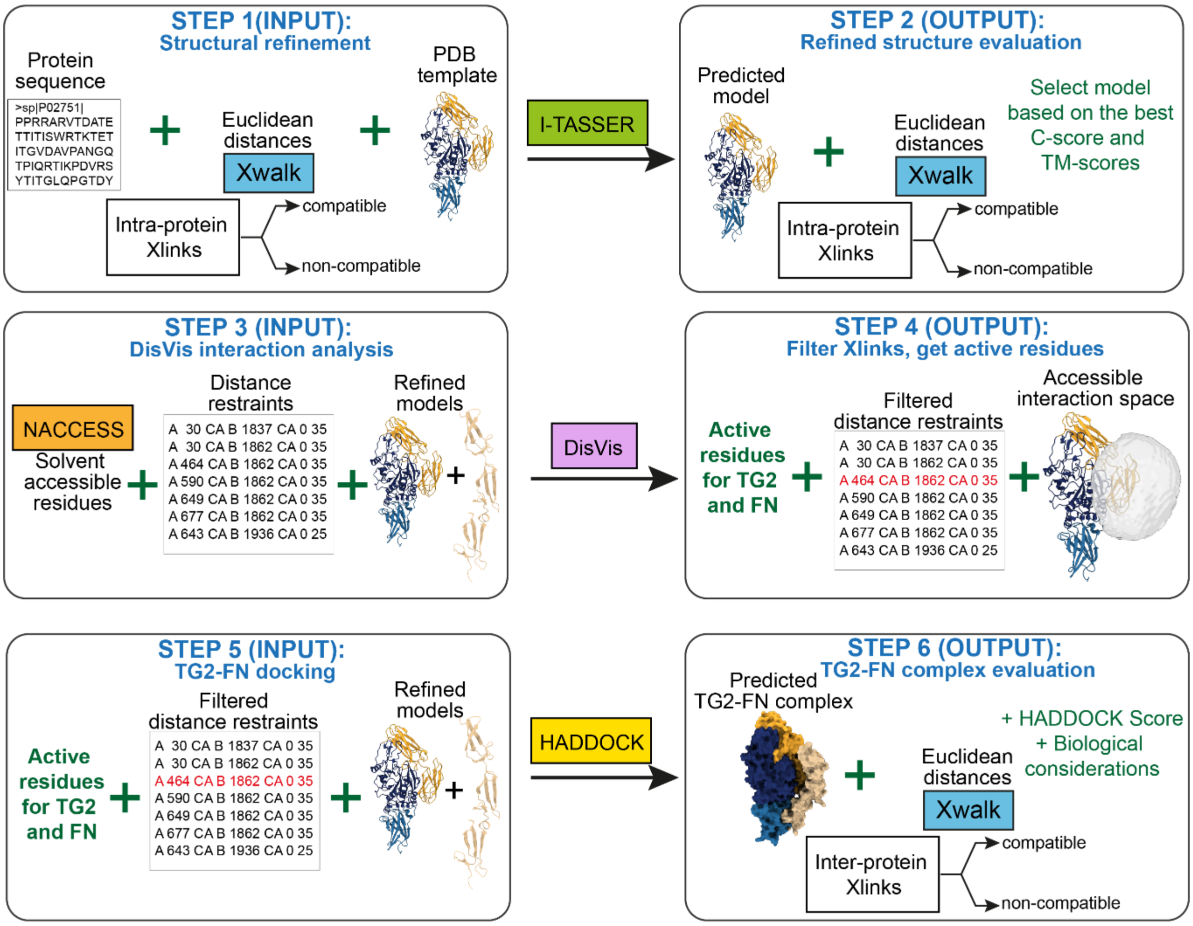


**Supplementary Figure 13: Schematic overview of the integrative structural modelling workflow of TG2 complexed to FN c, which is presented in detail in Supplementary Note 1-3. (STEP 1-STEP2) Structure prediction and structure refinement:** Structure prediction and structural refinement of available PDB templates was performed with I-TASSER[11]. At this step, PDB templates, corresponding amino acid sequences and experimental intra-protein crosslinks were submitted to I-TASSER. (STEP 1). Classification of crosslinks into "compatible" and "non-compatible" was performed with Xwalk[12] by calculation of Euclidean distances between the crosslinked residues. The best mode was selected for the next modelling steps based on the confidence score (C-score) and TM-score (STEP 2). **(STEP 3- STEP4) Analysis of accessible interaction space:** To predict active residues at the binding interfaces and to evaluate the compatibility of inter-protein crosslinks, DisVis[13] was employed. Solvent accessibility of residues on refined models obtained from STEP 2 was calculated with NACCESS[14]. Only residues with a relative solvent accessibility of at least 40% for either the main chain or the side chai were selected. Refined models, solvent accessible residues, and inter-protein crosslinks in form of distance restraints were submitted to DisVis for analysis of accessible interaction space (STEP 3). Incompatible inter-protein crosslinks, if any, were excluded from further analysis; active residues at the predicted binding interface were obtained for the protein-protein docking step. **(STEP 5- STEP6) Protein-protein docking:** Active residues, filtered inter-protein crosslinks in form of distance restraints and refined models were submitted to HADDOCK[15] for docking. The best models of predicted protein complexes were selected based on the HADDOCK score and compatibility of experimental crosslinks with resulting models.

**Supplementary Note 1: Structure prediction and structure refinement.**

**Structural modelling and refinement of 3D crystal structure templates of individual fibronectin fragments and TG2 using intra-protein crosslinks.**

To ensure accuracy in structural modelling of FN fragments, we exclusively used the UniProt P02751-1 FINC_HUMAN (isoform 1) sequence. This decision was based on the results of our XL-MS experiments, which unequivocally identified the presence of FN isoform 1. Additionally, our modelling approach considered the detected crosslinks between TG2 and FN, as well as the knowledge of FN domains in UniProt and the availability of PDB template structures. As a result, we selected the following FN fragments for modelling:

| FN fragment/domains | FN aa sequence in UniProt P02751-1 | PDB structure available? | Number of detected inter-protein crosslinks with TG2 |
|---|---|---|---|
| $FNI_{2-3}$ | 94-183 | PDB: 2cg6; 2cg7; 2rkz; 3cal; 3zrz; 4pz5 | 3 |
| $FNI_{6}$-$FNII_{1}$-$FNII_{2}$-$FNI_{7-9}$(GBD) | 306-604 | Partially. PDB:3mql; PDB:3ejh | 8 |
| $FNI_{7-9}$ | 467-604 | Partially. PDB:3ejh, 3gxe | 1 |
| $FNIII_{14-15}$ | 1816-1995 | PDB:1fnh; 3r8q | 7 |

To model and refine protein structures we followed the previously published protocols[16,17]. Briefly, we selected I-TASSER[11,18] server (iterative threading assembly refinement) for structural prediction and refinement of available templates, because this server allows to integrate experimental distance restraints from XL-MS into structure prediction and structure refinement. Intra-protein crosslinks were classified as compatible or non-compatible by measuring the Euclidean distance (ED) between the β-carbons of crosslinked residues using Xwalk[12].The filtering of non-compatible crosslinks was based on specific Euclidean distance cutoff, above which the crosslink was defined as non-compatible[16,19]: ED for DSS < 35 Å, ED for DMTMM < 25 Å, ED for PDH < 35Å. If there were both compatible and non-compatible crosslinks, we attempted structural refinements of available PDB templates using all, only compatible or only non-compatible crosslinks, to calculate, if possible, a structure of an alternative conformation to which non-compatible crosslinks could belong. To assess the quality of predicted structures, experimental crosslinks were mapped, and CB-CB distances were measured again. We then selected the best model for further modelling steps.

**$FNI_{2\text{-}3}$ structure refinement and evaluation**

Submitted amino acid sequence (UniProt P02751-1 94-183aa):

```
EETCFDKYTGNTYRVGDTYERPKDSMIWDCTCIGAGRGRISCTIANRCHEGGQSYKIGDTWRRPHETG
GYMLECVCLGNGKGEWTCKPIA
```

Several templates of the $FNI_{2\text{-}3}$ region were available in the PDB. We selected PDB:2cg7 as a template because of its high resolution. We detected in total four unique crosslinks within $FNI_{2\text{-}3}$, all of which were compatible with the PDB:2cg7 template. We submitted all these crosslinks, along with the PDB:2cg7 template to I-TASSER for further structural refinement. For the two DMTMM crosslinks, we set the distance restraint at 25 Å, while for the other two DSS crosslinks, we set a distance restraint at 35 Å. The best model was selected based on the confidence score (C-score=0.74) and estimated TM-score=0.81+/-0.09, which indicated high degree of structural similarity and correct topology, as evidenced by TM-score>0.5. Importantly, this model satisfied all of the detected crosslinks.

**Crosslinked residues within $FNI_{2\text{-}3}$:**

| AbsPos1 | AbsPos2 | Link | Crosslinker | CB-CB Euclidean distances (Å) Xwalk |
|---|---|---|---|---|
| 166 | 180 | GLU-LYS | DMTMM | 6.3 |
| 116 | 149 | LYS-LYS | DSS | 16.4 |
| 100 | 116 | LYS-LYS | DSS | 10.4 |
| 95 | 100 | GLU-LYS | DMTMM | 16.6 |

*Crosslinks within FNI$_{2-3}$ shown as Euclidean distances mapped onto the predicted model (compatible crosslinks - cyan):*

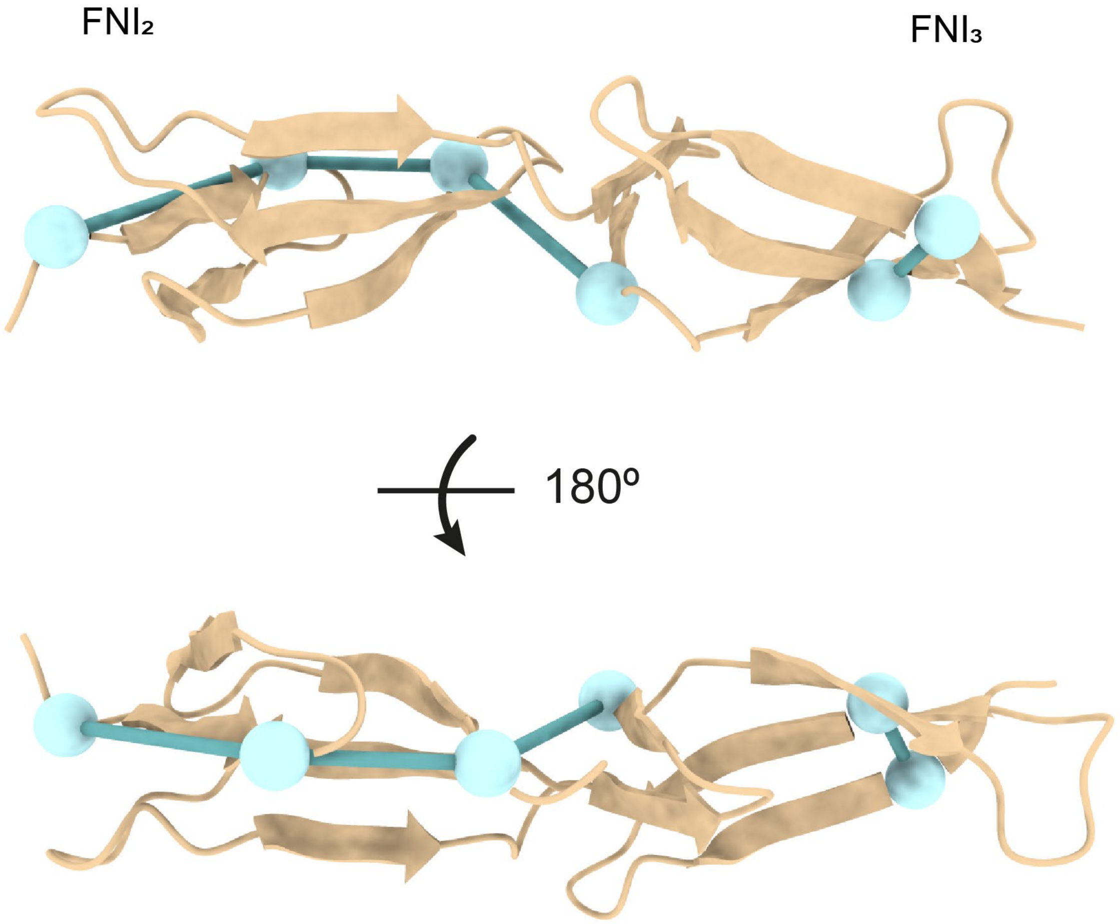

**GBD ($FNI_6$-$FNII_1$-$FNII_2$-$FNI_{7-9}$) structure modelling and evaluation**

Submitted amino acid sequence (UniProt P02751-1 306-604aa):

```
GHCVTDSGVVYSVGMQWLKTQGNKQMLCTCLGNGVSCQETAVTQTYGGNSNGEPCVLPFTYNGRTFYS
CTTEGRQDGHLWCSTTSNYEQDQKYSFCTDHTVLVQTRGGNSNGALCHFPFLYNNHNYTDCTSEGRRD
NMKWCGTTQNYDADQKFGFCPMAAHEEICTTNEGVMYRIGDQWDKQHDMGHMMRCTCVGNGRGEWTCI
AYSQLRDQCIVDDITYNVNDTFHKRHEEGHMLNCTCFGQGRGRWKCDPVDQCQDSETGTFYQIGDSWE
KYVHGVRYQCYCYGRGIGEWHCQPLQT
```

Currently, two PDB crystal structures (PDB:3mql and PDB:3ejh) cover regions 306-513aa ($FNI_6FNII_{1-2}FNI_7$) and 516-606aa ($FNI_{8-9}$), respectively, providing coverage for the entire GBD, except for the short 513-516aa linker that connects FNI7 to FNI8. As a result, the orientation of FNI6FNII1-2FNI7 and FNI8-9 regions relative to one another is not known. Initially, we attempted to model the entire GBD using I-TASSER to understand the orientation of FNI6FNII1-2FNI7 and FNI8-9 regions relative to one another, but the results were not satisfactory. Therefore, we utilized ROBETTA[20] and Alphafold to model the GBD. To assess the quality of the predicted structures, we utilized QMEAN[21] tool, which compares the degree of "nativeness" of the predicted model with experimental PDB structures. The predicted models from ROBETTA were more than one standard deviation away (QMEAN=1.86 and 1<|Z-score|<2), whereas Alphafold predicted structure demonstrated better scores (QMEAN =1.44 and |Z-score|<1). To further evaluate the quality of structures, we measured CB-CB distances with Xwalk on the predicted models and compared them to the distances between the same residues on the experimental crystal structure PDB:3mql. Interestingly, the Alphafold predicted structure had a shorter distance between Lys457-Lys397 (35.2 Å) compared to the distance between same residues on the crystal structure (36.5 Å), indicating that the predicted model better satisfied the crosslink. However, since this crosslink is located in a flexible region, structural rearrangements can be expected. Based on the above findings, the Alphafold predicted model was selected for further modelling steps.

**Crosslinked residues within GBD ($FNI_6$-$FNII_1$-$FNII_2$-$FNI_{7-9}$):**

| AbsPos1 | AbsPos2 | Link | Crosslinker | CB-CB Euclidean distances (Å) Xwalk |
|---|---|---|---|---|
| 467 | 444 | GLU-LYS | DMTMM | 10.3 |
| 457 | 397 | LYS-LYS | DSS | 35.2 |
| 468 | 444 | GLU-LYS | DMTMM | 15.2 |

*Crosslinks within GBD shown as Euclidean distances mapped onto the predicted model (compatible crosslinks - cyan):*

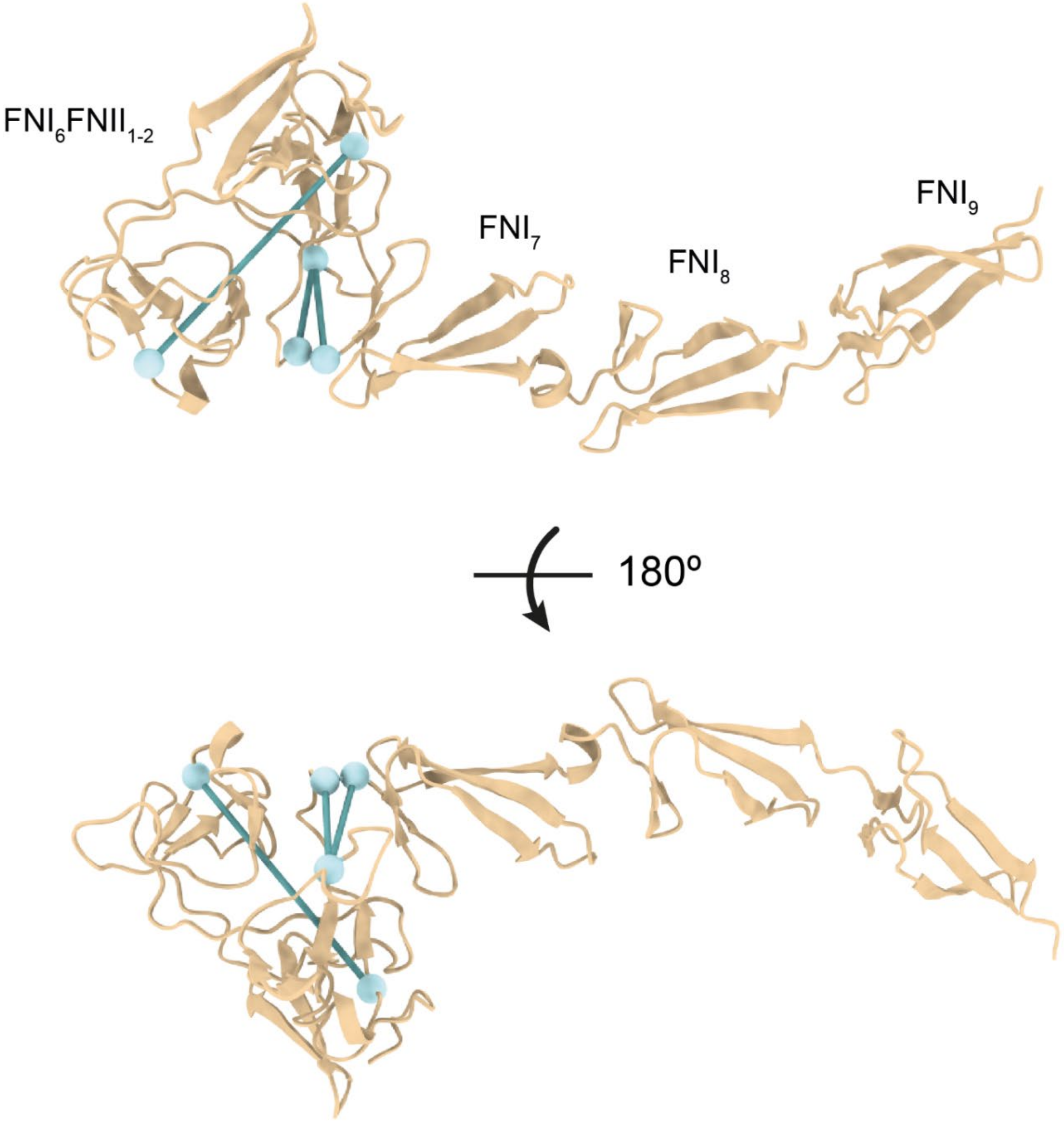

**$FNI_{7\text{-}9}$ structure modelling and evaluation**

Submitted amino acid sequence (UniProt: P02751-1 467-604aa):

EEICTTNEGVMYRIGDQWDKQHDMGHMMRCTCVGNGRGEWTCIAYSQLRDQCIVDDITYNVNDTFHKR
HEEGHMLNCTCFGQGRGRWKCDPVDQCQDSETGTFYQIGDSWEKYVHGVRYQCYCYGRGIGEWHCQPL
QT

No crosslinks could be detected within the $FNI_{7\text{-}9}$ region, which can be explained by two factors. Firstly, this region has low number of lysine residues, which are necessary for DSS reagents to crosslink on primary amines. As a result, generating DSS crosslinked peptides becomes more challenging in this case. Secondly, protein interaction interfaces can often be protected from crosslinking due to protein-protein contacts. Thus, crosslinks usually form around the contact sites rather than directly at the contact site itself. It is well-known that $FNI_{7\text{-}9}$ is the primary high-affinity interaction site for TG2. Consistent with this, only one crosslinked residue was found on the N-terminal domain of TG2, where the main interaction site for FN is located. To model the $FNI_{7\text{-}9}$ region, it was necessary to establish the orientation of $FNI_7$ and of $FNI_{8\text{-}9}$ regions with respect to each other. Our initial attempts to model the entire $FNI_{7\text{-}9}$ region with I-TASSER were not successful, so we utilized ROBETTA and Alphafold for modelling of this region. Since no crosslinks were available to assess the quality of the predicted model, we utilized QMEAN to select the best model. The highest scoring model predicted by Alphafold had a score that compared well with experimental PDB structures within one standard deviation (QMEAN=0.64 and |Z-score|<1). In contrast, ROBETTA predicted models had a worse score (QMEAN=2.02 and 1<|Z-score|<2), placing the predicted model more than one standard deviation away from experimental PDB structures of similar qualities. Therefore, Alphafold predicted model was selected for further modelling steps.

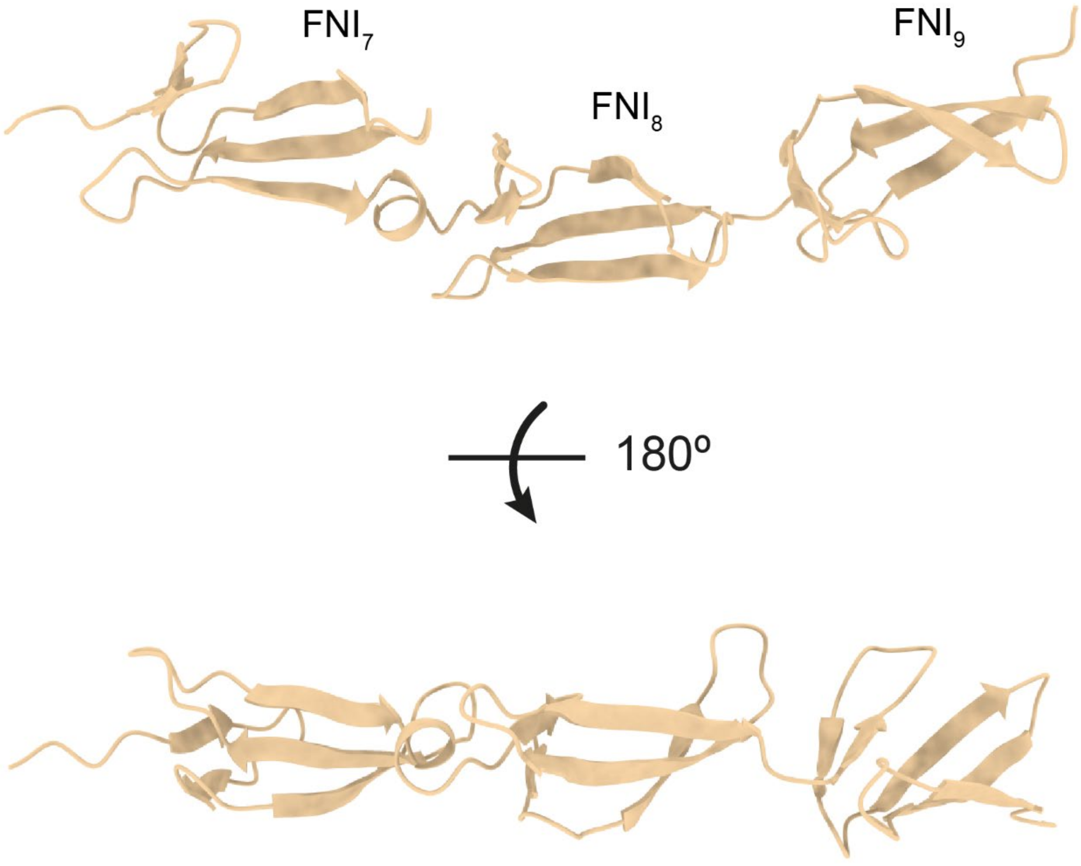

**FNIII$_{14-15}$ structure refinement and evaluation**

Submitted amino acid sequence (UniProt: P02751-1 1816-1995aa):

```
PPRRARVTDATETTITISWRTKTETITGFQVDAVPANGQTPIQRTIKPDVRSYTITGLQPGTDYKIYL
YTLNDNARSSPVVIDASTAIDAPSNLRFLATTPNSLLVSWQPPRARITGYIIKYEKPGSPPREVVPRP
RPGVTEATITGLEPGTEYTIYVIALKNNQKSEPLIGRKKTDELP
```

We selected PDB:1fnh to use as a suitable template for modelling of FNIII$_{14-15}$ region because it covered the entire region we aimed to model. We were able to identify 11 crosslinks within FNIII$_{14-15}$ region, of which only one (Lys1936-Lys1862) did not comply with the selected distance cutoff. on the PDB:1fnh template. We performed structural refinement of the PDB:1fnh template using I-TASSER in three steps: using all crosslinks (compatible and non-compatible), only compatible crosslinks, or only one non-compatible crosslink. We set the distance restraint at 35 Å for the 7 DSS crosslinks and 25 Å for the 4 DMTMM crosslinks. Recalculation of structures did not produce any distinct models with alternative conformation. The models refined using all crosslinks or only compatible crosslinks had identical C-scores and TM-scores (C-score=0.80; TM-score=0.82+/-0.08). The model derived by structural refinement of the template using one non-compatible crosslink had a slightly lower C-score=0.79. However, all structures satisfied 10/11 crosslinks, indicating the similarity of the protein in solution to the model. One non-compatible crosslink connects residues one both the FNIII$_{14}$ and FNIII$_{15}$ domains, which are connected by a flexible linker and can move with respect to one another. This suggests that in an ensemble of conformations, some of the molecules were present in a more compact conformation. Therefore, we selected model refined with all crosslinks.

**Crosslinked residues within FNIII$_{14-15}$:**

| AbsPos1 | AbsPos2 | Link | Crosslinker | CB-CB Euclidean distances (Å) Xwalk |
|---|---|---|---|---|
| 1981 | 1983 | LYS-GLU | DMTMM | 7.8 |
| 1946 | 1939 | GLU-LYS | DMTMM | 8.7 |
| 1964 | 1939 | GLU-LYS | DMTMM | 7.5 |
| 1981 | 1862 | LYS-LYS | DSS | 34.8 |
| 1936 | 1981 | LYS-LYS | DSS | 11.0 |
| 1981 | 1939 | LYS-LYS | DSS | 22.2 |
| 1936 | 1862 | LYS-LYS | DSS | 43.7 |
| 1936 | 1939 | LYS-LYS | DSS | 11.9 |
| 1936 | 1983 | LYS-GLU | DMTMM | 13.1 |
| 1862 | 1880 | LYS-LYS | DSS | 23.0 |
| 1862 | 1837 | LYS-LYS | DSS | 13.2 |

*Crosslinks within* FNIII$_{14\text{-}15}$ *shown as Euclidean distances mapped onto the predicted model (compatible crosslinks – cyan; non-compatible crosslinks - magenta):*

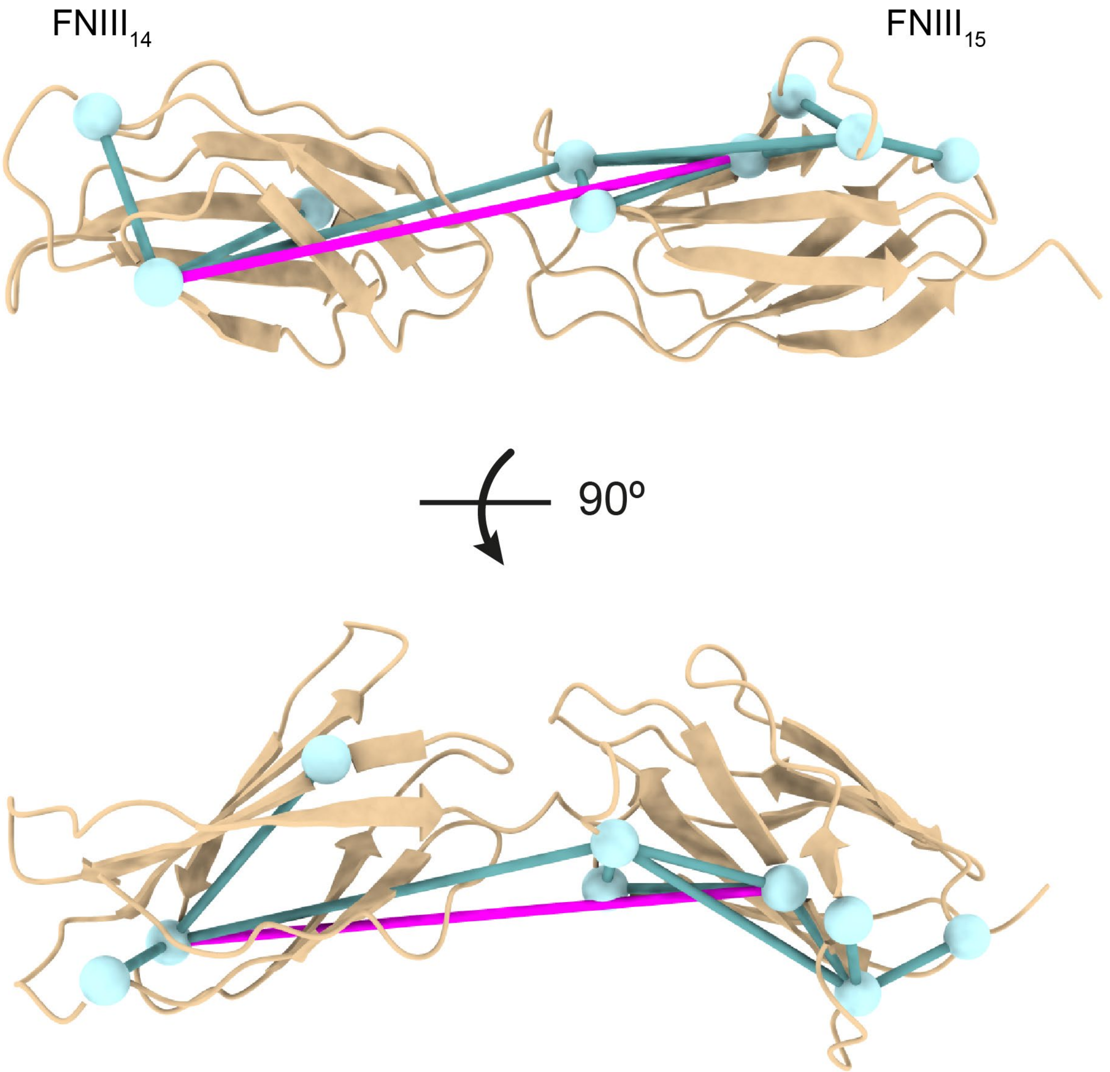

**TG2 structure refinement and evaluation**

Submitted amino acid sequence (UniProt P21980):

```
MAEELVLERCDLELETNGRDHHTADLCREKLVVRRGQPFWLTLHFEGRNYEASVDSLTFS
VVTGPAPSQEAGTKARFPLRDAVEEGDWTATVVDQQDCTLSLQLTTPANAPIGLYRLSLE
ASTGYQGSSFVLGHFILLFNAWCPADAVYLDSEEERQEYVLTQQGFIYQGSAKFIKNIPW
NFGQFEDGILDICLILLDVNPKFLKNAGRDCSRRSSPVYVGRVVSGMVNCNDDQGVLLGR
WDNNYGDGVSPMSWIGSVDILRRWKNHGCQRVKYGQCWVFAAVACTVLRCLGIPTRVVTN
YNSAHDQNSNLLIEYFRNEFGEIQGDKSEMIWNFHCWVESWMTRPDLQPGYEGWQALDPT
PQEKSEGTYCCGPVPVRAIKEGDLSTKYDAPFVFAEVNADVVDWIQQDDGSVHKSINRSL
IVGLKISTKSVGRDEREDITHTYKYPEGSSEEREAFTRANHLNKLAEKEETGMAMRIRVG
QSMNMGSDFDVFAHITNNTAEEYVCRLLLCARTVSYNGILGPECGTKYLLNLNLEPFSEK
SVPLCILYEKYRDCLTESNLIKVRALLVEPVINSYLLAERDLYLENPEIKIRILGEPKQK
RKLVAEVSLQNPLPVALEGCTFTVEGAGLTEEQKTVEIPDPVEAGEEVKVRMDLLPLHMG
LHKLVVNFESDKLKAVKGFRNVIIGPA
```

High resolution crystal structures of TG2 in the closed conformation bound to GTP (guanosine triphosphate) and GDP (guanosine diphosphate) are available. For this study, we selected PDB:4pyg due to its better quality. We were able to detect in total 46 unique crosslinks within TG2. On the PDB:4pyg templated structure 8 out of 46 crosslinks violated the distance cutoff. To refine the template structure, we submitted all detected crosslinks as distance restraints of 35 Å for DSS and PDH or 25 Å for DMTMM crosslinks to I-TASSER. Additionally, we submitted only non-compatible crosslinks to see if an alternative conformation of the structure can be obtained. We evaluated the resulting models by measuring the distances between crosslinked residues with Xwalk. No alternative structures could be obtained and all resulting models violated 8 crosslinks out of 46. However, the model refined with only non-compatible crosslinks had a higher confidence score and TM-score (C-score=1.75 and TM-score0.96+/-0.05), compared to the model refined with all crosslinks (C-score=1.46 and TM-score=0.92+/-0.06). Notably, the relative solvent accessibility of that Lys30 on PDB:4pyg did not conform to our solvent accessibility criteria of at least 40% for the backbone or the side chain. However, the relative solvent accessibility of Lys30 side chain increased considerably after refinement using non-compatible crosslinks (48.6% on refined model vs 24.7% on PDB:4pyg). Therefore, we selected the model structurally refined with noncompatible crosslinks for further docking steps.

*Crosslinks within* TG2 *shown as Euclidean distances mapped onto the predicted model (compatible crosslinks – cyan; non-compatible crosslinks - magenta):*

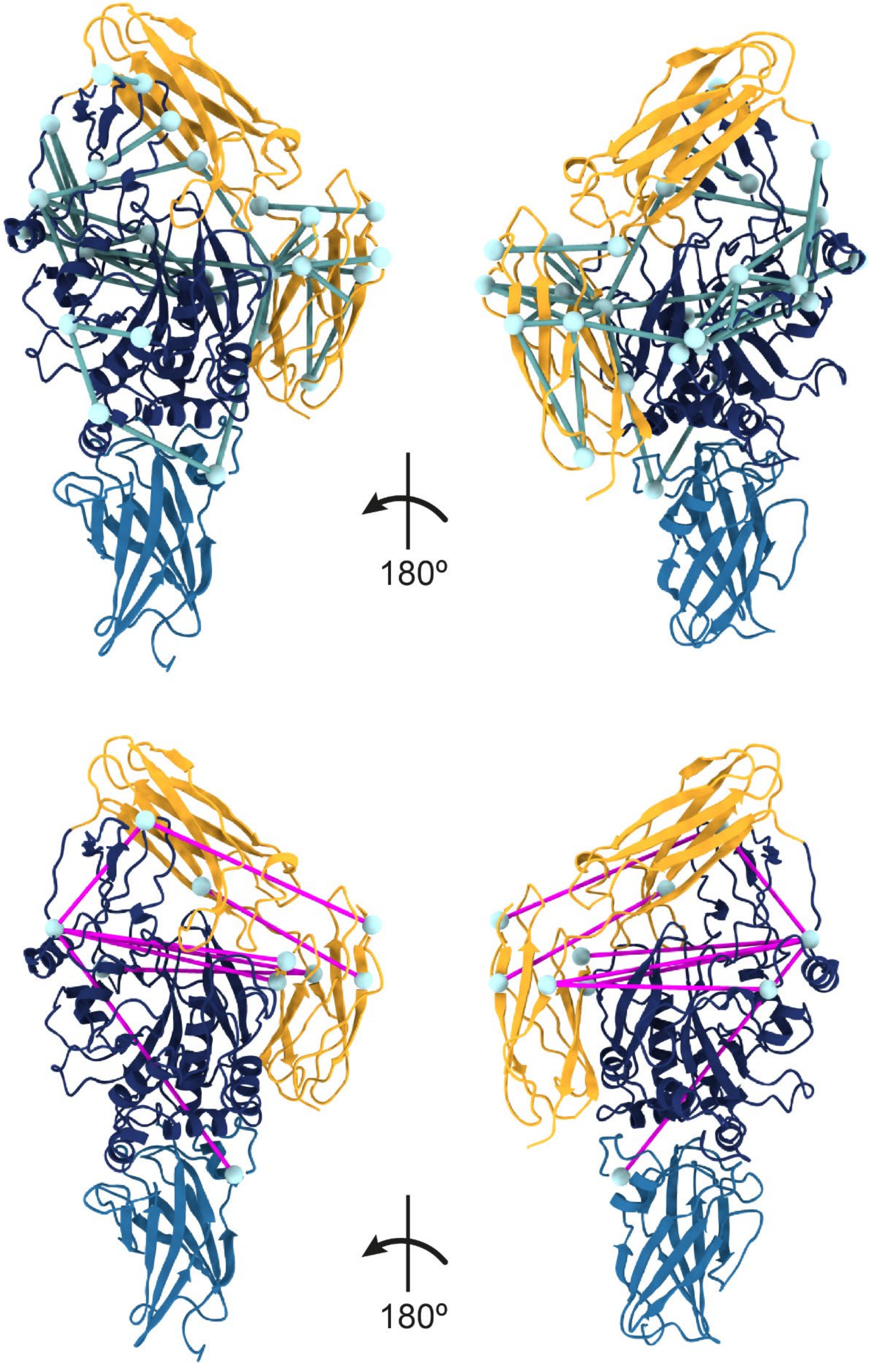

### Crosslinked residues within TG2

| AbsPos1 | AbsPos2 | Link | Crosslinker | CB-CB Euclidean distances (Å) Xwalk |
|---|---|---|---|---|
| 444 | 380 | LYS-LYS | DSS | 6.3 |
| 380 | 464 | LYS-LYS | DSS | 15.5 |
| 649 | 590 | LYS-LYS | DSS | 10.2 |
| 663 | 677 | LYS-LYS | DSS | 11.8 |
| 444 | 464 | LYS-LYS | DSS | 12.5 |
| 562 | 677 | LYS-LYS | DSS | 22.4 |
| 672 | 677 | LYS-LYS | DSS | 16.0 |
| 562 | 464 | LYS-LYS | DSS | 26.0 |
| 444 | 429 | LYS-LYS | DSS | 20.8 |
| 677 | 30 | LYS-LYS | DSS | 34.4 |
| 205 | 30 | LYS-LYS | DSS | 28.8 |
| 464 | 674 | LYS-LYS | DSS | 40.3 |
| 444 | 677 | LYS-LYS | DSS | 30.5 |
| 444 | 425 | LYS-LYS | DSS | 11.7 |
| 425 | 464 | LYS-LYS | DSS | 21.3 |
| 590 | 429 | LYS-LYS | DSS | 22.6 |
| 444 | 590 | LYS-LYS | DSS | 39.1 |
| 444 | 468 | LYS-LYS | DSS | 20.5 |
| 464 | 30 | LYS-LYS | DSS | 53.3 |
| 590 | 464 | LYS-LYS | DSS | 48.1 |
| 590 | 598 | LYS-LYS | DSS | 22.9 |
| 464 | 677 | LYS-LYS | DSS | 38.6 |
| 590 | 677 | LYS-LYS | DSS | 11.7 |
| 429 | 380 | LYS-LYS | DSS | 21.1 |
| 649 | 598 | LYS-LYS | DSS | 25.1 |
| 649 | 562 | LYS-LYS | DSS | 36.3 |
| 444 | 435 | LYS-GLU | DMTMM | 22.4 |
| 444 | 454 | LYS-GLU | DMTMM | 11.7 |
| 640 | 590 | ASP-LYS | DMTMM | 20.7 |
| 366 | 205 | GLU-LYS | DMTMM | 12.9 |

| | | | | |
|---|---|---|---|---|
| 425 | 437 | LYS-GLU | DMTMM | 10.6 |
| 669 | 677 | GLU-LYS | DMTMM | 10.1 |
| 640 | 677 | ASP-LYS | DMTMM | 23.0 |
| 437 | 429 | GLU-LYS | DMTMM | 6.7 |
| 447 | 468 | GLU-LYS | DMTMM | 15.6 |
| 671 | 677 | ASP-LYS | DMTMM | 17.6 |
| 464 | 319 | LYS-GLU | DMTMM | 46.1 |
| 425 | 434 | LYS-ASP | DMTMM | 17.2 |
| 444 | 467 | LYS-GLU | DMTMM | 21.0 |
| 637 | 669 | GLU-GLU | PDH | 10.3 |
| 640 | 669 | ASP-GLU | PDH | 15.2 |
| 643 | 585 | GLU-GLU | PDH | 21.5 |
| 232 | 366 | ASP-GLU | PDH | 10.9 |
| 409 | 319 | ASP-GLU | PDH | 9.8 |
| 643 | 319 | GLU-GLU | PDH | 46.1 |
| 314 | 322 | GLU-GLU | PDH | 15.8 |

**Supplementary Note 2: Analysis of accessible interaction space.**

### Validation of inter-protein crosslinks and prediction of interaction interfaces with DisVis

To evaluate the compatibility of inter-protein crosslinks and to predict active residues at the binding interface, we employed the following workflow. We utilized DisVis, a tool that allows to incorporate experimental crosslinks in form of distance restraints to identify active residues involved in the interaction, simultaneously it can help to filter out incompatible inter-protein crosslinks[13,22]. To aid the identification process, we combined DisVis with the NACCESS[14] tool, which calculates the relative solvent accessibility of each residue. For DisVis interaction analysis, we submitted residues with a relative solvent accessibility of at least 40% for either the main chain or the side chain. As putative active residues, we considered residues with more than 0.5 interaction on average for complexes consistent with the maximum number of restraints. The identified putative active residues were then utilized in the subsequent steps for docking with HADDOCK.

**Solvent accessible residues with at least 40% relative solvent accessibility for either the main chain or the side chain calculated with NACCESS, which were submitted to DisVis interaction analysis:**

**Fixed chain (TG2):** 1 2 3 6 7 8 9 10 11 13 15 16 19 20 22 24 25 27 28 29 30 38 40 42 46 47 48 49 51 52 53 54 56 65 66 67 69 70 72 76 78 80 81 82 84 85 86 87 92 93 94 95 96 97 105 107 109 114 116 122 124 125 126 127 128 129 130 131 132 134 142 144 145 151 152 153 154 157 172 173 175 177 187 191 195 201 202 204 205 206 207 209 213 231 232 233 238 242 243 244 246 247 248 249 263 265 266 267 268 269 270 273 306 308 309 310 312 320 322 324 326 327 329 343 345 346 348 349 350 352 361 362 364 365 366 367 368 369 377 385 387 391 407 408 409 410 411 412 413 419 420 421 425 432 433 435 436 438 441 446 447 448 450 451 453 454 457 458 461 462 464 465 466 467 468 469 470 472 474 475 476 480 481 482 484 487 488 490 498 499 500 501 502 504 506 519 520 522 523 526 528 529 530 531 533 535 536 537 538 539 541 543 545 547 549 550 552 553 555 570 571 572 573 579 580 588 590 592 593 594 595 596 600 601 602 610 612 614 615 618 619 631 632 634 635 637 639 640 641 643 644 645 646 649 651 653 655 657 659 660 661 671 674 677 679 680 681 683 685 686 687

**Scanning chain (FNI$_{2-3}$):** 94 95 96 98 100 101 102 103 104 105 107 108 109 110 111 112 113 115 116 117 118 119 126 127 128 129 130 132 134 135 136 138 139 142 144 145 146 147 149 150 151 153 155 157 159 160 161 162 163 164 170 172 173 174 176 177 178 179 180 181 182 183

**Scanning chain (GBD):** 308 309 311 312 313 315 317 318 319 323 324 325 326 327 328 329 330 336 337 338 339 340 341 342 343 344 346 347 349 350 351 352 353 354 356 358 359 360 361 362 367 368 369 376 377 378 379 380 381 382 383 393 394 395 396 397 402 403 404 405 406 407 409 411 413 414 416 417 422 423 427 428 429 430 432 433 436 437 438 439 440 441 442 450 451 453 454 455 456 465 466 471 473 474 475 479 480 481 482 483 485 486 487 489 490 491 493 495 501 502 503 505 507 508 509 513 514 515 516 519 521 522 523 524 526 527 528 529 530 532 534 535 536 537 538 542 546 548 549 550 552 554 555 556 557 562 564 565 566 567 568 569 571 572 573 574 575 577 579 581 582 583 584 588 590 592 593 594 596 597 598 600 601 602 603 604

**Scanning chain (FNI$_{7-9}$):** 465 466 471 473 474 475 479 480 481 482 483 485 486 487 489 490 491 493 495 501 502 503 505 507 508 509 513 514 515 516 519 521 522 523 524 526 527 528 529 530 532 534 535 536 537 538 542 546 548 549 550 552 554 555 556 557 562 564 565 566 567 568 569 571 572 573 574 575 577 579 581 582 583 584 588 590 592 593 594 596 597 598 600 601 602 603 604

**Scanning chain (FNIII$_{14-15}$):** 1816 1818 1819 1820 1821 1823 1824 1825 1826 1828 1835 1838 1839 1840 1841 1842 1851 1852 1853 1854 1855 1856 1857 1858 1859 1860 1862 1863 1864 1866 1867 1869 1871 1872 1878 1880 1887 1888 1889 1890 1891 1893 1894 1896 1898 1907 1908 1909 1911 1912 1913 1914 1915 1917 1924 1925 1926 1927 1929 1930 1931 1940 1941

1942 1943 1944 1945 1947 1949 1950 1952 1953 1954 1955 1956 1957 1959 1961 1962 1964 1965 1966 1978 1979 1980 1981 1983 1984 1986 1987 1988 1990 1992 1993 1994 1995

**DisVis analysis of TG2 and $FNI_{2-3}$**

We identified three unique crosslinks between TG2 and $FNI_{2-3}$, all of which we subjected to DisVis analysis. For the two DMTMM crosslinks, we set the upper distance limit to 25 Å and 35 Å for the DSS crosslink. However, DisVis analysis failed to identify complexes that were consistent with all three restraints, and highlighted restraint 1 GLU319(TG2)-LYS116(FN) as a false positive candidate with the highest Z-score. In addition, the GLU319(TG2)-LYS116(FN) restraint violated complexes consistent with two restraints 95.92% of the time, while the other two restraints violated the same complexes 4.08% and 0% of the time. After excluding GLU319(TG2)-LYS116(FN) restraint and re-running DisVis analysis, we were able to identify 1346192 complexes consistent with 2 restraints. Given the evidence of GLU319(TG2)-LYS116(FN) incompatibility, we decided to exclude it from further docking steps with HADDOCK.

**Crosslinked residues between TG2 and $FNI_{2-3}$**

| AbsPos FN | AbsPos TG2 | Link | Crosslinker |
|---|---|---|---|
| 116 | 319 | LYS-GLU | DMTMM |
| 116 | 637 | LYS-GLU | DMTMM |
| 100 | 590 | LYS-LYS | DSS |

***Accessible complexes consistent with at least N restraints***

| # of consistent restraints | # of accessible complexes consistent with at least N restraints | Fraction of accessible complexes consistent with at least N restraints |
|---|---|---|
| **0** | 36202322 | 1 |
| **1** | 7018163 | 0.193859 |
| **2** | 1403420 | 0.038766 |
| **3** | 0 | 0 |

***Z-score for each restraint.***

| # of consistent restraints | Average violated fraction | Standard deviation | Z-score | Restraint |
|---|---|---|---|---|
| **1** | 0.861 | 0.098 | 1.329 | A319(CA)-B116(CA) |
| **2** | 0.394 | 0.353 | -0.245 | A637(CA)-B116(CA) |
| **3** | 0.145 | 0.145 | -1.084 | A590(CA)-B100(CA) |

**Active residues output from DisVis interaction analysis with at least 0.5 interactions on average in the complexes satisfying the maximum number of distance restraints:**

**Fixed chain (TG2):** 639, 640, 641, 637, 643, 619, 646, 618, 651, 649, 653, 645, 644, 247, 602, 246, 615, 635, 634, 614, 594, 601, 671, 248, 267, 595, 266

**Scanning chain (FNI$_{2\text{-}3}$):** 103, 117, 100, 101, 102, 116, 104, 118, 145, 130, 105, 98, 115, 129, 144, 119, 146, 94, 132, 96, 128, 95, 107, 134, 112

**DisVis analysis of TG2 and GBD ($FNI_6$-$FNII_1$-$FNII_2$-$FNI_{7\text{-}9}$)**

We were able to identify a total of 8 unique crosslinks between TG2 and GBD, all of which we submitted to analysis with DisVis. Since all crosslinks were on lysines, we set the upper distance limit at 35 Å. Complexes consistent with all 8 restraints could not be found, and as evident by the highest Z-score, DisVis analysis indicated that restraint 5 (LYS550(TG2)-LYS397(FN)) was the most likely false positive. Restraint 8 (LYS30(TG)-LYS486(FN)) had the second highest Z-score, indicating that it too might also be a false positive. However, residue LYS30(TG2) is well-known to be part of the main FN-binding site on TG2 N-terminal domain[23]. The high Z-score for restraint 8 may be due to the asymmetric distribution of detected restraints, where all but one restraint are clustered on the two FN residues (4 on LYS457 and 3 on LYS397), creating a heavy bias. Additionally, restraint 5 violated complexes consistent with the maximum number of restraints 100% and 99.68% of the time for the complexes consistent with 6 restraints, whereas restraint 8 violated those complexes 0% and 94.5% respectively. After excluding the LYS550(TG2)-LYS397(FN) restraint and re-running the DisVis analysis using same parameters, accessible complexes consistent with 7 submitted restraints could be found. Therefore, we excluded restraint LYS550(TG2)-LYS397(FN) from further docking steps with HADDOCK.

***Crosslinked residues between TG2 and GBD***

| AbsPos FN | AbsPos TG2 | Link | Crosslinker |
|---|---|---|---|
| 486 | 30 | LYS-LYS | DSS |
| 457 | 562 | LYS-LYS | DSS |
| 457 | 273 | LYS-LYS | DSS |
| 457 | 364 | LYS-LYS | DSS |
| 397 | 364 | LYS-LYS | DSS |
| 397 | 387 | LYS-LYS | DSS |
| 397 | 550 | LYS-LYS | DSS |
| 457 | 550 | LYS-LYS | DSS |

***Accessible complexes consistent with at least N restraints***

| # of consistent restraints | # of accessible complexes consistent with at least N restraints | Fraction of accessible complexes consistent with at least N restraints |
|---|---|---|
| **0** | 63210314 | 1 |
| **1** | 16953225 | 0.268203 |
| **2** | 6825639 | 0.107983 |
| **3** | 1453687 | 0.022998 |
| **4** | 370523 | 0.005862 |
| **5** | 44879 | 0.00071 |
| **6** | 4847 | 0.000077 |

| | | |
|---|---|---|
| **7** | 12 | 0 |
| **8** | 0 | 0 |

*Z-score for each restraint.*

| # of consistent restraints | Average violated fraction | Standard deviation | Z-score | Restraint |
|---|---|---|---|---|
| **1** | 0.288 | 0.336 | -0.944 | A273(CA)-B457(CA) |
| **2** | 0.44 | 0.297 | -0.184 | A364(CA)-B397(CA) |
| **3** | 0.321 | 0.331 | -0.78 | A364(CA)-B457(CA) |
| **4** | 0.527 | 0.323 | 0.255 | A387(CA)-B397(CA) |
| **5** | 0.823 | 0.123 | 1.737 | A550(CA)-B397(CA) |
| **6** | 0.321 | 0.286 | -0.78 | A550(CA)-B457(CA) |
| **7** | 0.321 | 0.317 | -0.777 | A562(CA)-B457(CA) |
| **8** | 0.771 | 0.317 | 1.474 | A30(CA)-B486(CA) |

**Active residues output from DisVis interaction analysis with at least 0.5 interactions on average in the complexes satisfying the maximum number of distance restraints:**

**Fixed chain (TG2):** 233, 232, 231, 238, 368, 367, 242, 361, 364, 204, 369, 201, 205, 326, 362, 270, 269, 268, 273, 244, 365, 366, 202, 519, 243, 329, 520, 327, 246, 522, 247, 206, 324, 267, 30

**Scanning chain (GBD):** 442, 441, 440, 439, 482, 481, 438, 436, 483, 428, 429, 427, 437, 479, 368, 480, 369, 485, 367, 430, 453, 495, 515, 514

**DisVis analysis of TG2 and $FNI_{7\text{-}9}$**

We detected one unique crosslink between TG2 and $FNI_{7\text{-}9}$, which we submitted to DisVis for interaction analysis. The upper distance limit was set to 35 Å. We then used the active residues obtained for both the fixed and scanning chains for further docking steps with HADDOCK. Notably, DisVis analysis revealed that Glu29 formed at least 0.5 interactions on average in the complexes that satisfied the maximum number of restraints, rather than Lys30, which was crosslinked and is known to be part of the binding interface with FN.

**Crosslinked residues between TG2 and $FNI_{7\text{-}9}$**

| AbsPos FN | AbsPos TG2 | Link | Crosslinker |
|---|---|---|---|
| 486 | 30 | LYS-LYS | DSS |

**Active residues output from DisVis interaction analysis with at least 0.5 interactions on average in the complexes satisfying the maximum number of distance restraints:**

**Fixed chain (TG2):** 687, 1, 8, 9, 686, 96, 2, 600, 97, 268, 601, 29, 47, 46, 266, 15

**Scanning chain ($FNI_{7\text{-}9}$):** 474, 475, 473, 465, 466, 489, 502, 503, 490, 491, 471, 501, 487, 479, 482, 481, 485, 486, 480, 483, 549, 522, 528, 548, 505, 515, 529, 521, 527, 526

**DisVis analysis of TG2 and $FNIII_{14-15}$**

We identified a total of 7 unique crosslinks between TG2 and $FNIII_{14-15}$, all of which we submitted to DisVis for analysis. For the six DSS crosslinks we set the upper distance limit to 35 Å, and for one DMTMM crosslink, we set the upper limit to 25 Å. However, DisVis analysis was unable to identify complexes consistent with all seven restraints. The analysis revealed that restraint 3 (LYS464(TG2)-LYS1862(FN)) had the highest Z-score and was highlighted by DisVis as the candidate for a false positive. This was further supported by the fact that restraint 3 had a 100% and 99.74% violation fractions for the complexes consistent with 6 and 5 restraints respectively. After excluding restraint 3 and re-running the DisVis analysis, we were able to identify 10599 complexes consistent with 6 restraints. As a result, we excluded the LYS464(TG2)-LYS1862(FN) restraint from further docking steps with HADDOCK.

**Crosslinked residues between TG2 and $FNIII_{14-15}$**

| AbsPos FN | AbsPos TG2 | Link | Crosslinker |
|---|---|---|---|
| 1936 | 643 | LYS-GLU | DMTMM |
| 1862 | 677 | LYS-LYS | DSS |
| 1862 | 30 | LYS-LYS | DSS |
| 1862 | 464 | LYS-LYS | DSS |
| 1862 | 590 | LYS-LYS | DSS |
| 1862 | 649 | LYS-LYS | DSS |
| 1837 | 30 | LYS-LYS | DSS |

***Accessible complexes consistent with at least N restraints***

| # of consistent restraints | # of accessible complexes consistent with at least N restraints | Fraction of accessible complexes consistent with at least N restraints |
|---|---|---|
| **0** | 45457055 | 1 |
| **1** | 17186935 | 0.378092 |
| **2** | 7517654 | 0.165379 |
| **3** | 3729043 | 0.082034 |
| **4** | 889924 | 0.019577 |
| **5** | 344514 | 0.007579 |
| **6** | 10599 | 0.000233 |
| **7** | 0 | 0 |

*Z-score for each restraint.*

| # of consistent restraints | Average violated fraction | Standard deviation | Z-score | Restraint |
|---|---|---|---|---|
| **1** | 0.4 | 0.305 | -0.183 | A30(CA)-B1837(CA) |
| **2** | 0.381 | 0.313 | -0.253 | A30(CA)-B1862(CA) |
| **3** | 0.933 | 0.094 | 1.81 | A464(CA)-B1862(CA) |
| **4** | 0.211 | 0.275 | -0.89 | A590(CA)-B1862(CA) |
| **5** | 0.224 | 0.272 | -0.839 | A649(CA)-B1862(CA) |
| **6** | 0.224 | 0.286 | -0.84 | A677(CA)-B1862(CA) |
| **7** | 0.768 | 0.345 | 1.194 | A643(CA)-B1936(CA) |

**Active residues output from DisVis interaction analysis with at least 0.5 interactions on average in the complexes satisfying the maximum number of distance restraints:**

**Fixed chain (TG2):** 639, 640, 637, 653, 602, 641, 634, 619, 601, 635, 655, 600, 643, 266, 631, 651, 618, 632, 267, 615, 247, 644, 646, 263, 268, 246, 645, 657, 614

**Scanning chain ($FNIII_{14\text{-}15}$):** 1864, 1866, 1867, 1824, 1869, 1862, 1823, 1826, 1835, 1871, 1863, 1908, 1872, 1828, 1983, 1986, 1825, 1907, 1984, 1987, 1838, 1821, 1860, 1819, 1842, 1859, 1988, 1909

**Supplementary Note 3: Protein-protein docking.**

**Crosslink-driven HADDOCK protein-protein docking of TG2 with FN fragments**

For all docking runs the maximum number of models considered for clustering was limited to 200.

**TG2 and $FNI_{2\text{-}3}$**

To dock TG2 and $FNI_{2\text{-}3}$ we submitted the two crosslinks as unambiguous distance restraints, which DisVis interaction analysis identified as true positives as well as the active residues that DisVis predicted to be involved in the interaction. The linker between the $FNI_2$ and $FNI_3$ domains (Thr136-Arg140) was defined as a flexible segment. HADDOCK clustered 140 structures in 14 clusters, which represented 70.0% of the water-refined models HADDOCK generated. The distance restraints were mapped onto the best 4 predicted models from the top 10 clusters each (a total of 40 models), and the ED was measured. We found that all predicted models satisfied the distance restraints used for guiding of docking. Based on the HADDOCK scores, we selected the cluster4_1 model, which had the highest score of -210.3+/-11.3.

*Submitted distance restraints:*

```
assi (segid A and resid 637 and name CA) (segid B and resid 116 and name CA) 25 25 0
assi (segid A and resid 590 and name CA) (segid B and resid 100 and name CA) 35 35 0
```

**TG2 and GBD ($FNI_6$-$FNII_1$-$FNII_2$-$FNI_{7\text{-}9}$)**

For docking of TG2 and GBD, we submitted to HADDOCK seven crosslinks as unambiguous distance restraints, which DisVis interaction analysis confirmed to be true positive. Interestingly, DisVis interaction analysis instead of a well-known from the literature residue Lys30, identified a nearby residue Glu29 as an active residue at the binding interface. To the list of the active residues on TG2, we added Lys30, His134, Arg116, which were identified by the mutagenesis studies as the high affinity binding site for FN on TG2[23]. HADDOCK clustered 181 structures in 8 clusters, which represented 90.5% of the water-refined models HADDOCK generated. We mapped crosslinks used for guiding of the docking onto best 4 predicted models in each cluster, which resulted in a total of 32 models. Among those 32 models, only 9 models satisfied the distance cutoff for all 7 crosslinks. Out of those 9 models, 3 models belonged to the top-scoring cluster1 with the highest HADDOCK score of -196.6 +/- 18.6. Therefore, we selected the cluster1_1 model.

*Submitted distance restraints:*

```
assi (segid A and resid 273 and name CA) (segid B and resid 457 and name CA) 35 35 0
assi (segid A and resid 364 and name CA) (segid B and resid 397 and name CA) 35 35 0
assi (segid A and resid 364 and name CA) (segid B and resid 457 and name CA) 35 35 0
assi (segid A and resid 387 and name CA) (segid B and resid 397 and name CA) 35 35 0
assi (segid A and resid 550 and name CA) (segid B and resid 457 and name CA) 35 35 0
assi (segid A and resid 562 and name CA) (segid B and resid 457 and name CA) 35 35 0
assi (segid A and resid  30 and name CA) (segid B and resid 486 and name CA) 35 35 0
```

**TG2 and $FNI_{7\text{-}9}$**

To dock TG2 and $FNI_{7\text{-}9}$, we submitted a single Lys30(TG2)-Lys486(FN) crosslink as an unambiguous distance restraint. As with the docking of TG2 and GBD, we added residues Lys30, His134, Arg116 to the list of active residues directly involved in the interaction. We also defined linkers between the three FN domains (Ala510-Asp515; Pro557-Gln560) as flexible regions. HADDOCK clustered 147 structures in 11 clusters, which represents 73.5% of the water-refined models HADDOCK generated. Since all predicted models satisfied the crosslink, we carefully examined the best structure in each cluster and found that the second highest scoring model (cluster9_1) supported the parallel alignment of TG2 C-terminal β-barrels with $FNI_{8-9}$ domains, while $FNI_7$ was in contact with the high affinity binding site (Lys30, Arg116, His134) on the N-terminal β-sandwich of TG2. Moreover, the standard deviations of the second and first scoring models was also overlapping (top scoring: -200.5 +/- 6.4 and second: -184.1 +/- 19.0).

*Submitted distance restraints:*

```
assi (segid A and resid 30 and name CA) (segid B and resid 486 and name CA) 35 35 0
```

**TG2 and $FNIII_{14-15}$**

To dock TG2 and $FNIII_{14-15}$ we submitted six crosslinks as unambiguous distance restraints, which DisVis identified as true positives, alongside the active residues at the putative binding interface. The linker connecting the $FNIII_{14}$ and $FNIII_{15}$ domains (Ser1900-Asp1904) was defined as a flexible segment. HADDOCK clustered 193 structures in 7 clusters, which represents 96.5 % of the water-refined models HADDOCK generated. We mapped crosslinks on the best 4 structures in each cluster (28 models) and found that all models satisfied all 6 crosslinks within the distance cutoff. Therefore, we selected the model with the highest HADDOCK score, which was the cluster1_2 predicted model.

**Submitted distance restraints:**

```
assi (segid A and resid 30  and name CA) (segid B and resid 1837 and name CA) 35 35 0
assi (segid A and resid 30  and name CA) (segid B and resid 1862 and name CA) 35 35 0
assi (segid A and resid 590 and name CA) (segid B and resid 1862 and name CA) 35 35 0
assi (segid A and resid 649 and name CA) (segid B and resid 1862 and name CA) 35 35 0
assi (segid A and resid 677 and name CA) (segid B and resid 1862 and name CA) 35 35 0
assi (segid A and resid 643 and name CA) (segid B and resid 1936 and name CA) 25 25 0
```

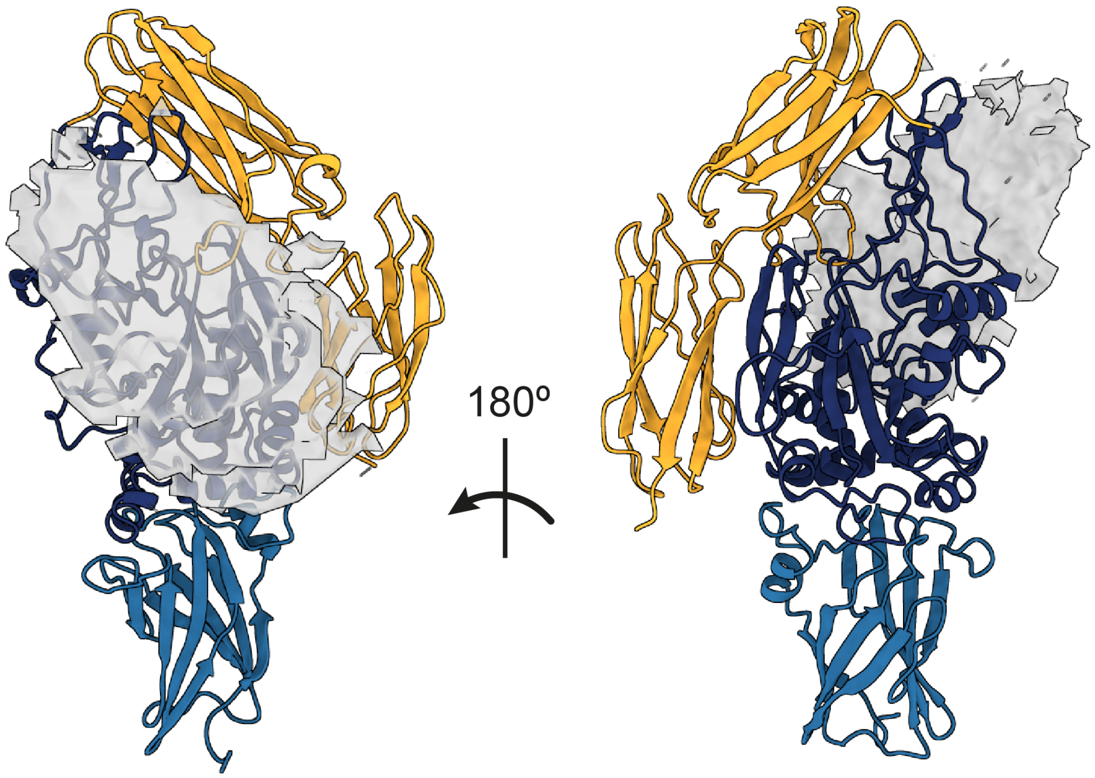


**Supplementary Figure 12: DisVis analysis of the accessible interaction space between TG2 and GBD.** Grey density represents the center of mass of the GBD for complexes consistent 6 restraints. The number of restraints was adjusted for visualization purposes to show interaction interfaces. DisVis analysis was performed using 7 distance restraints.

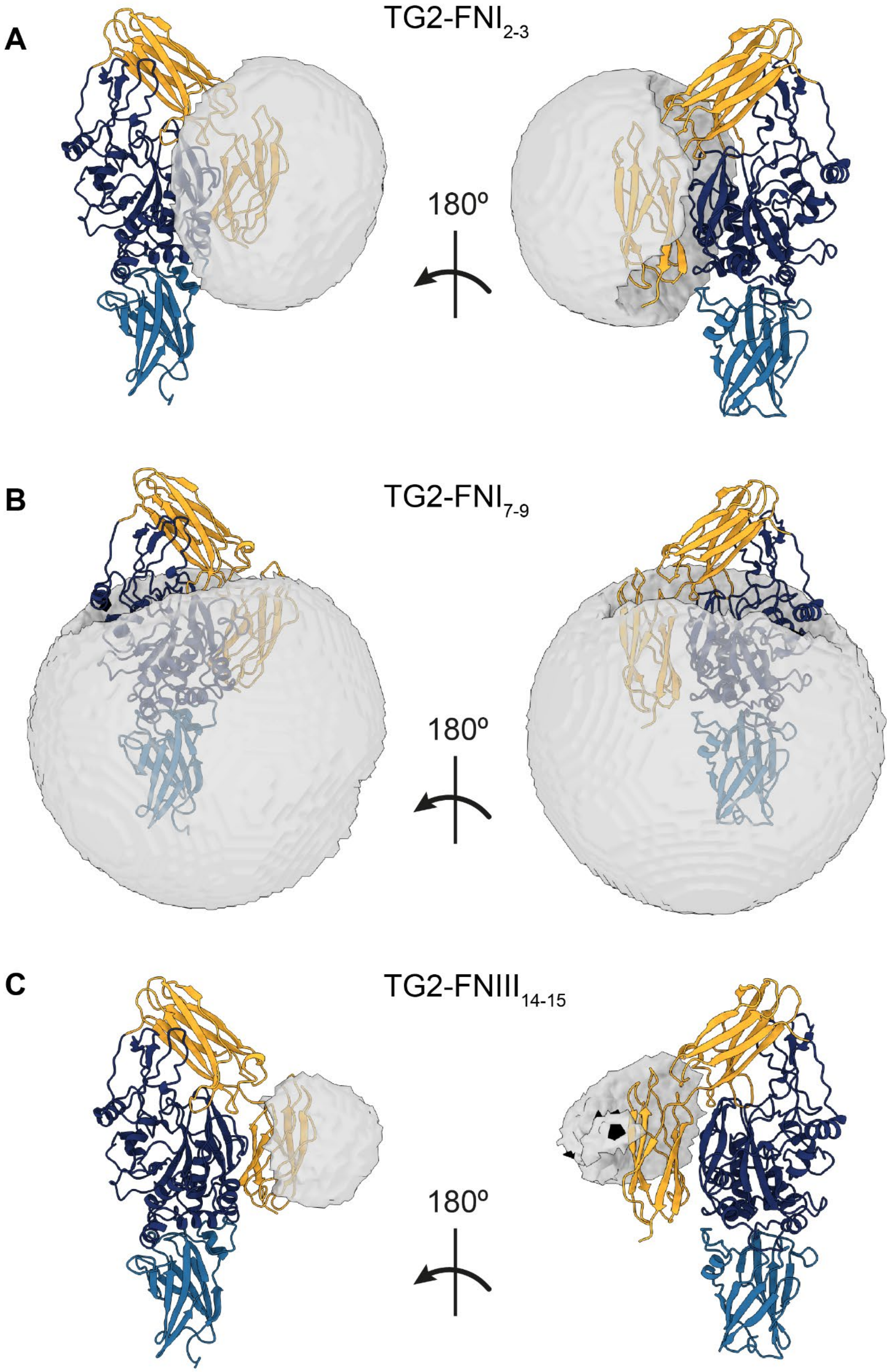
A
TG2-FNI2-3
180º
B
TG2-FNI7-9
180º
C
TG2-FNIII14-15
180º

**Supplementary Figure 13: DisVis analysis of the accessible interaction space between TG2 and FN fragments.** (A) Center of mass of $FNI_{2-3}$ shown in a density map representation for complexes consistent with 2 restraints. (B) Center of mass of $FNI_{7-9}$ shown in a density map representation for complexes consistent with 1 restraint. (C) Center of mass of $FNIII_{14-15}$ shown in a density map representation for complexes consistent with 6 restraints. The number of restraints was adjusted for visualization purposes to show interaction interfaces.

TG2 bound to FN's GBD instead of TG2-GDB

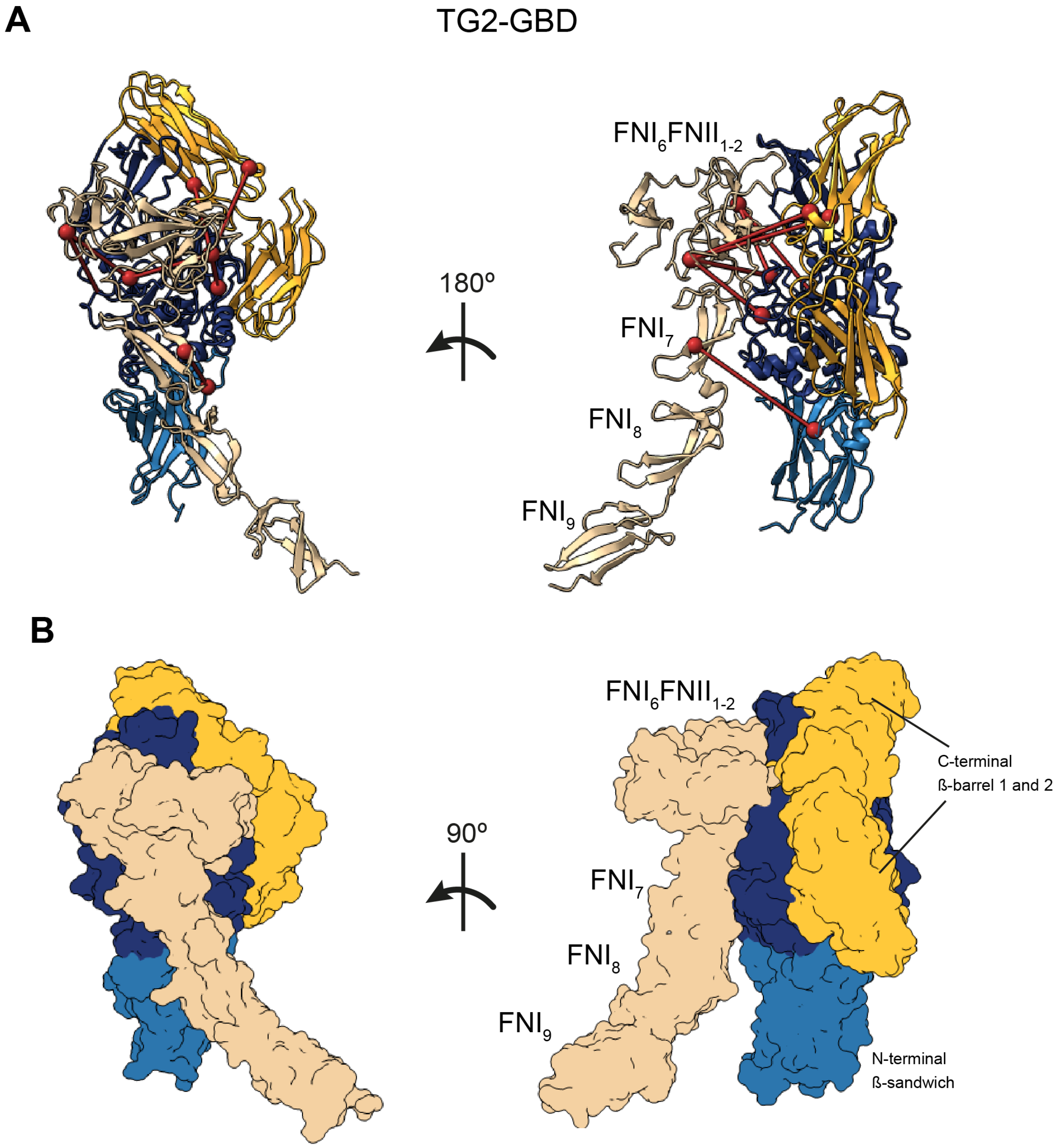


**Supplementary Figure 14: Predicted model of TG2 and GBD complex obtained using crosslink-guided docking with HADDOCK** (A) Ribbon visualization of the TG2 and GBD complex. The seven experimental crosslinks used for guiding of the docking are visualized together with α-carbons. (B) Surface visualization of the TG2 and GBD complex.

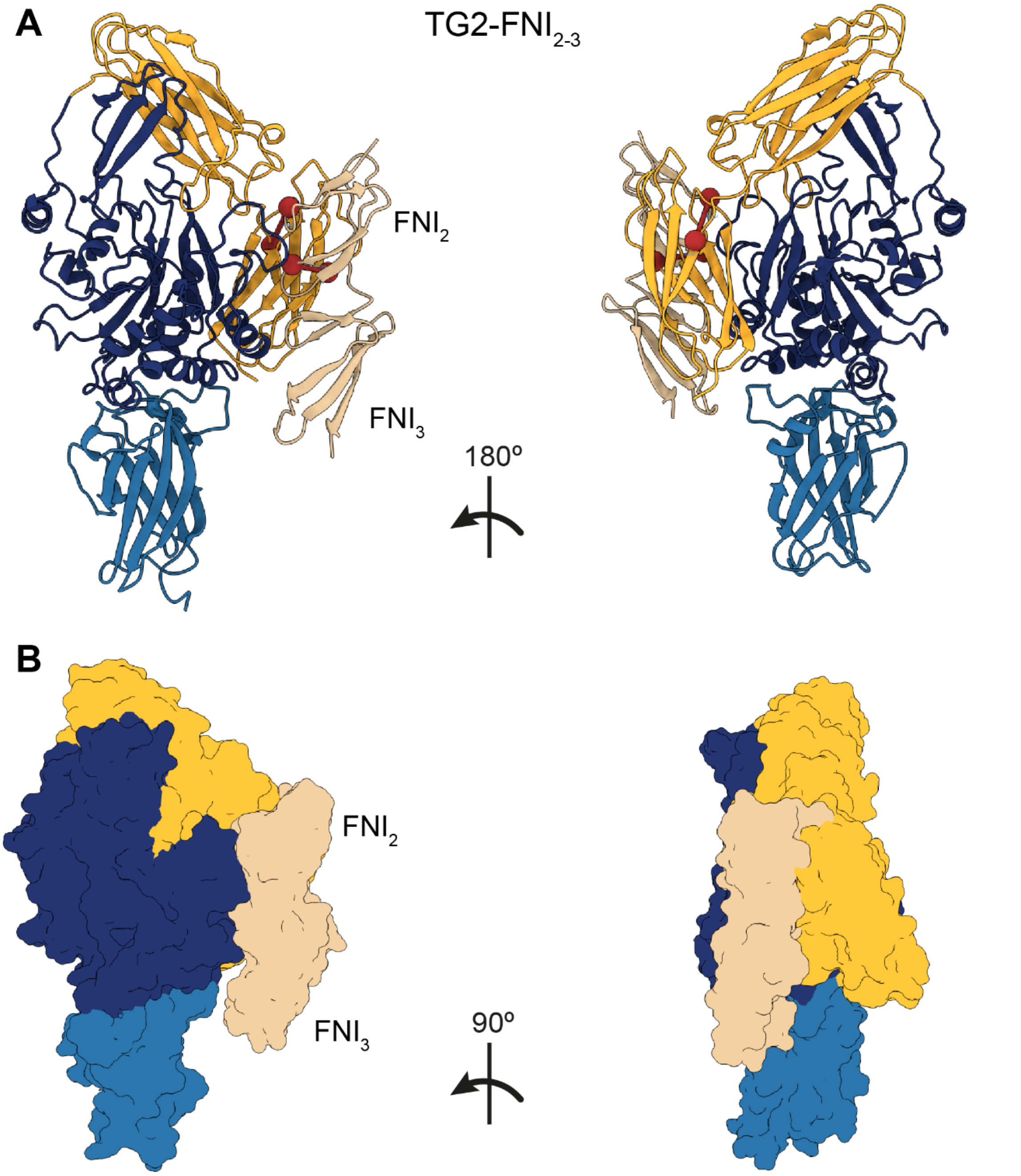


**Supplementary Figure 15: Predicted model of TG2 and FNI$_{2\text{-}3}$ complex obtained using crosslink-guided docking with HADDOCK** (A) Ribbon visualization of the TG2 and FNI$_{2\text{-}3}$ complex. The two experimental crosslinks (637(TG2)-116(FN), 590(TG2)- 100(FN)), used for guiding of the docking are visualized together with α-carbons. (B) Surface visualization of the TG2 and FNI$_{2\text{-}3}$ complex.

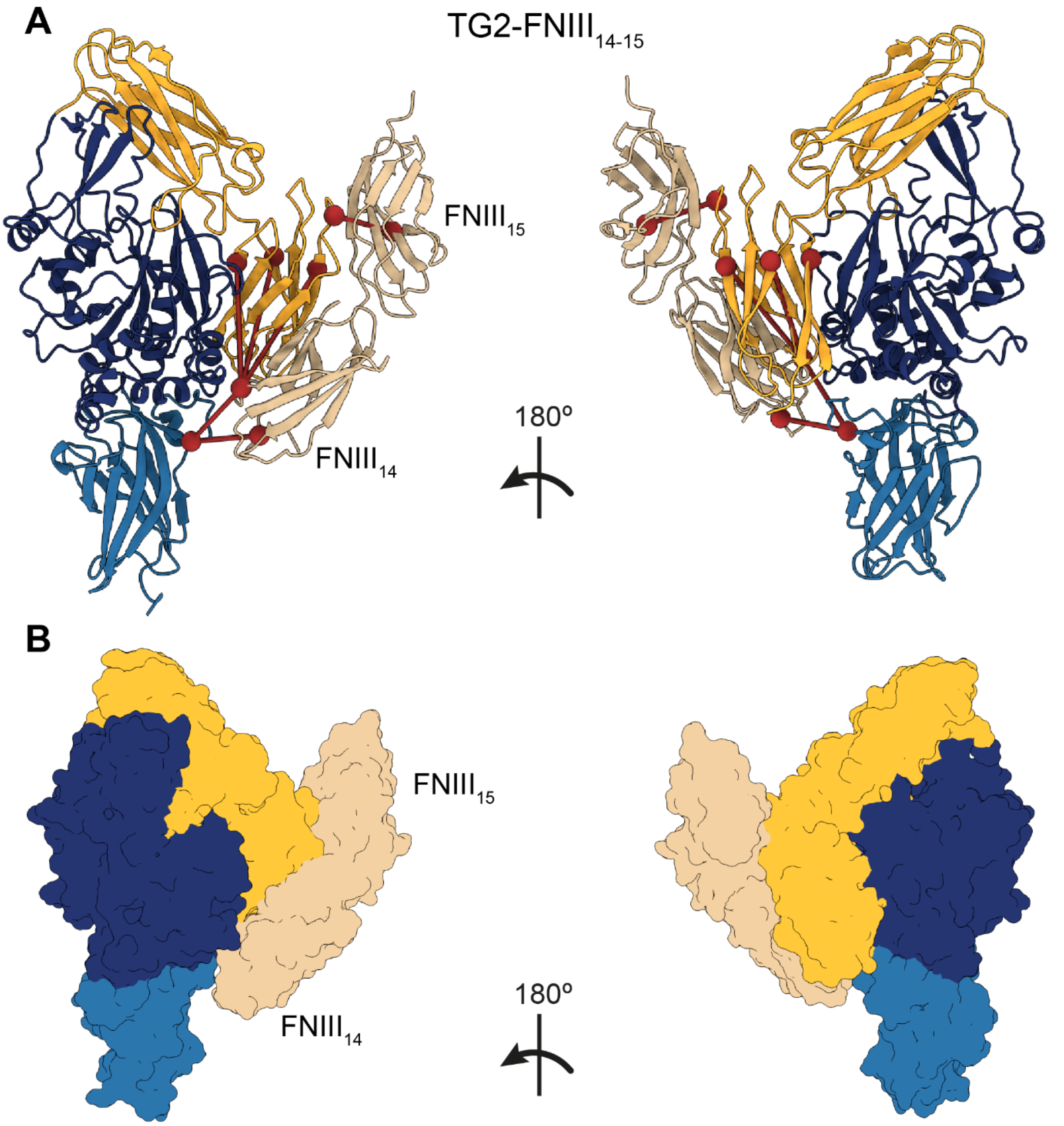


**Supplementary Figure 16: Predicted model of TG2 in complex with $FNIII_{14\text{-}15}$ as obtained using crosslink-guided docking with HADDOCK** (A) Ribbon visualization of the TG2 and $FNIII_{14\text{-}15}$ complex. The six experimental crosslinks used for guiding of the docking are visualized together with α-carbons. (B) Surface visualization of the TG2 and $FNIII_{14\text{-}15}$ complex.

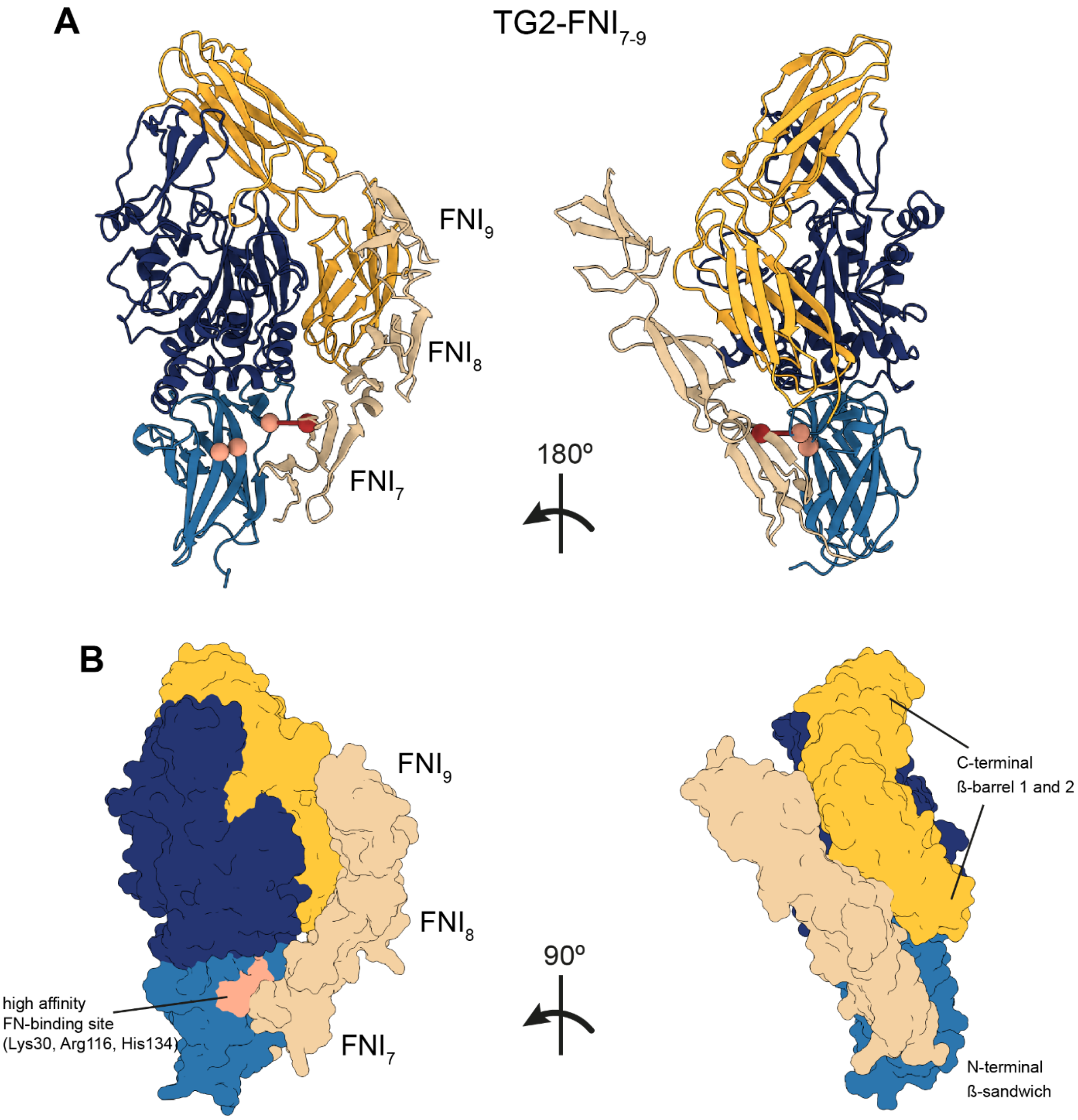


**Supplementary Figure 17: Predicted model of TG2 and FNI$_{7\text{-}9}$ complex obtained using crosslink-guided docking with HADDOCK** (A) Ribbon visualization of the TG2 and FNI$_{7\text{-}9}$ complex. The single experimental crosslink (30(TG2)-486(FN)) used for guiding of the docking is visualized together with the α-carbons. Three residues known to comprise a high affinity binding site on TG2 (Lys30, Arg116, His134) are shown as pink spheres. (B) Surface visualization of the TG2 and FNI$_{7\text{-}9}$ complex. The high affinity FN-binding site on the N-terminal β-sandwich of TG2 (Lys30, Arg116, His134) is coloured in pink. FNI$_{8\text{-}9}$ domains of FN align parallel to C-terminal β-barrel domains of TG2, while FNI$_7$ is in contact with the high affinity binding site on the N-terminal β-sandwich.
